# Generative augmentation recovers prognostic disease signatures from underpowered transcriptomic datasets

**DOI:** 10.64898/2026.04.23.720348

**Authors:** Alok Anand, Manas Pratiti, Syed Yasser Ali, Samanyu Kamra, Shriya Verma, Shrishti Singh, Tanav Bajaj, Tavpritesh Sethi

**Author notes:** Corresponding author: Tavpritesh Sethi.

## Abstract

Identifying reliable gene-expression signatures from small transcriptomic datasets remains challenging. We present GeneLift, a framework that augments underpowered datasets using Gaussian mixture modelling in an analytically invertible latent space, retains genes reproducibly recovered across augmentations, and dates each against the literature using EvidenceClock, a Bayesian PubMed co-occurrence score, as antecedent, subsequent or unreported. Across 52 microarray and RNA-seq datasets spanning breast cancer, ovarian cancer, diabetes, sepsis and tuberculosis, GeneLift recovered candidate genes missed by standard differential expression analysis in all five diseases, including 326 in breast cancer and 976 in ovarian cancer. These candidates carried disease-specific literature support, including genes reported up to 18 years later such as the survival-associated ARL4D, and stratified overall survival in independent cohorts: TCGA-BRCA and METABRIC (log-rank P=6×10^-21^ and 1×10^-42^) and TCGA high-grade serous ovarian cancer (P=1×10^-37^). In benchmarking, Gaussian mixture modelling outperformed more computationally intensive transformer-based generative adversarial networks. Together, GeneLift and EvidenceClock provide an efficient, reproducible framework for recovering prognostically informative signatures from underpowered transcriptomic studies.

---

Reliable disease-specific gene signatures are central to diagnosis, prognosis and risk stratification^1^. High-throughput sequencing has filled public repositories such as GEO, GTEx and single-cell atlases with transcriptomic data^2^, yet most individual studies remain small. This is the central difficulty in signature discovery: the data contain many genes and few samples^3^. In this p >> n setting statistical power is low, so real signals are missed and the signatures that are reported often fail to replicate^4^.

Small-sample and class-imbalance problems have long been addressed with general statistical and machine-learning tools, several of which have been borrowed to synthesise biological data. Bayesian networks, the Synthetic Minority Oversampling Technique (SMOTE) and PCA have all been used to generate synthetic datasets^5,6,7^, and they perform well when the data are low-dimensional. They tend to fail on transcriptomic data, where thousands of genes interact through nonlinear regulation that these low-dimensional models do not capture^8^.

GANs, VAEs and diffusion models can reproduce high-dimensional distributions and generate realistic samples^9,10,11^, and they have been used for privacy-preserving data sharing, dataset augmentation and representation learning^12,13^. Reproducing the data statistically does not mean preserving its biology. They reproduce single-gene distributions but miss the regulatory networks, pathway structure and functional enrichment that downstream discovery requires, and this shortfall is largest for small datasets^14^.

More recent work learns general representations of expression from very large single-cell datasets, from variational models such as scVI to foundation models such as Geneformer, scGPT and scFoundation^15,16,17^. These models need very large training datasets and do not identify individual genes well. A typical disease study has only a few dozen samples and needs specific genes.

Some methods add phenotypic or cross-modal information to improve prediction, but they rely on expert-curated mappings that are not always available^18^. A common gap runs through these approaches: there is no standard way to check whether the augmented data actually carry the biological signal that gene-signature discovery and therapeutic prioritisation need^19^.

A further problem is the space in which augmentation is performed. Generating samples directly in gene-expression space is expensive and overfits when samples are few. Reducing the dimensions first helps, but most common embeddings cannot be inverted exactly, so the synthetic samples cannot be mapped back to individual genes for biological interpretation. Augmentation works best in low dimensions, but interpreting the biology needs gene-level detail. A useful method has to provide both.

Here we present GeneLift, a framework that recovers gene signatures from small transcriptomic studies and weighs each recovered gene against the literature, dating it as antecedent, subsequent or novel. We evaluated GeneLift across five diseases studied with microarray or RNA-seq technologies and benchmarked it against alternative generative models to test whether its recovered genes predict survival in independent clinical cohorts.

## Results

### The GeneLift framework

GeneLift recovers gene signatures from a small dataset in three stages (Fig. 1). A generative model augments the dataset in a PCA-reduced latent space that can be inverted exactly, so every synthetic sample maps back to gene-level values. Genes absent from the original analysis but reproducible across augmentation levels and batches are retained as augmentation-revealed candidates, separating stable signals from incidental ones. EvidenceClock then scores each candidate against the literature and dates it against the first disease-specific report, as antecedent (already known), subsequent (reported only after the source data) or novel (not yet reported). One run therefore returns two kinds of genes: those that recover established disease biology and those with no prior report in the disease. We first compared generative models for their fidelity to the original datasets (Supplementary Table S17).

**Fig. 1.**
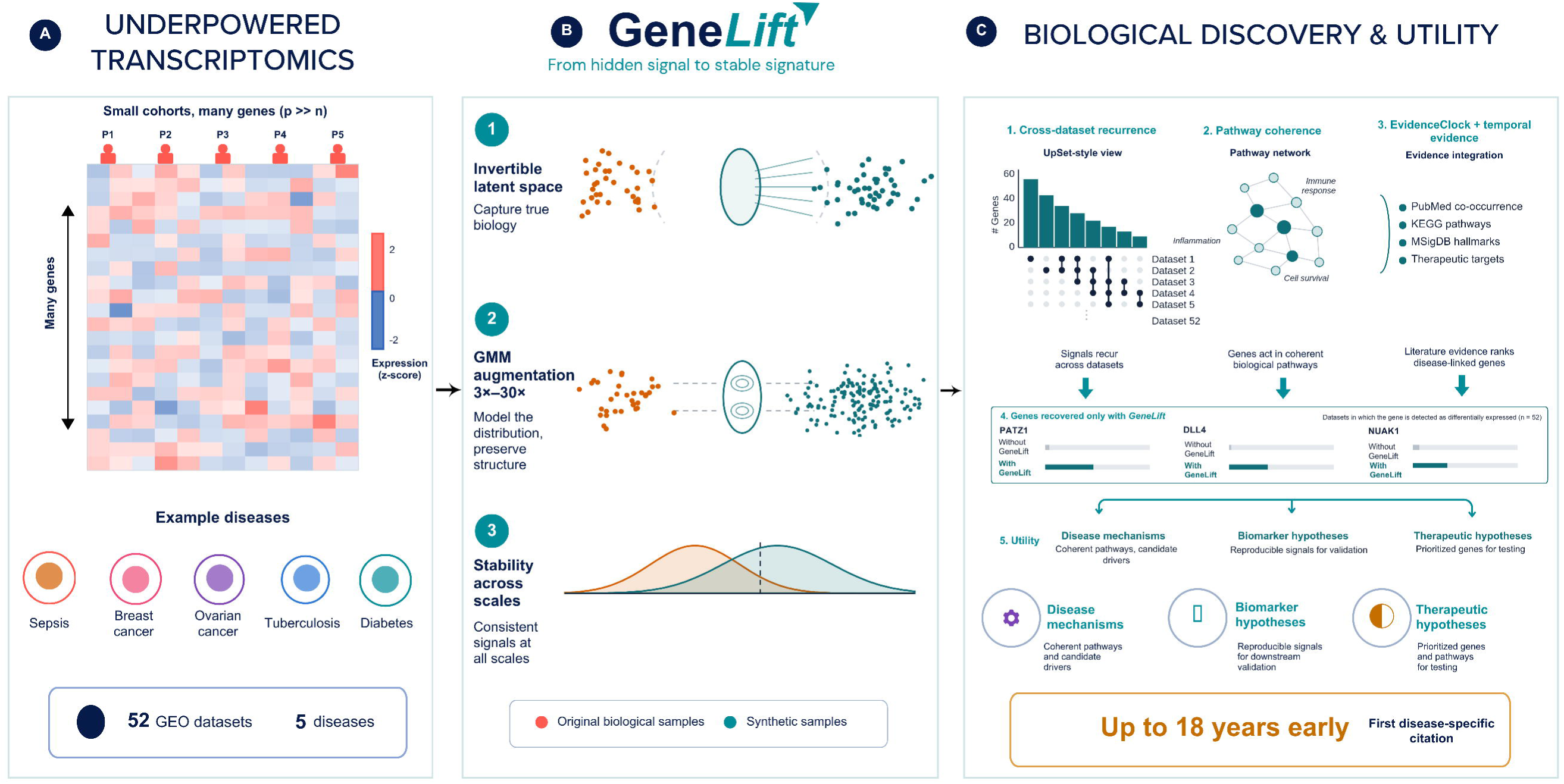
Overview of the GeneLift framework. **A**: Transcriptomic datasets are small, with many genes and few samples (p >> n); GeneLift is tested across 52 GEO datasets spanning five diseases (sepsis, breast cancer, ovarian cancer, tuberculosis and type 2 diabetes) and two technologies Microarray and RNA-seq. **B: Augmentation**. (1) Each dataset is embedded in an analytically invertible latent space so that synthetic samples map back to individual genes; (2) a Gaussian mixture model expands the dataset three-to thirty-fold while preserving its distribution; (3) only genes whose signal is stable across augmentation scales are retained. **C: Discovery and utility**. (1) Augmentation-revealed genes are prioritised by cross-dataset recurrence, (2) pathway coherence, and (3) weighted by EvidenceClock, which integrates PubMed evidence and dates each gene against the literature as antecedent, subsequent or novel; (4) genes obtained post augmentation and absent from original datasets are detected as differentially expressed; (5) generating hypotheses for disease mechanisms, biomarkers and therapeutic targets. In some cases GeneLift flags a gene up to 18 years before its first disease-specific report.

### Benchmarking generative approaches for transcriptomic data augmentation

We benchmarked four generative approaches (Gaussian mixture models, CTGAN, RealTabFormer and Bayesian networks) across 52 datasets and five diseases. The Gaussian mixture model outperformed other models in staying closest to the original data on every fidelity metric: Wasserstein distance, maximum mean discrepancy and a gene-wise t-test.

The gap was clearest when synthetic samples were mapped back to gene space. GMM samples spread across the full original distribution, whereas CTGAN collapsed them into narrow regions and RealTabFormer left a visibly skewed distribution. This held across all five diseases, identifying the GMM as the best structural match for high-dimensional, small-sample transcriptomic data (Table 1). Table 1 shows the best model based on statistical metrics including Maximum Mean Discrepancy, Wasserstein Distance and t test. The complete raw performance metrics for quantitative analysis are reported in Supplementary Table S17.

**Table 1.** Benchmarking of generative approaches, evaluated over 10-fold bootstrapping. Statistical fidelity was quantified as the number of genes for which mean expression did not differ significantly between the original and synthetic datasets (P > 0.05).

| S.No. | GSE ID | Mean (Wasserstein Distance) | Mean (Maximum Mean Discrepancy) | Statistical Fidelity | Disease Class |
| --- | --- | --- | --- | --- | --- |
| 1 | GSE32646 | GMM | GMM | GMM | Breast Cancer |
| 2 | GSE32531 | GMM | GMM | GMM | Breast Cancer |
| 3 | GSE74667 | GMM | GMM | GMM | Breast Cancer |
| 4 | GSE60785 | GMM | GMM | GMM | Breast Cancer |
| 5 | GSE23640 | GMM | GMM | GMM | Breast Cancer |
| 6 | GSE43358 | GMM | GMM | GMM | Breast Cancer |
| 7 | GSE18728 | RTF | RTF | GMM | Breast Cancer |
| 8 | GSE58135 | GMM | GMM | GMM | Breast Cancer |
| 9 | GSE52194 | GMM | GMM | GMM | Breast Cancer |
| 10 | GSE65986 | GMM | GMM | GMM | Ovarian Cancer |
| 11 | GSE63885 | GMM | GMM | GMM | Ovarian Cancer |
| 12 | GSE57477 | GMM | GMM | GMM | Ovarian Cancer |
| 13 | GSE54388 | GMM | GMM | GMM | Ovarian Cancer |
| 14 | GSE25427 | GMM | GMM | GMM | Ovarian Cancer |
| 15 | GSE14407 | GMM | GMM | GMM | Ovarian Cancer |
| 16 | GSE38666 | GMM | GMM | GMM | Ovarian Cancer |
| 17 | GSE29450 | GMM | GMM | GMM | Ovarian Cancer |
| 18 | GSE235641 | GMM | GMM | GMM | Ovarian Cancer |
| 19 | GSE223426 | GMM | GMM | GMM==BN | Ovarian Cancer |
| 20 | GSE160626 | GMM | GMM | BN | Ovarian Cancer |
| 21 | GSE143897 | GMM | GMM | GMM | Ovarian Cancer |
| 22 | GSE102094 | GMM | GMM | GMM==BN | Ovarian Cancer |
| 23 | GSE152532 | GMM | GMM | GMM | Tuberculosis |
| 24 | GSE139871 | GMM | GMM | GMM | Tuberculosis |
| 25 | GSE98461 | GMM | GMM | GMM | Tuberculosis |
| 26 | GSE69581 | GMM | GMM | GMM | Tuberculosis |
| 27 | GSE54992 | GMM | GMM | GMM | Tuberculosis |
| 28 | GSE42832 | GMM | GMM | GMM | Tuberculosis |
| 29 | GSE42830 | GMM | RTF | RTF | Tuberculosis |
| 30 | GSE42825 | RTF | GMM | GMM | Tuberculosis |
| 31 | GSE39939 | GMM | GMM | GMM | Tuberculosis |
| 32 | GSE39940 | GMM | GMM | GMM | Tuberculosis |
| 33 | GSE193777 | GMM | BN | BN | Tuberculosis |
| 34 | GSE126614 | GMM | GMM | GMM | Tuberculosis |
| 35 | GSE103147 | GMM | GMM | GMM==BN | Tuberculosis |
| 36 | GSE79362 | GMM | GMM | GMM | Tuberculosis |
| 37 | GSE166502 | GMM | GMM | GMM | Diabetes |
| 38 | GSE76895 | GMM | GMM | GMM | Diabetes |
| 39 | GSE19420 | GMM | GMM | GMM | Diabetes |
| 40 | GSE253502 | BN | GMM | GMM | Diabetes |
| 41 | GSE184050 | BN | BN | BN | Diabetes |
| 42 | GSE49753 | GMM | GMM | GMM | Sepsis |
| 43 | GSE9692 | GMM | GMM | GMM | Sepsis |
| 44 | GSE69528 | GMM | GMM | GMM | Sepsis |
| 45 | GSE64457 | GMM | GMM | GMM | Sepsis |
| 46 | GSE28750 | GMM | GMM | GMM | Sepsis |
| 47 | GSE66099 | GMM | GMM | GMM | Sepsis |
| 48 | GSE137340 | GMM | GMM | GMM | Sepsis |
| 49 | GSE196117 | GMM | GMM | GMM==BN | Sepsis |
| 50 | GSE185263 | GMM | BN | GMM==BN | Sepsis |
| 51 | GSE133822 | GMM | BN | BN | Sepsis |
| 52 | GSE122274 | GMM | GMM | GMM==BN | Sepsis |

### Augmentation preserves effect-size structure while improving statistical power

Across every augmentation level from 3-to 30-fold batches, the log-fold-change distribution stayed symmetric around zero and followed the original distribution, showing that the Gaussian mixture model introduced no directional bias. With increasing sample size across batches, an increasing trend in statistical significance was observed (Supplementary Fig. S2).

### Breast cancer

Augmentation across seven microarray and two RNA-seq datasets returned 326 genes that were significant in the augmented data but absent from the original TNBC-versus-non-TNBC contrasts (Tables S7 and S12). These genes converged on estrogen-receptor signalling, consistent with established breast-cancer biology (Table S1). EvidenceClock weighed each against the breast-cancer literature. The combined 326-gene set stratified survival in both TCGA-BRCA (1,092 patients) and METABRIC (1,964 patients), with clear separation of risk groups (log-rank P = 6 × 10^−21^ and 1 × 10^−42^; Fig. 2). Sixteen genes appeared in both cohorts of which three were significant and directionally concordant in both (Table 2).

EvidenceClock classified several genes as subsequent, meaning their breast-cancer literature appeared only after our source data. The clearest example is ARL4D^20^, a small GTPase of the ADP-ribosylation-factor-like family that regulates membrane trafficking. It was first reported in breast cancer only after our source data, and then as a passenger locus without functional characterisation, yet in our analysis it marked longer survival in both TCGA-BRCA and METABRIC (Table 2). Microarray datasets identified the transcription factor BATF2^21^ and the long non-coding RNA LINC00963^22^, whereas RNA-seq datasets identified the PCTAIRE-family kinase CDK17^23^ and the PARP-family enzyme TIPARP^24^. These genes, involved in cell proliferation and DNA regulation, were subsequently reported in breast cancer studies 7.2 to 11.9 years after the release of our source datasets.

The strongest novel gene is SDR9C7^25^, a short-chain dehydrogenase/reductase with no breast-cancer literature. It still marked higher risk in both TCGA-BRCA and METABRIC (Table 2). Notably, this represents a specific candidate for functional testing as it has no prior report or signal in the original analysis, yet is reproducible across two independent cohorts. Four of its closest STRING network neighbours (ABCA12, RBP1, CYP26A1 and STRA6) are each independently associated with ceramide homeostasis or retinoid signalling in breast cancer. Genes classified as novel also included TAF4B, a gonad-enriched subunit of the TFIID transcription complex^26^, which marked longer survival in TCGA-BRCA (HR 0.80) and trended the same way in METABRIC (HR 0.75). Its network neighbours have themselves been linked to breast-cancer progression: PPARGC1A promotes breast-cancer metastasis^27^, and inhibition of TAF1 induces antitumour immunity in triple-negative breast cancer^28^.

GeneLift also recovered established breast-cancer biology. Among its recovered antecedent genes were the androgen receptor AR and the angiogenic ligand VEGFA^29^, both absent from the original contrast; these sit alongside well-characterised breast-cancer drivers such as the CDK4/6-inhibitor target CDK6^30^ and the basal-like factor FOXC1^31^. The genes classified as antecedent carried the strongest survival signal. DLL4, the endothelial Notch ligand and an established angiogenic driver of breast cancer^32^, marked higher risk in both cohorts at a hazard ratio near 1.6. In the triple-negative subgroup the established poor-prognosis gene CDCA7^33^ tracked the same direction. The shared signature stayed prognostic after adjustment for triple-negative status (log-rank P = 1 × 10⁻⁵), so its value is not explained by receptor subtype alone.

**Table 2.** Genes prognostic in breast-cancer cohorts.

| Gene | TCGA HR<br>(95% CI) | METABRIC HR<br>(95% CI) | Direction | EvidenceClock | Role |
| --- | --- | --- | --- | --- | --- |
| DLL4 | 1.62<br>(1.24-2.13) | 1.63 (1.08-2.46) | high-risk | antecedent | Notch angiogenic<br>ligand |
| SDR9C7 | 1.27<br>(1.02-1.57) | 1.57 (1.05-2.35) | high-risk | no prior literature | short-chain<br>dehydrogenase/red<br>uctase |
| ARL4D | 0.78<br>(0.65-0.93) | 0.62 (0.44-0.87) | protective | subsequent | ARF-like GTPase |

### Ovarian cancer

Augmentation across nine microarray and four RNA-seq datasets returned 976 genes that were significant in the augmented data but absent from the original analyses (Tables S6 and S11). These genes converged on mitotic-spindle-checkpoint and nucleotide-metabolism signalling, consistent with the proliferative biology of high-grade serous disease (Table S1). These genes separated overall survival (log-rank P = 1 × 10^−37^; Fig. 2), in the TCGA high-grade serous ovarian cancer cohort (420 patients).

EvidenceClock classified several recovered genes as subsequent, meaning their ovarian-cancer literature postdates our source data. Three of these are metabolic enzymes. PYCR1 catalyses the final, NADPH-consuming step of proline synthesis^34^; SAT1 drives polyamine catabolism, which shapes reactive-oxygen output and ferroptosis sensitivity^35^; and MAT2A supplies the S-adenosylmethionine that links one-carbon metabolism to tumour methylation^36^. All three were recovered before their first ovarian-cancer reports. One gene classified as subsequent also carried survival: HABP2, a secreted serine protease active in coagulation and fibrinolysis, is among the panel’s strongest protective markers (HR 0.53).

Genes classified as novel, with no ovarian report at any date, account for most of the prognostic panel: thirteen of its fifteen significant genes are novel or subsequent (Table 3). The two clearest examples map to the compartments most strongly associated with serous survival. FCRLA^37^, a B-cell marker, tracks longer survival where it is high (HR 0.35), matching the favourable course of immune-infiltrated tumours. The vitamin-K-dependent carboxylase GGCX marks the shortest survival^38^ (HR 1.80), consistent with the procoagulant microenvironment of advanced disease. Neither has been reported in ovarian cancer before. We were also able to recover known biological associations from the same underpowered data, with the bevacizumab target VEGFA^39^, the centromeric histone CENPA behind the chromosomal instability that defines the disease^40^ and the tumour suppressor DAB2^41^, with strongest literature evidence. These established drivers surface alongside the new candidates, showing that GeneLift and EvidenceClock recover known biology.

**Table 3.** Featured prognostic genes in TCGA high-grade serous ovarian cancer.

| Gene | HR (95% CI) | Direction | EvidenceClock | Role |
| --- | --- | --- | --- | --- |
| GGCX | 1.80 (1.26-2.57) | high-risk | novel | Vitamin-K-dependent carboxylase; unfavourable in ovarian cancer (Human Protein Atlas) |
| KLHL8<sup>42</sup> | 1.60 (1.18-2.17) | high-risk | novel | Kelch-family Cullin-3 substrate adaptor |
| AKAP11<sup>43</sup> | 1.50 (1.09-2.07) | high-risk | novel | A-kinase anchoring protein 11 |
| BACE1 | 0.71 (0.58-0.87) | protective | novel | Beta-secretase 1 |
| HABP2<sup>44</sup> | 0.53 (0.39-0.72) | protective | subsequent | Secreted serine protease (hyaluronan-binding protein 2) |
| FCRLA | 0.35 (0.21-0.57) | protective | novel | B-cell/plasma-cell Fc-receptor-like molecule; tracks immune infiltration |

**Fig. 2.**
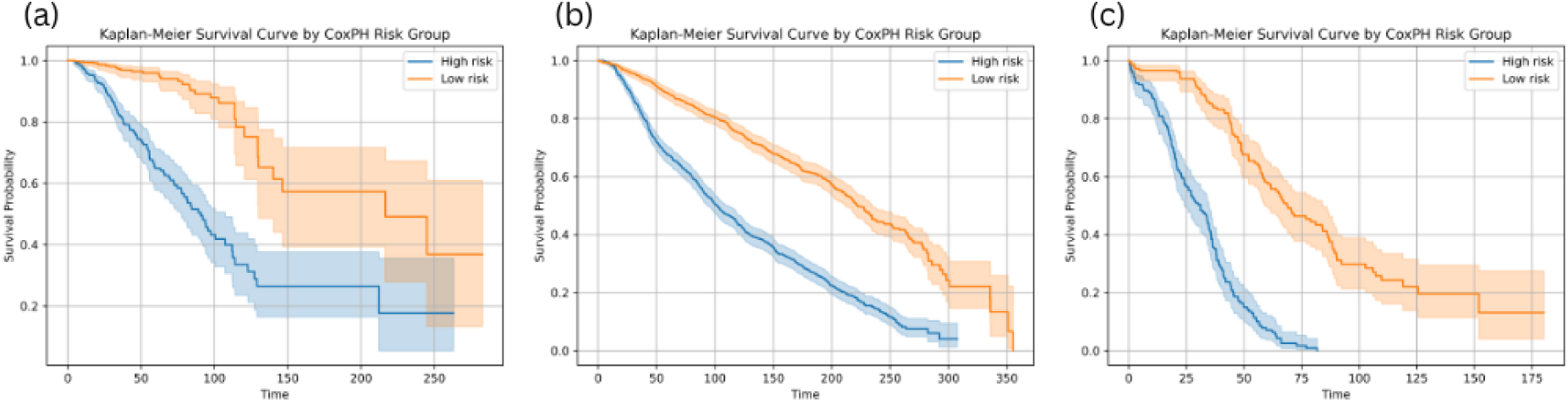
Kaplan-Meier overall survival by GeneLift risk group. In each cohort, the augmentation-revealed candidate genes were fitted in a penalised Cox proportional-hazards model and risk scores for patients (the model linear predictor) were calculated and further stratified into Low and High risk group based on median (a) TCGA-BRCA breast cancer (1,092 patients; log-rank P = 6×10^−21^). (b) METABRIC breast cancer (1,964 patients; log-rank P = 1×10^−42^). (c) TCGA high-grade serous ovarian cancer (420 patients; log-rank P = 1×10^−37^). Curves show survival probability against time (days); shaded bands are 95% confidence intervals.

### Augmentation-revealed genes and 28-day mortality in sepsis

We asked whether augmentation-revealed sepsis genes carry prognostic information in cohorts that had no part in their discovery. In GSE65682 (MARS consortium; 479 sepsis patients, 114 deaths within 28 days), 635 of the 1,200 augmentation-revealed genes were measured and 436 remained after variance and correlation filtering. Age-adjusted Cox regression identified 57 genes associated with 28-day mortality at FDR < 0.05, 17 with hazard ratios above one and 40 below. A direction-weighted score over these genes separated outcome within this cohort (C-index 0.686; log-rank P = 8.3 × 10^−7^; Fig. 3a,b).

The directions of the recovered genes were consistent with established sepsis biology. Genes carrying higher risk were dominated by cell-cycle and granulocyte programmes, including CCNE2, UBE2C and SKA1 alongside the granulocyte activation markers CEACAM6 and CEACAM8, a pattern expected where immature granulocytes are released from the marrow in severe disease. These were accompanied by stress and metabolic genes including DDIT4, MAFG, SLC7A5 and the lactate transporter SLC16A1. Genes associated with longer survival were predominantly related to T-cell and adaptive immunity (TRAF3IP3, HR 0.53; JAK1, 0.51; NLRC5, 0.57; DENND2D, 0.55; ABR, 0.43), consistent with the lymphopenia that accompanies poor outcome in late sepsis.

We then froze the 57 genes and their directions and applied the score, without refitting, to GSE95233 (51 patients with septic shock, profiled on a different array platform). The frozen score separated survivors from non-survivors (AUROC 0.689; Mann–Whitney P = 0.030; 28-day mortality 44% and 23% in the high- and low-scoring halves; Fig. 3c). Performance was essentially unchanged relative to the discovery cohort (C-index 0.686 to AUROC 0.689) despite the change of platform, indicating that the score transfers without the shrinkage expected of an overfitted model.

**Fig. 3.**
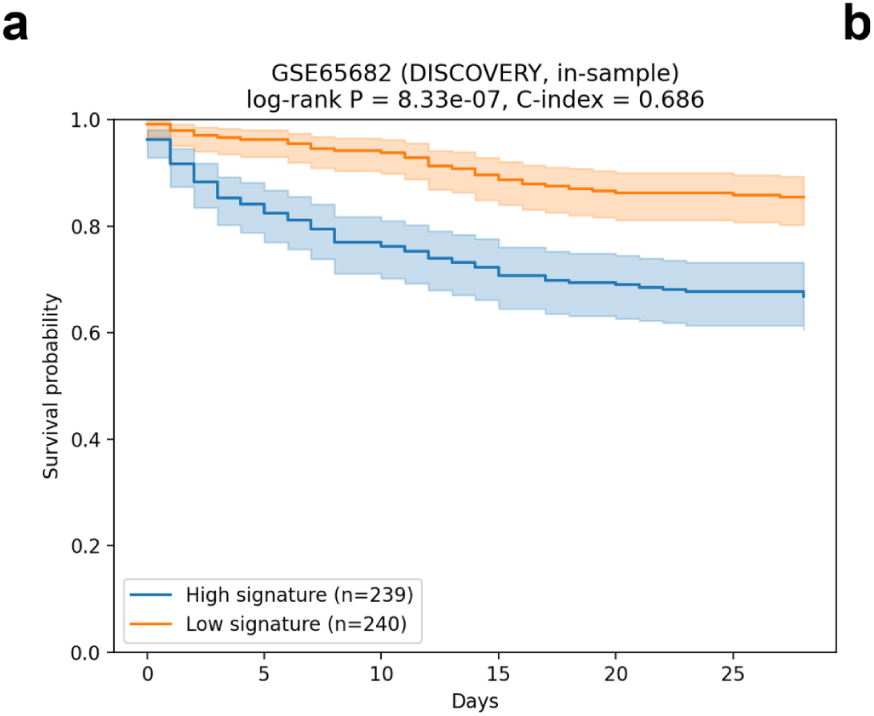
Augmentation-revealed genes and 28-day mortality in sepsis. (a) Kaplan–Meier survival in GSE65682 (479 patients, 114 deaths) by median of the 57-gene signature score; log-rank P = 8.3 × 10^−7^, C-index 0.686, with numbers at risk beneath.

### Generalisation across diseases and technologies, and therapeutic relevance

GeneLift gave similar findings across microarray and RNA-seq technologies. Between 25% and 60% of the pathways found by microarray were also enriched in the RNA-seq data for that disease (Fig. S1; Tables S1, S2, S14, S15); type 2 diabetes showed no such overlap, consistent with comparing whole-blood RNA-seq against islet/adipose microarray (tissue mismatch). Pathway structure and temporal classification both reproduced across the two technologies, indicating that recovery depends on the augmentation procedure rather than on the measurement platform.

To assess the translational relevance of the recovered genes, we cross-referenced them against the Therapeutic Target Database (TTD)^45^. Between 41% and 50% of the reported genes per disease are catalogued TTD drug targets (sepsis 20/40, tuberculosis 15/37, type 2 diabetes 16/38, ovarian cancer 19/39, breast cancer 17/40). Several map to on-target or repurposable agents: in type 2 diabetes GeneLift recovered SLC5A2, the SGLT2 target of the gliflozin class^46^ (bexagliflozin); in tuberculosis IL1B, blocked by canakinumab in IL-1–driven inflammation; in ovarian and breast cancer the bevacizumab target VEGFA, alongside MET (capmatinib, crizotinib) in ovarian cancer and ALK (alectinib, lorlatinib) in breast cancer; and HMGCR, the statin target, in sepsis and ovarian cancer. Recovering approved drug targets from underpowered data lets GeneLift nominate repurposing candidates alongside novel biology.

## Discussion

Despite rapid advances in generative AI, its utility in biology remains constrained by limited biological grounding and inconsistent evaluation frameworks^47^. Here, GeneLift extends beyond gains in statistical power, recovering biologically meaningful gene signals from datasets that are otherwise underpowered for conventional analysis.

Genes classified as antecedent serve as a positive control: from the same underpowered datasets GeneLift recovered established associations, including the gliflozin target SLC5A2 in diabetes, the Mendelian susceptibility gene SP110 in tuberculosis^48^, and the angiogenic Notch ligand DLL4 in breast cancer. Recovering known drivers alongside the novel and subsequent calls is what makes the latter credible.

Because their associations were established independently by the field after the source data were deposited, genes classified as subsequent provide an external test of whether augmentation recovers real biology rather than noise. In sepsis, where the recovered genes converged on T-cell proliferation and TNF/IL-12 control, GeneLift flagged the transcriptional regulator PATZ1^49^ and the deiminase PADI2 years before either was associated with the disease; PADI2 catalyses the protein citrullination required for neutrophil extracellular traps^50^, a mechanism now central to how sepsis is understood. In tuberculosis, where the recovered genes converged on Toll-like and pattern-recognition-receptor signalling, the interferon-induced kinase CMPK2^51^, which supplies nucleotides for mitochondrial DNA and feeds inflammasome signalling, and the glucose transporter SLC2A3^52^, a marker of the metabolic switch in activated macrophages, were both recovered before their first tuberculosis reports. In breast cancer the same class included the PCTAIRE-family kinase CDK17 and the PARP-family enzyme TIPARP, which surfaced from RNA-seq five to seven years before their first breast-cancer reports. In type 2 diabetes, where the recovered genes converged on glutamate secretion and regulation of natural-killer- and T-cell-mediated cytotoxicity, it flagged the AMPK-related stress kinase NUAK1^53^ and the coenzyme-Q biosynthesis factor COQ9^54^, pointing to the mitochondrial-quality-control axis the disease is now understood to involve. The set recovered in ovarian cancer is consistent with evidence for perturbation of proline, polyamine and methylation programmes. Notably, PYCR1 in NADPH-coupled proline synthesis, SAT1 in polyamine turnover and ferroptosis sensitivity, and MAT2A in meeting the methylation demand of the tumour were all found ahead of the recent work validating metabolic dependencies in high-grade serous disease^55^. These candidates are mechanistically coherent with the biology the field later confirmed.

At a broader level, our findings suggest that genes identified as subsequent may represent a less readily discoverable component of disease biology. None of the 95 Subsequent entries (S3–S12) repeated elsewhere in the tables, suggesting that the biological relevance of the Subsequent set was difficult to recognise within the knowledge available at the time. This interpretation remains a hypothesis, however, and will be tested in a future analysis by comparing historical publication density, pathway annotation, disease associations, and network connectivity between antecedent and subsequent genes at the time each dataset was generated.

Genes in the set of novel discoveries have no disease record and are offered on biological grounds, without a citation. Several are plausible from their known molecular function. In tuberculosis, GeneLift uncovered the macrophage-stimulating receptor MST1R (RON)^56^ and the myeloid activating receptor CD300E^57^, both positioned in the host-pathogen interaction. In sepsis it uncovered the pseudokinase TRIB3^58^, which acts through the ATF4-CHOP arm of the endoplasmic-reticulum stress response, consistent with the immunosuppressive phase. In type 2 diabetes it proposed the CoA-biosynthetic enzyme PANK3^59^, which ties the disease to lipid metabolism, and the actin-regulating formin INF2^60^, whose mutations cause focal segmental glomerulosclerosis. In breast cancer the novel short-chain dehydrogenase SDR9C7, uncharacterised in the disease, reproduced as a risk marker across two independent survival cohorts.

We also examined whether recovered genes carry prognostic information at the level of individual patients. In sepsis, a signature drawn from the augmentation-revealed genes transferred between cohorts and across platforms without loss of discrimination, which indicates that the score is not overfitted to its discovery cohort. The composition of the signature is also interpretable. Genes carrying higher risk were dominated by cell-cycle and granulocyte programmes, while those associated with longer survival were largely related to T-cell and adaptive immunity. This division corresponds to the two phases through which severe sepsis is usually described, an early inflammatory response followed by a decline in adaptive immune capacity, and it emerged from the recovered gene set without being sought. In ovarian cancer the vitamin-K-dependent carboxylase GGCX marked the shortest survival, consistent with the procoagulant microenvironment of advanced serous disease, while the B- and plasma-cell marker FCRLA marked the longest, matching the better outcome of immune-infiltrated tumours; the Cullin-3 substrate adaptor KLHL8 and the A-kinase anchoring protein AKAP11 also carried higher risk, extending the novel class into survival analysis. In breast cancer the prognostic genes spanned all three classes, from the subsequent ARL4D through the novel SDR9C7 to the established angiogenic driver DLL4, so genes recovered only after augmentation carried independent prognostic weight and added to the markers already available. Genes classified as subsequent that also carried survival, ARL4D in both breast cohorts and HABP2 in high-grade serous ovarian cancer, carry that external test through to patient outcome.

Whereas recent systems generate hypotheses by reasoning over the published literature^61^, GeneLift derives candidates directly from datasets too small for conventional analysis. It likewise differs from transcriptomic foundation models such as Geneformer and scGPT^15,17^, which are trained on tens of millions of cells; GeneLift is compact and interpretable, and is therefore suited to cohorts of a few dozen samples, where such models cannot be trained and where gene-level attribution is the primary objective.

The framework does have some limitations. It is at present single-omic. There are also counterexamples which are unexplained and need to be explored further. For example, for GATA5, GeneLift recovered a survival-associated signal from expression (HR 1.15, 95% CI 1.03–1.29; Table S16), which is opposite to the expectation from GATA5’s role as a tumour suppressor silenced through promoter hypermethylation^62,63^. Neither result validates the other; resolving it would need GeneLift extended to multi-omic settings covering both expression and methylation.

We release GeneLift and EvidenceClock as an open framework, to further extend its use on diseases and datasets beyond those studied here.

## Online Methods

### Data sources and preprocessing

The framework was applied to five biologically distinct diseases: sepsis, ovarian cancer, breast cancer, tuberculosis, and diabetes (Table 4). Microarray gene expression datasets were obtained from the Gene Expression Omnibus (GEO) and filtered by disease-relevant search terms and platform type to combat platform heterogeneity (GPL570, GPL6480, and GPL10558); RNA-seq counts were sourced from ARCHS4 v2.5^64^ (uniformly kallisto-aligned GEO and SRA counts, GRCh38, Ensembl annotation) to remove upstream-pipeline batch variance, with contrasts matched to the corresponding microarray datasets for the same five diseases. To maintain statistical power and limit noise from underpowered studies, datasets with fewer than 12 samples were excluded. Only datasets with clinically grounded group contrasts were retained.

For microarray data, platform-specific annotation files were used to map probe-level expression values to gene symbols, and expression values were averaged where more than one probe corresponded to the same gene; matrices were then log-transformed and quantile-normalised to make expression distributions comparable across samples. For RNA-seq data, each matrix was normalised with group-aware edgeR filterByExpr, TMM, and limma-voom (quantile)^65,66,67^, yielding a log2 counts-per-million matrix structurally equivalent to the microarray log-quantile matrix.

High-dimensional transcriptomic datasets typically have thousands of features, much of which carry noise. As highlighted above, any augmentation strategy in a reduced-dimensional space relies on a mapping that is analytically invertible to preserve gene-level interpretability. This motivated our use of PCA: as a linear, orthogonal projection, PCA satisfies this requirement with an exact algebraic inverse. In the non-linear alternatives such as t-SNE, UMAP, or autoencoder-based embeddings, back-projection is either undefined or lossy. All generative models were trained on a reduced space of components explaining 95% of the total variance. After data generation, inverse transform was applied to the synthetic samples and they were projected back into the original gene expression space to enable all downstream analyses on gene-level coordinates.

Because Gaussian mixture models had already outperformed the deep generative models on microarray data, we applied the GMM directly to RNA-seq data without modification. The remaining steps were unchanged across both platforms: PCA-latent embedding, 3-to 30-fold augmentation titration, B-statistic stability selection (top 1% in at least four augmented batches, absent from the original differential expression), and EvidenceClock temporal annotation. Every antecedent and subsequent gene-disease citation was verified against the primary PubMed record; acronym collisions and wrong-disease matches were removed, and a genuine earliest paper substituted where one existed.

**Table 4.** Data profiling across the five disease conditions.

| **ID** | **Target Class**<br>**(as specified in**<br>**metadata)** | **Control Class**<br>**(as specified in**<br>**metadata)** | **No. of**<br>**Target**<br>**samples** | **No. of**<br>**Control**<br>**samples** | **Platform** | **Disease** |
| --- | --- | --- | --- | --- | --- | --- |
| GSE137340 | Severe Sepsis | Healthy | 26 | 12 | Microarray | Sepsis |
| GSE28750 | HEALTHY | SEPSIS | 20 | 10 | Microarray | Sepsis |
| GSE49753 | Septic plasma | Uninfected<br>plasma | 24 | 12 | Microarray | Sepsis |
| GSE64457 | Patient | healthy control | 15 | 8 | Microarray | Sepsis |
| GSE66099 | SepticShock | Control | 181 | 47 | Microarray | Sepsis |
| GSE69528 | Septicemic<br>melioidosis | Uninfected<br>healthy | 29 | 28 | Microarray | Sepsis |
| GSE9692 | Septic shock | Controls | 30 | 15 | Microarray | Sepsis |
| GSE65986 | Serous | Endometrioid | 16 | 14 | Microarray | Ovarian<br>cancer |
| GSE63885 | serous ovarian | ca.endometrioid<br>ovarian ca. | 73 | 12 | Microarray | Ovarian<br>cancer |
| GSE57477 | Serous ovarian<br>adenocarcinoma | Serous ovarian<br>borderline<br>tumor | 46 | 6 | Microarray | Ovarian<br>cancer |
| GSE54388 | serous ovarian<br>cancer | healthy<br>(normal) | 16 | 6 | Microarray | Ovarian<br>cancer |
| GSE25427 | serous epithelial<br>ovarian cancer<br>cells | serous<br>borderline<br>ovarian cells | 24 | 4 | Microarray | Ovarian<br>cancer |
| GSE14407 | ovarian<br>adenocarcinoma | Normal | 12 | 12 | Microarray | Ovarian<br>cancer |
| GSE38666 | Ovarian Cancer<br>Epithelium | Ovarian Normal<br>Stroma | 18 | 8 | Microarray | Ovarian<br>cancer |
| GSE29450 | Primary tumors<br>from clear cell<br>ovarian cancer | Normal ovarian<br>surface<br>epithelium | 10 | 19 | Microarray | Ovarian<br>cancer |
| GSE235641 | Ovarian cancer | Benign Tissue | 6 | 6 | Microarray | Ovarian<br>cancer |
| GSE74667 | NTNBC | TNBC | 81 | 14 | Microarray | Breast<br>cancer |
| GSE60785 | NTNBC | TNBC | 42 | 13 | Microarray | Breast<br>cancer |
| GSE43358 | NTNBC | TNBC | 40 | 17 | Microarray | Breast<br>cancer |
| GSE32646 | NTNBC | TNBC | 89 | 26 | Microarray | Breast<br>cancer |
| GSE32531 | NTNBC | TNBC | 16 | 15 | Microarray | Breast<br>cancer |
| GSE23640 | NTNBC | TNBC | 10 | 4 | Microarray | Breast<br>cancer |
| GSE18728 | NTNBC | TNBC | 39 | 22 | Microarray | Breast<br>cancer |
| GSE152532 | active_tb | control | 25 | 11 | Microarray | Tuberculosi<br>s |
| GSE139871 | mycobacterium_t<br>uberculosis_infec<br>tion | healthy_control | 16 | 4 | Microarray | Tuberculosi<br>s |
| GSE98461 | TB patient | healthy control | 4 | 4 | Microarray | Tuberculosi<br>s |
| GSE69581 | active | subclinical | 15 | 10 | Microarray | Tuberculosi<br>s |
| GSE54992 | tuberculosis | healthy_donor | 9 | 6 | Microarray | Tuberculosi<br>s |
| GSE42832 | tb | control | 30 | 30 | Microarray | Tuberculosi<br>s |
| GSE42830 | tb | control | 16 | 38 | Microarray | Tuberculosi<br>s |
| GSE42825 | tb | control | 8 | 23 | Microarray | Tuberculosi<br>s |
| GSE39940 | active_tuberculos<br>is | other_disease | 111 | 169 | Microarray | Tuberculosi<br>s |
| GSE39939 | active_tuberculosis__culture_confirmed_ | other_disease | 35 | 55 | Microarray | Tuberculosis |
| GSE166502 | type_2_diabetes_t2d | normal_glucose_tolerance_ngt | 26 | 26 | Microarray | Diabetes |
| GSE19420 | IGT | NGT | 12 | 12 | Microarray | Diabetes |
| GSE76895 | ND | IGT | 32 | 15 | Microarray | Diabetes |
| GSE122274 | Sepsis | Control | 5 | 10 | RNA-seq | Sepsis |
| GSE133822 | Sepsis | Healthy | 33 | 19 | RNA-seq | Sepsis |
| GSE185263 | Sepsis | Healthy | 348 | 44 | RNA-seq | Sepsis |
| GSE196117 | Sepsis | Control | 33 | 7 | RNA-seq | Sepsis |
| GSE103147 | TB | Control | 41 | 71 | RNA-seq | Tuberculosis |
| GSE126614 | TB | Control | 20 | 19 | RNA-seq | Tuberculosis |
| GSE193777 | TB | Control | 22 | 70 | RNA-seq | Tuberculosis |
| GSE79362 | TB | Control | 110 | 245 | RNA-seq | Tuberculosis |
| GSE184050 | T2D | Control | 25 | 33 | RNA-seq | Diabetes |
| GSE253502 | T2D | Control | 22 | 22 | RNA-seq | Diabetes |
| GSE102094 | Tumor | Normal | 84 | 9 | RNA-seq | Ovarian cancer |
| GSE143897 | Tumor | Normal | 79 | 12 | RNA-seq | Ovarian<br>cancer |
| GSE160626 | Tumor | Normal | 18 | 7 | RNA-seq | Ovarian<br>cancer |
| GSE223426 | Tumor | Normal | 10 | 16 | RNA-seq | Ovarian<br>cancer |
| GSE52194 | TNBC | non-TNBC | 6 | 11 | RNA-seq | Breast<br>cancer |
| GSE58135 | TNBC | non-TNBC | 42 | 42 | RNA-seq | Breast<br>cancer |

### Generative models and data augmentation

To evaluate the suitability of generative models in small-sample transcriptomic studies, four representative generative approaches were evaluated: Conditional Tabular Generative Adversarial Networks (CTGAN)^68^, Bayesian Networks^69^, Gaussian Mixture Models (GMMs), and transformer-based Real TabFormer models^70^. Models were trained independently on the lower dimensional space data and then used to generate synthetic samples having the same dimensionality as the original samples. The performance of models was not solely assessed on the marginal distribution similarity but also on the basis of the preservation of gene-level statistical properties and the stability of downstream biological inference. To facilitate a comparative evaluation, all generative procedures were applied consistently across disease conditions.

### Evaluation of synthetic data fidelity

Several statistical measures were employed to compare the original and synthetic data. The statistical fidelity of the augmented dataset was evaluated by Wasserstein distance and Maximum Mean Discrepancy (MMD) and t test. This quantified the differences in gene-level distribution between original and synthetic data. Two-tailed t-tests were performed gene-wise to check the statistical significance of differences of expression between the two datasets (original vs. synthetic). Genes with a t-test p-value over 0.05 were considered indistinguishable between original and synthetic data as a measure of augmentation fidelity. Further, ten iterations of bootstrap sampling were performed on both original and synthetic data to account for sampling variability. Mean and standard deviation were calculated from the divergence metrics of every bootstrap instance.

### Differential expression analysis and functional enrichment

Before differential expression analysis, to reduce the noise, we removed the genes within the lowest 20% of variance across samples. Limma was used to perform differential expression analysis in R. The high-confidence candidates are the top 1% of the genes in the empirical Bayes B-statistic distribution.

Subsequently, the candidate genes were subjected to pathway and functional enrichment analyses using g:Profiler^71^ against Gene Ontology biological-process terms. The impact of data augmentation on biological signal detection was evaluated by interpreting enrichment results in the context of disease-specific dysregulation, and comparing them between original and augmented datasets.

### Augmentation titration and stability assessment

The effect of augmentation on statistical stability was studied by generating synthetic datasets at increasing levels. The sample size of original data was expanded from threefold to thirtyfold in fixed increments. At each augmentation level, differential expression and enrichment analyses were repeated.

### EvidenceClock

We define G0 as the differentially expressed genes recovered from the original, unaugmented data, and GA as the augmentation-revealed candidates: genes that appeared consistently in at least four augmented batches. This criterion was used to identify signals uncovered through increased sample diversity rather than cohort-specific effects.

To quantify literature support for each gene in GA, we developed EvidenceClock, a literature co-occurrence score derived via Bayes’ rule that treats the biomedical literature as an external evidence base. For a gene G and a target disease D, the score is 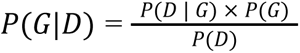 where P(D | G) is the fraction of PubMed records mentioning G that also mention D, P(G) is the prevalence of G across the full PubMed corpus, and P(D) is the prevalence of D in PubMed estimated from a MeSH-plus-Title/Abstract query.

P(G | D) is computed at the time of analysis from live PubMed queries published till March 2026. Genes with zero disease-specific publications at the time of scoring receive P(G | D) = 0, interpreted as an absence of background evidence and a signal of potential novelty. We define T0 as the date each source dataset was published.

Each candidate gene was classified into one of three classes based on the timing of its earliest gene-disease association in the literature relative to T₀. Associations reported before T₀, denoted T−1, were classified antecedent. Associations reported after T₀, denoted T+1, were classified subsequent, with the interval from T₀ to T+1 recorded as the lead time. Candidates with no gene-disease association on record were classified as novel.

### Survival analysis

Augmentation-revealed candidates were tested for prognostic value in clinical cohorts independent of their discovery: TCGA-BRCA and METABRIC for breast cancer, and the TCGA high-grade serous ovarian cancer cohort.

For breast cancer, TCGA-BRCA (cBioPortal^72^) retained 1,092 of 1,097 patients with complete data (115 triple-negative, 977 non-triple-negative) across 20,531 genes, and METABRIC (cBioPortal) retained 1,964 of 1,980 patients (319 triple-negative, 1,645 non-triple-negative) across 20,603 genes. The 326 combined microarray and RNA-seq candidates were mapped to each cohort (199 present in METABRIC) and log2(x+1)-transformed. For ovarian cancer, TPM-normalised RNA-seq from the TCGA high-grade serous cohort provided 40,796 transcript features across 428 samples; Entrez identifiers were mapped to HUGO symbols and duplicate mappings averaged, yielding 40,732 genes with survival data for 420 patients (262 deaths, 158 censored), of which 806 of 976 candidates (nine microarray and four RNA-seq datasets) were present and log2-transformed.

Survival-informative genes were then selected in each cohort by penalised Cox proportional-hazards regression tuned by cross-validated concordance, retaining 51 genes in TCGA-BRCA, 111 in METABRIC, and 48 in the ovarian cohort.

Selected genes were fitted in a multivariable Cox model, yielding per-gene hazard ratios with 95% confidence intervals and Wald P values (22 of 51 significant in TCGA-BRCA, 15 of 48 in ovarian). A per-patient risk score, taken as the model linear predictor, was split at the median, and high-versus low-risk survival was compared by Kaplan-Meier estimation and the log-rank test (TCGA-BRCA P = 6×10^−21^; METABRIC P = 1×10^−42^; triple-negative-adjusted P = 1×10^−5^; TCGA high-grade serous ovarian P = 1×10^−37^).

Statistical analyses used Python 3.7.3 (scikit-learn 1.0.2, scikit-survival 0.17.2, lifelines 0.27.8, numpy 1.21.6, pandas 1.3.5, scipy 1.7.3, statsmodels) and R 4.4.1 (limma 3.62.2, edgeR 4.4.2, bnlearn 5.1.20250224 via rpy2). Functional enrichment used g:Profiler (gprofiler-official 1.0.0); generative benchmarking used CTGAN 0.7.2 and RealTabFormer 0.1.5.

## Data Availability

All datasets analysed in this study are publicly available. The 36 microarray and 16 RNA-seq discovery datasets were obtained from the Gene Expression Omnibus (GEO); their accession numbers are listed in Table 4. Clinical-validation data are also publicly available: TCGA Breast Invasive Carcinoma (TCGA-BRCA) and METABRIC expression and clinical data were obtained from cBioPortal (https://www.cbioportal.org), and TCGA high-grade serous ovarian cancer (TCGA-OV) RNA-seq and survival data from the NCI Genomic Data Commons (https://portal.gdc.cancer.gov). No restrictions apply to the use of these data. Validation cohorts for the sepsis prognostic analysis are available from the Gene Expression Omnibus under accessions GSE65682 and GSE95233.

## Code Availability

The GeneLift framework and all scripts required to reproduce the analyses are available at https://github.com/tavlab-iiitd/GeneLift. The code is released under the MIT License.

## Acknowledgement

The authors gratefully acknowledge funding support from the Centre of Excellence in Healthcare (CoEHE), Indraprastha Institute of Information Technology Delhi (IIIT-Delhi), and from the Department of Biotechnology, Government of India, awarded to Prof. Tavpritesh Sethi. We thank the Department of Computational Biology, Indraprastha Institute of Information Technology Delhi (IIIT-Delhi), for infrastructural support, and the IIIT-Delhi IT Department for their technical assistance. We would also like to extend our gratitude to Ayushi Gupta and Harleen Kaur for their support.

## Author Contributions

TS-(Conceptualization [equal], Methodology [equal], Supervision [lead]); AA-(Conceptualization [equal], Data curation [equal], Software [equal], Visualization [equal], Writing: original draft [equal], Writing: review & editing [equal]); MP-(Conceptualization [equal], Data curation [equal], Software [equal], Visualization [equal], Writing: original draft [equal], Writing: review & editing [equal]); SS-(Data curation [equal], Software [equal]); SYA-(Data curation [equal], Software [equal]); TB-(Software [equal]); SK-(Software [equal]); SV-(Software [equal]).

All authors read and approved the final version of the manuscript and agree to be accountable for all aspects of the work.

## Supplementary Note 1. Full results for sepsis, tuberculosis and type 2 diabetes Sepsis

Sepsis has two immune phases. An early hyperinflammatory phase is followed by an immunosuppressive phase that accounts for most deaths^1^. The second phase is difficult to resolve in small blood-transcriptome datasets. Across seven microarray and four RNA-seq datasets, GeneLift recovered genes associated with αβ T-cell proliferation and with suppression of IL-12 and TNF (Tables S1, S2, S3, S8, S14). These genes were absent from the original analyses. EvidenceClock then dated each against the sepsis literature.

Among the genes classified as subsequent, PATZ1 gave the longest lead time in this study. It was first reported in sepsis eighteen years after GeneLift recovered it. PATZ1 regulates T-lymphocyte differentiation and proliferation^2^, consistent with the immunosuppressive phase. PADI2 was also recovered early. It is a peptidyl-arginine deiminase whose citrullination activity is required for neutrophil extracellular-trap formation^3^. The macrophage-restricted MS4A4A was likewise recovered before its first sepsis report.

Genes classified as novel had no sepsis record at the time of analysis (PubMed, March 2026). SLC1A5, the glutamine transporter ASCT2, is consistent with the amino-acid demand of activated and exhausted immune cells^4^. TRIB3 is a stress-induced pseudokinase acting through the ATF4-CHOP arm of the endoplasmic-reticulum stress response^5^. NLRC5 is a transcriptional regulator of MHC class I^6^ with established innate-immune roles but no prior sepsis-specific study. Genes classified as subsequent included the glycosyltransferase GALNT14 and RGL4, both first reported in sepsis after our source data^7,8^.

Genes classified as antecedent comprised established sepsis immunology. IDO1 is a tryptophan-catabolising enzyme associated with sepsis-induced immune tolerance^9^. SOCS3 is a negative regulator of cytokine signalling. The set also included the established markers STAT4, IL1R2 and the glycolytic enzyme HK2.

### Tuberculosis

Despite being an intracellular pathogen, and notoriously difficult to discover signatures from blood-transcriptome datasets, GeneLift recovered the host signatures of pathogen recognition^10,11^, intracellular defence^12^ and immune-signal control^13,14^ (Tables S1, S2, S4, S9, S14) which were absent from the original analyses.

Three genes were classified as subsequent. CMPK2 is an interferon-induced kinase that supplies nucleotides for mitochondrial DNA^15^ and contributes to inflammasome^16^ and cGAS-STING^17^ sensing of Mycobacterium tuberculosis. It was recovered from RNA-seq before its first tuberculosis report. SLC2A3 (GLUT3) is the high-affinity glucose transporter associated with the metabolic switch of activated macrophages^18^. It was recovered more than a decade before its first tuberculosis report, as was the stress-response gene GADD45B^19^.

Three candidates had no tuberculosis record at the time of analysis (PubMed, March 2026). MST1R (RON) is a macrophage-stimulating receptor that regulates macrophage activation^20^. CD300E is a myeloid activating receptor^21^. Both are consistent with the intracellular site of infection. BCL2L2 is an anti-apoptotic regulator^22^ and is supported less strongly, but could affect the survival of infected cells. None has been tested functionally in tuberculosis.

Genes classified as antecedent comprised established tuberculosis biology. SP110 is a Mendelian susceptibility gene whose loss predisposes to mycobacterial infection^23^. IL1B is a pro-inflammatory cytokine central to the host response^16^. MR1 presents mycobacterial metabolites to MAIT cells^24^. The set also included the chemokine receptors CCR2 and CXCR3 and the NOD2-pathway kinase RIPK2.

### Type 2 diabetes

In blood, type 2 diabetes presents as a diffuse metabolic and inflammatory signal. This signal is difficult to resolve to individual genes in small datasets. Across the type 2 diabetes datasets, GeneLift recovered candidates spanning renal, mitochondrial and immune-metabolic biology (Tables S1, S2, S5, S10, S14). These candidates were absent from the original analyses.

Genes classified as subsequent were recovered from both platforms. The microarray datasets returned the stress kinase NUAK1, first associated with diabetes fifteen years after our source data^25^. They also returned the deubiquitinase USP13, HSPA12A and the synaptic regulator IL1RAPL1. The RNA-seq datasets returned the coenzyme-Q biosynthesis factor COQ9^26^ and the arginine methyltransferase PRMT5^27^. These genes are associated with mitochondrial quality control and metabolic reprogramming.

Two candidates had no diabetes record at the time of analysis (PubMed, March 2026). PANK3 catalyses the first step of coenzyme-A biosynthesis^28^, linking the disease to lipid metabolism, although the pantothenate kinase family has been linked to diabetes since 2015^29^. INF2 was recovered from the type 2 diabetes versus control comparison and PANK3 from the impaired glucose tolerance comparison. INF2 regulates actin filament assembly and disassembly and associates with the cytoplasmic surface of the endoplasmic reticulum. ER-bound INF2 promotes actin-dependent ER-mitochondrial contact, mitochondrial calcium uptake, inner-membrane constriction and mitochondrial division^30^. INF2 also mediates calcium-triggered actin reorganisation, termed calcium-mediated actin reset, which supports acute cellular adaptation to stress^31^ and has a role in focal segmental glomerulosclerosis^32^.

Genes classified as antecedent included established drug targets. SLC5A2 is the sodium-glucose cotransporter SGLT2, targeted by the gliflozin class^33^, and was recovered directly from expression counts. The set also included the GWAS locus TMEM163^34^, the receptor tyrosine kinase PDGFRB, the interferon regulator IRF5 and the glycogen-metabolism subunit PPP1R3B.

## Supplementary Figures

**Fig. S1.**
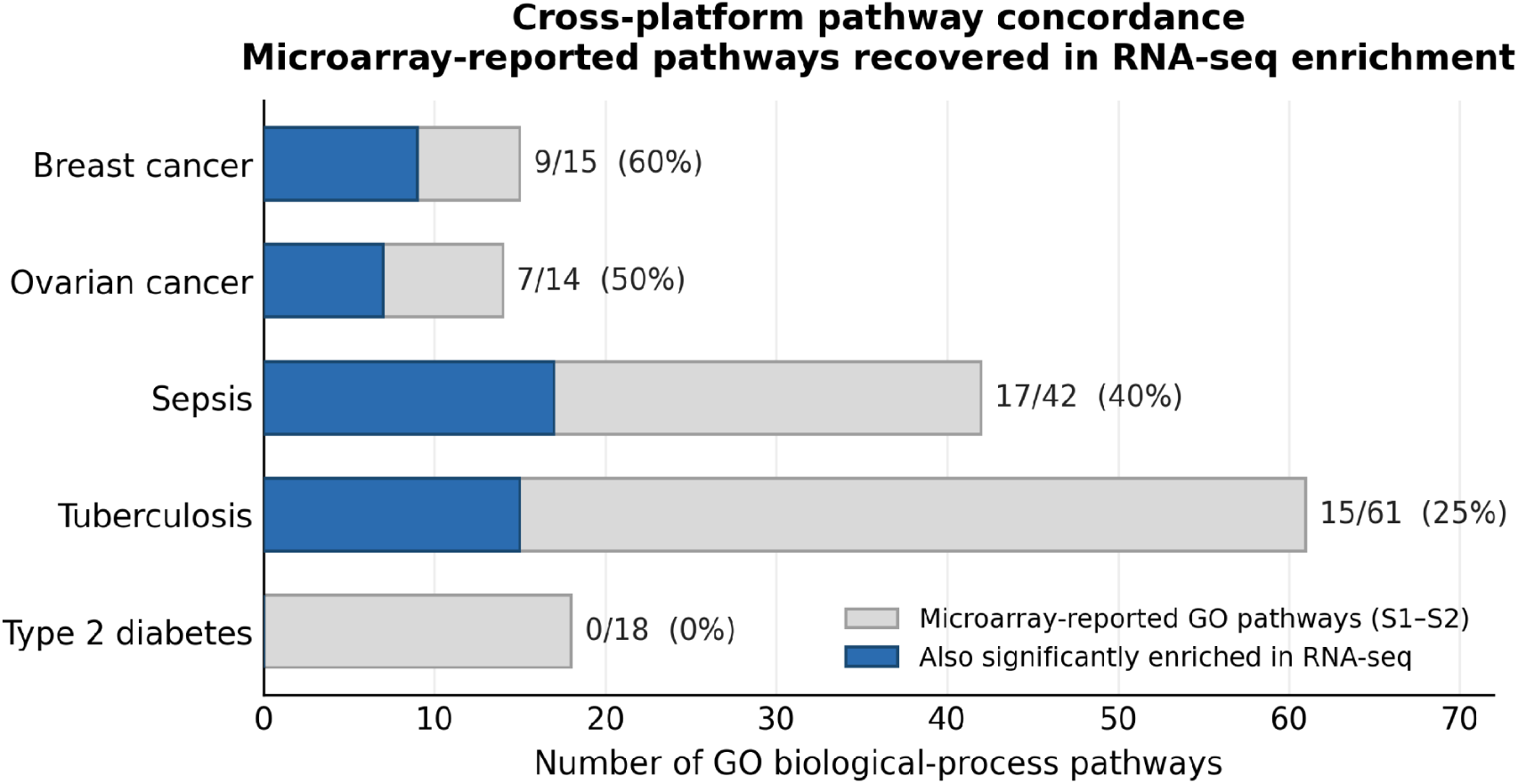
Cross-technology pathway concordance. For each disease, the number of microarray-reported GO biological-process pathways (Tables S1 and S2) that are also significantly enriched (g:Profiler, adjusted p < 0.05) in the augmentation-revealed RNA-seq **gene** sets for the same disease: breast cancer 60%, ovarian cancer 50%, sepsis 40% and tuberculosis 25%. Type 2 diabetes shows no pathway overlap, consistent with the comparison of whole-blood RNA-seq against islet/adipose microarray (tissue mismatch). Recovered themes are biologically coherent, e.g. estrogen-receptor signalling (breast), mitotic-spindle-checkpoint and nucleotide catabolism (ovarian), and Toll-like/pattern-recognition-receptor signalling (sepsis and tuberculosis).

**Fig. S2.**
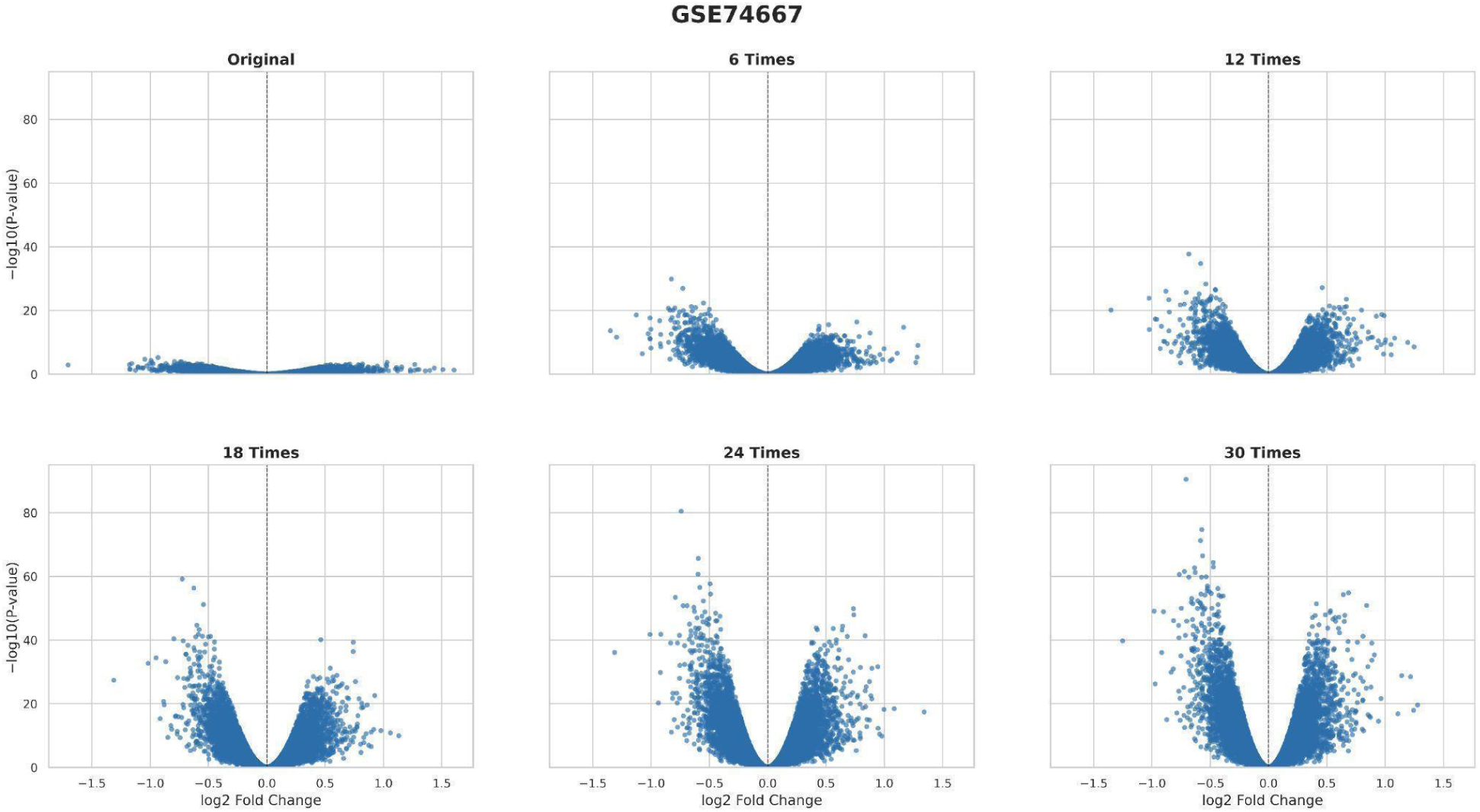
Effect of data augmentation on differential gene expression logFC and adjusted p-values for one of the data GSE74667. All genes with negative log10 of adj.p.val >0 were selected.

### Breast cancer

Fig. S3 Effect of data augmentation on differential gene expression logFC and adjusted p-values across nine breast-cancer datasets (GSE32646, GSE32531, GSE74667, GSE60785, GSE23640, GSE43358, GSE18728, GSE58135 and GSE52194).

### Ovarian cancer

Fig. S4 Effect of data augmentation on differential gene expression logFC and adjusted p-values across thirteen ovarian-cancer datasets (GSE65986, GSE63885, GSE57477, GSE54388, GSE25427, GSE14407, GSE38666, GSE29450, GSE235641, GSE160626, GSE143897, GSE102094 and GSE223426).

### Sepsis

**Fig. S5.**
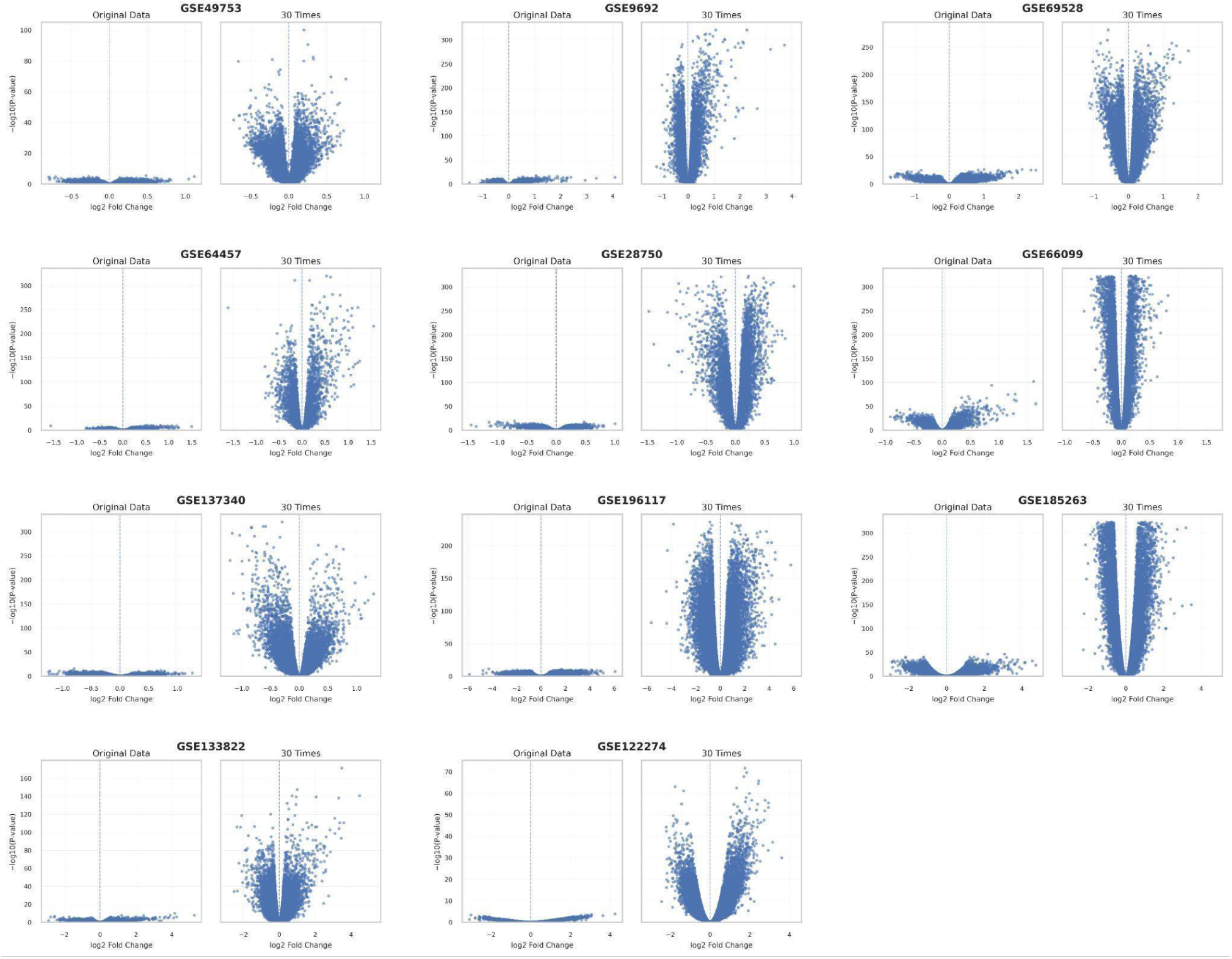
Effect of data augmentation on differential gene expression logFC and adjusted p-values across eleven sepsis datasets (GSE49753, GSE9692, GSE69528, GSE64457, GSE28750, GSE66099, GSE137340, GSE122274, GSE133822, GSE185263 and GSE196117).

### Tuberculosis

**Fig. S6.**
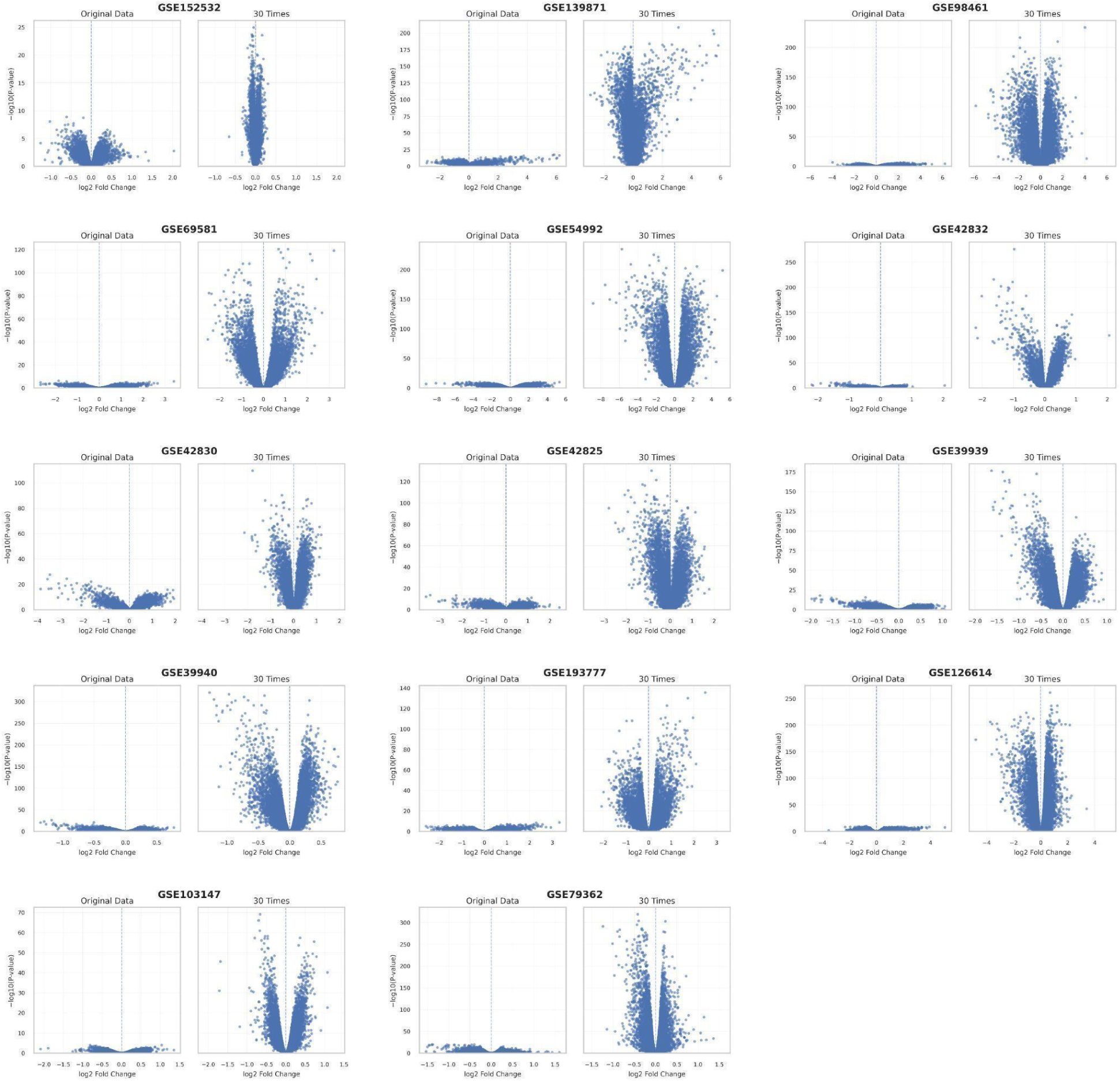
Effect of data augmentation on differential gene expression logFC and adjusted p-values across fourteen tuberculosis datasets (GSE152532, GSE139871, GSE98461, GSE69581, GSE54992, GSE42825, GSE42830, GSE42832, GSE39939, GSE39940, GSE79362, GSE103147, GSE126614 and GSE193777).

### Diabetes

**Fig. S7.**
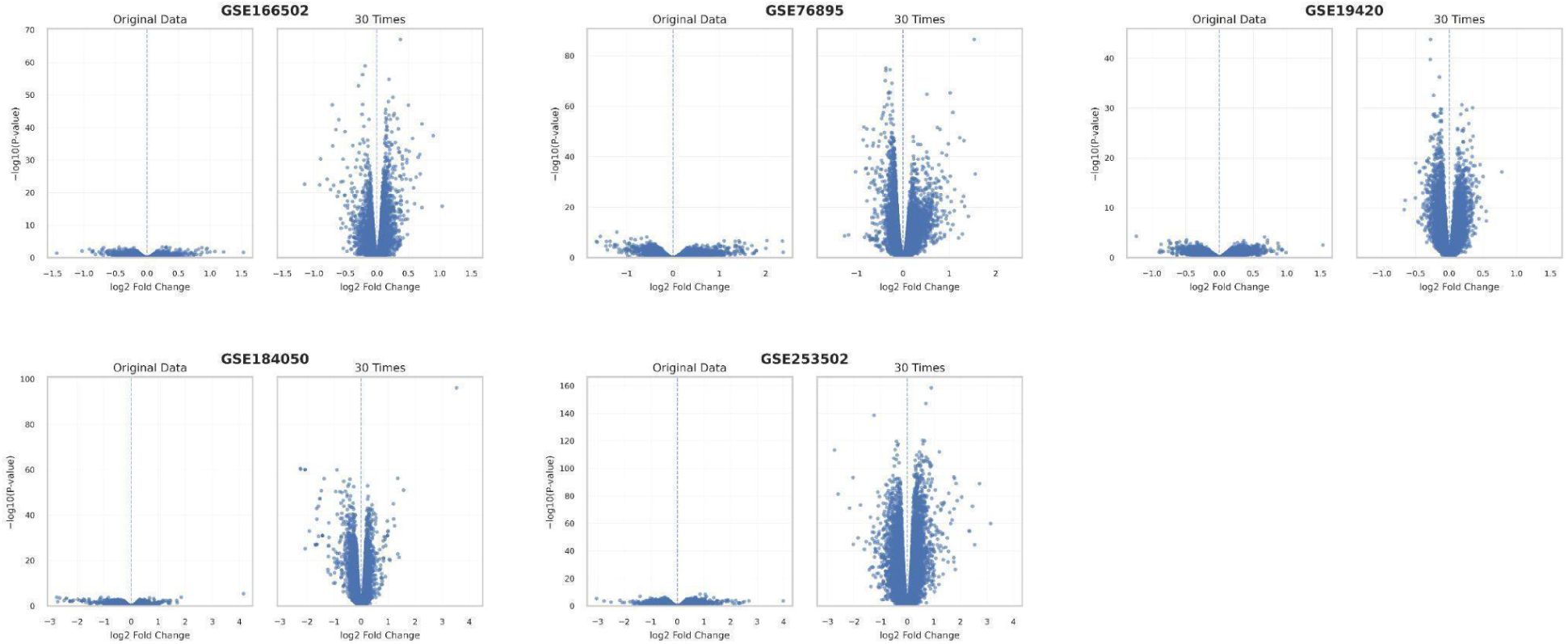
Effect of data augmentation on differential gene expression logFC and adjusted p-values across five type 2 diabetes datasets (GSE166502, GSE76895, GSE19420, GSE184050 and GSE253502).

## Supplementary Tables

**Table S1.** Highly specific biological pathways recurrently observed across two or more datasets of the same disease.

| GO:Term | Biological Role | Relevance to Disease | Disease |
| --- | --- | --- | --- |
| Positive regulation of alpha-beta T cell proliferation | Enhances the expansion of antigen-specific T cells for adaptive immune defence. | Impaired T cell expansion can weaken adaptive immunity during sepsis. | Sepsis |
| Negative regulation of *interleukin-12* production | Suppresses pro-inflammatory cytokine signalling to limit immune overactivation. | Excess suppression may hinder *Th1* responses, worsening infection control in sepsis. | Sepsis |
| Regulation of tumor necrosis factor production | Modulates inflammatory responses and cell survival via *TNF* cytokine control. | Dysregulated *TNF* levels drive systemic inflammation and organ damage in sepsis. | Sepsis |
| Purine deoxyribonucleoside monophosphate catabolic process | Breakdown of purine DNA building blocks | Increased DNA turnover; associated with high proliferation rates | Ovarian cancer |
| Purine deoxyribonucleotide catabolic process | Degradation of purine nucleotides | Dysregulation promotes genomic instability, fuelling tumour progression | Ovarian cancer |
| dAMP catabolic process | Catabolism of deoxyadenosine monophosphate | Affects nucleotide balance, which can impact DNA replication fidelity | Ovarian cancer |
| dGMP catabolic process | Catabolism of deoxyguanosine monophosphate | Imbalance can impair DNA repair and increase mutation burden | Ovarian cancer |
| Mitotic spindle checkpoint signaling | Ensures proper chromosome segregation | Dysregulation leads to aneuploidy, a hallmark of high-grade serous ovarian cancer | Ovarian cancer |
| Negative regulation of microtubule depolymerization | Stabilizes microtubules during mitosis | Disruption linked to paclitaxel resistance, a common chemotherapeutic | Ovarian cancer |
| Spindle assembly checkpoint signaling | Monitors correct spindle formation before division | Defects cause chromosomal missegregation, contributing to tumour heterogeneity | Ovarian cancer |
| Negative regulation of mitotic metaphase/anaphase transition | Delays anaphase onset until chromosomes are aligned | Loss can accelerate cell division and lead to mitotic errors | Ovarian cancer |
| Mitotic spindle assembly checkpoint signaling | Overlaps with mitotic checkpoints ensuring fidelity | Central to preventing chromosomal instability | Ovarian cancer |
| Meiosis I | Division phase in gametogenesis | Reactivation of meiotic genes in somatic tumours is linked to uncontrolled cell division ("meiomitosis") | Ovarian cancer |
| Positive regulation of sodium ion transport | Modulates ion gradients across membranes | Altered ion fluxes can drive proliferation, migration, and cell survival | Ovarian cancer |
| Positive regulation of sprouting angiogenesis | Formation of new blood vessels | Facilitates tumour growth and metastasis; *VEGF* pathway is targeted in ovarian cancer therapy | Ovarian cancer |
| Pyrimidine deoxyribonucleoside metabolic process | Metabolism of pyrimidine nucleosides (e.g., thymidine) | Supports DNA synthesis; upregulated in highly proliferative tumours | Ovarian cancer |
| *NLS*-bearing protein import into nucleus | Nuclear import of proteins with nuclear localization signals | Disruptions can affect localization of tumour suppressors and oncogenes | Ovarian cancer |
| Thymidine metabolic process | Metabolism of thymidine, a DNA precursor | Elevated in rapidly dividing cells; targeted by nucleoside analog chemotherapies (e.g., *5-FU*) | Ovarian cancer |
| CENP-A containing chromatin assembly | Deposition of the centromere-specific histone variant CENP-A to build kinetochore chromatin. | CENP-A overexpression drives chromosomal instability and is an established biomarker of aggressive breast cancer. | Breast Cancer |
| Chromatin remodeling at centromere | Remodels centromeric chromatin structure to maintain kinetochore function during division. | Centromere dysregulation is a recognised route to the aneuploidy seen in breast tumours. | Breast Cancer |
| Estrogen receptor signaling pathway | Nuclear hormone-receptor cascade activated by estrogen binding, driving proliferative gene transcription. | The principal oncogenic driver and first-line therapeutic target in hormone-receptor-positive breast cancer. | Breast Cancer |
| Positive regulation of intracellular estrogen receptor signaling pathway | Enhances estrogen-receptor transcriptional activity within the cell. | Amplified activity sustains proliferation in ER-positive tumours and underlies resistance to endocrine therapy. | Breast Cancer |
| Estrogen response element binding | Direct binding of the estrogen receptor to its DNA response elements to initiate transcription. | A defining molecular event of estrogen-driven gene expression in luminal breast cancer. | Breast Cancer |
| miRNA transcription | Production of microRNA precursors that post-transcriptionally silence target mRNAs. | Dysregulated miRNA transcription reshapes oncogene and tumour-suppressor networks in breast tumours. | Breast Cancer |
| Regulation of miRNA transcription | Controls the rate of microRNA gene transcription. | Altered regulation is linked to subtype-specific miRNA expression signatures in breast cancer. | Breast Cancer |
| miRNA metabolic process | Processing and turnover of microRNAs from primary transcript to mature form. | Disrupted miRNA processing contributes to the aberrant gene expression seen across breast cancer subtypes. | Breast Cancer |
| Mammary gland branching involved in pregnancy | Ductal branching morphogenesis of mammary epithelium during pregnancy. | Reactivation of this developmental programme reflects the shared regulatory circuitry between mammary morphogenesis and tumour growth. | Breast Cancer |
| Protein K63-Linked Ubiquitination | Involves the attachment of ubiquitin chains to proteins, affecting protein function and signalling. | Tags *Mycobacterium tuberculosis* for autophagic clearance; the pathogen evades this to persist in host cells.<sup>1</sup> | Tuberculosis |
| Negative Regulation of MAP Kinase Activity | Regulates the MAPK signalling cascade, controlling immune cell responses to stress, growth, and differentiation. | MAPK signalling (p38, ERK) drives TNF-α, IL-10 and MCP-1 secretion during *M. tuberculosis* infection.<sup>2</sup> | Tuberculosis |
| Negative Regulation of Protein Serine/Threonine Kinase Activity | Controls protein phosphorylation, which regulates cell cycle, apoptosis, and immune signalling. | Regulates mycolic-acid export and cell-envelope remodelling in *M. tuberculosis*, a potential antimycobacterial drug target.<sup>3</sup> | Tuberculosis |
| Cell Surface Pattern Recognition Receptor Signaling Pathway | Detects pathogen-associated molecular patterns (PAMPs) and activates immune responses. | Detects *M. tuberculosis* and activates innate immunity; dysregulation impairs detection and promotes TB persistence.<sup>4</sup> | Tuberculosis |
| Cell Surface Toll-Like Receptor Signaling Pathway | Recognises microbial components like LPS and activates immune signalling. | TLR2 and TLR4 detect *M. tuberculosis*; the bacterium evades TLR signalling to persist.<sup>4</sup> | Tuberculosis |
| Cytoplasmic Pattern Recognition Receptor Signaling Pathway | Detects intracellular pathogens or damage-associated signals. | NOD1, NOD2 and NLRP3 detect cytosolic *M. tuberculosis* and trigger autophagy and inflammasome activation.<sup>4</sup> | Tuberculosis |
| Endolysosomal Toll-Like Receptor Signaling Pathway | Detects intracellular pathogens within phagosomes and triggers immune responses. | TLR2 and TLR4 recognise *M. tuberculosis* proteins, activating NF-κB and cytokine production; a vaccine target.<sup>5</sup> | Tuberculosis |
| Positive Regulation of Interferon-Beta Production | Enhances the production of interferon-beta, important for antiviral and immune responses. | Interferon-beta, induced by RNA sensing of *M. tuberculosis*, modulates the immune response and bacterial replication.<sup>4</sup> | Tuberculosis |
| Positive Regulation of *Tumor Necrosis Factor* Production | Stimulates the production of *TNF*, a cytokine critical for inflammation and immune activation. | Drives *TNF* production that sustains granuloma integrity; *TNF* loss disorganises granulomas and control of infection.<sup>6</sup> | Tuberculosis |
| *Toll-Like Receptor 4* Signaling Pathway | Recognises lipopolysaccharides and activates immune cells to respond to pathogens. | TLR4 recognises *M. tuberculosis* components; the bacterium evades TLR4 signalling, enabling persistent infection.<sup>7</sup> | Tuberculosis |
| Inflammasome-Mediated Signaling Pathway | Detects pathogens and cellular damage, triggering cytokine release (*IL-1β* and *IL-18*) and inflammation. | NLRP3 and AIM2 inflammasomes restrict *M. tuberculosis* via *IL-1β* and pyroptosis; type-I interferon dampens this.<sup>8</sup> | Tuberculosis |
| *Toll-Like Receptor*<br>Signaling Pathway | Detects pathogen-associated molecules and activates immune signalling. | TLR2, TLR4 and TLR9 recognise *M. tuberculosis*; the bacterium manipulates TLR signalling to persist.<sup>4</sup> | Tuberculosis |

**Table S2.** Newly identified Gene Ontology pathways ranked by maximum enrichment depth (top 3).

| Pathway | Depth | Disease |
| --- | --- | --- |
| isotype switching | 12 | Sepsis |
| negative regulation of pentose-phosphate shunt | 11 | Sepsis |
| positive regulation of *type II interferon* production | 10 | Sepsis |
| regulation of pentose-phosphate shunt | 10 | Sepsis |
| somatic recombination of immunoglobulin gene segments | 10 | Sepsis |
| somatic recombination of immunoglobulin genes involved in immune response | 11 | Sepsis |
| *toll-like receptor* signaling pathway | 11 | Sepsis |
| transcription initiation-coupled chromatin remodeling | 10 | Sepsis |
| UMP biosynthetic process | 11 | Sepsis |
| UMP salvage | 12 | Sepsis |
| negative regulation of B cell differentiation | 10 | Sepsis |
| negative regulation of T cell cytokine production | 11 | Sepsis |
| negative regulation of immunoglobulin production | 10 | Sepsis |
| negative regulation of *interleukin-12* production | 10 | Sepsis |
| negative regulation of plasma cell differentiation | 11 | Sepsis |
| pattern recognition receptor signaling pathway | 10 | Sepsis |
| plasma cell differentiation | 10 | Sepsis |
| positive regulation of *CD4*-positive, alpha-beta T-cell activation | 10 | Sepsis |
| positive regulation of *CD4*-positive, alpha-beta T-cell proliferation | 11 | Sepsis |
| positive regulation of *NF-kappaB* transcription factor activity | 11 | Sepsis |
| positive regulation of T cell cytokine production | 11 | Sepsis |
| positive regulation of alpha-beta T cell proliferation | 10 | Sepsis |
| positive regulation of cytokine production involved in immune response | 10 | Sepsis |
| pyrimidine nucleotide salvage | 10 | Sepsis |
| pyrimidine ribonucleoside monophosphate biosynthetic process | 10 | Sepsis |
| pyrimidine ribonucleotide salvage | 11 | Sepsis |
| regulation of *CD4*-positive, alpha-beta T cell proliferation | 10 | Sepsis |
| regulation of T cell cytokine production | 10 | Sepsis |
| regulation of *interleukin-1* beta production | 10 | Sepsis |
| regulation of plasma cell differentiation | 10 | Sepsis |
| regulation of tumor necrosis factor production | 10 | Sepsis |
| *toll-like receptor* signaling pathway | 11 | Sepsis |
| T cell receptor V(D)J recombination | 11 | Sepsis |
| positive regulation of *alpha-beta* T cell proliferation | 10 | Sepsis |
| positive regulation of leukocyte degranulation | 10 | Sepsis |
| somatic recombination of T cell receptor gene segments | 10 | Sepsis |
| D-glucose import | 10 | Sepsis |
| *T-helper 1* cell differentiation | 12 | Sepsis |
| arachidonate binding | 10 | Sepsis |
| negative regulation of chemokine (C-C motif) ligand 5 production | 11 | Sepsis |
| negative regulation of cytokine production involved in immune response | 10 | Sepsis |
| negative regulation of cytokine production involved in inflammatory response | 10 | Sepsis |
| negative regulation of *interferon-alpha* production | 11 | Sepsis |
| negative regulation of *interleukin-12* production | 10 | Sepsis |
| positive regulation of neutrophil degranulation | 11 | Sepsis |
| regulation of *tumor necrosis factor* production | 10 | Sepsis |
| adenylate cyclase-inhibiting G protein-coupled acetylcholine receptor signaling pathway | 8 | Breast cancer |
| epidermal lamellar body | 10 | Breast cancer |
| negative regulation of endoplasmic reticulum unfolded protein response | 8 | Breast cancer |
| negative regulation of transcription by RNA polymerase II | 10 | Breast cancer |
| estrogen receptor signaling pathway | 8 | Breast cancer |
| estrogen response element binding | 10 | Breast cancer |
| ferrous iron transmembrane transporter activity | 8 | Breast cancer |
| mammary gland branching involved in pregnancy | 8 | Breast cancer |
| *miRNA* metabolic process | 8 | Breast cancer |
| *miRNA* transcription | 10 | Breast cancer |
| olfactory learning | 8 | Breast cancer |
| positive regulation of intracellular estrogen receptor signaling pathway | 8 | Breast cancer |
| regulation of *miRNA* transcription | 10 | Breast cancer |
| positive regulation of *miRNA* transcription | 11 | Breast cancer |
| retinal rod cell differentiation | 8 | Breast cancer |
| dAMP catabolic process | 12 | Ovarian cancer |
| dGMP catabolic process | 12 | Ovarian cancer |
| purine deoxyribonucleotide catabolic process | 11 | Ovarian cancer |
| GMP catabolic process | 11 | Ovarian cancer |
| IMP catabolic process | 11 | Ovarian cancer |
| dAMP catabolic process | 12 | Ovarian cancer |
| dGMP catabolic process | 12 | Ovarian cancer |
| negative regulation of tumor necrosis factor production | 11 | Ovarian cancer |
| purine deoxyribonucleotide catabolic process | 11 | Ovarian cancer |
| mitotic spindle assembly checkpoint signaling | 12 | Ovarian cancer |
| negative regulation of mitotic metaphase/anaphase transition | 11 | Ovarian cancer |
| negative regulation of voltage-gated potassium channel activity | 11 | Ovarian cancer |
| positive regulation of ubiquitin protein ligase activity | 12 | Ovarian cancer |
| positive regulation of ubiquitin-protein transferase activity | 11 | Ovarian cancer |
| spindle assembly involved in female meiosis I | 11 | Ovarian cancer |
| de novo' NAD biosynthetic process | 12 | Ovarian cancer |
| de novo' NAD biosynthetic process from tryptophan | 13 | Ovarian cancer |
| positive regulation of *CD4*-positive, alpha-beta T cell proliferation | 11 | Tuberculosis |
| positive regulation of T cell differentiation in thymus | 11 | Tuberculosis |
| protein *K63*-linked ubiquitination | 11 | Tuberculosis |
| GTP biosynthetic process | 11 | Tuberculosis |
| IMP biosynthetic process | 11 | Tuberculosis |
| IMP salvage | 12 | Tuberculosis |
| *L*-alanine import across plasma membrane | 12 | Tuberculosis |
| *L*-alanine transmembrane transport | 11 | Tuberculosis |
| *L*-leucine import across plasma membrane | 11 | Tuberculosis |
| negative regulation of *Arp2/3* complex-mediated actin nucleation | 11 | Tuberculosis |
| negative regulation of *MAP* kinase activity | 12 | Tuberculosis |
| negative regulation of protein serine/threonine kinase activity | 11 | Tuberculosis |
| negative regulation of protein tyrosine kinase activity | 11 | Tuberculosis |
| positive regulation of immature T cell proliferation in thymus | 11 | Tuberculosis |
| purine ribonucleotide salvage | 11 | Tuberculosis |
| DNA damage response, signal transduction by *p53* class mediator<br>resulting in cell cycle arrest | 13 | Tuberculosis |
| *MDA-5* signaling pathway | 12 | Tuberculosis |
| cell surface pattern recognition receptor signaling pathway | 11 | Tuberculosis |
| cell surface toll-like receptor signaling pathway | 12 | Tuberculosis |
| cytoplasmic pattern recognition receptor signaling pathway | 11 | Tuberculosis |
| endolysosomal toll-like receptor signaling pathway | 12 | Tuberculosis |
| negative regulation of *IP-10* production | 11 | Tuberculosis |
| negative regulation of *chemokine (C-X-C motif) ligand 2* production | 11 | Tuberculosis |
| negative regulation of dendritic cell cytokine production | 11 | Tuberculosis |
| positive regulation of cysteine-type endopeptidase activity involved in<br>apoptotic signaling pathway | 12 | Tuberculosis |
| positive regulation of *interferon-beta* production | 11 | Tuberculosis |
| positive regulation of monocyte chemotactic protein-1 production | 11 | Tuberculosis |
| positive regulation of *tumor necrosis factor* production | 11 | Tuberculosis |
| regulation of cysteine-type endopeptidase activity involved in apoptotic signaling pathway | 11 | Tuberculosis |
| *toll-like receptor 4* signaling pathway | 13 | Tuberculosis |
| *MyD88*-dependent *toll-like receptor* signaling pathway | 12 | Tuberculosis |
| T-helper 1 cell differentiation | 12 | Tuberculosis |
| T-helper cell differentiation | 11 | Tuberculosis |
| activation of cysteine-type endopeptidase activity | 11 | Tuberculosis |
| cell surface pattern recognition receptor signaling pathway | 11 | Tuberculosis |
| cell surface *toll-like receptor* signaling pathway | 12 | Tuberculosis |
| cytoplasmic *pattern recognition receptor* signaling pathway | 11 | Tuberculosis |
| inflammasome-mediated signaling pathway | 12 | Tuberculosis |
| positive regulation of *NF-kappaB* transcription factor activity | 11 | Tuberculosis |
| positive regulation of *interferon-beta* production | 11 | Tuberculosis |
| positive regulation of *interleukin-1 beta* production | 11 | Tuberculosis |
| positive regulation of protein tyrosine kinase activity | 11 | Tuberculosis |
| positive regulation of regulatory T cell differentiation | 11 | Tuberculosis |
| *toll-like receptor 4* signaling pathway | 13 | Tuberculosis |
| *toll-like receptor* signaling pathway | 11 | Tuberculosis |
| cell surface *pattern recognition receptor* signaling pathway | 11 | Tuberculosis |
| cell surface *toll-like receptor* signaling pathway | 12 | Tuberculosis |
| cytoplasmic *pattern recognition receptor* signaling pathway | 11 | Tuberculosis |
| endolysosomal *toll-like receptor* signaling pathway | 12 | Tuberculosis |
| inflammasome-mediated signaling pathway | 12 | Tuberculosis |
| positive regulation of *tumor necrosis factor* production | 11 | Tuberculosis |
| *toll-like receptor 4* signaling pathway | 13 | Tuberculosis |
| *toll-like receptor 8* signaling pathway | 13 | Tuberculosis |
| *toll-like receptor 9* signaling pathway | 13 | Tuberculosis |
| *toll-like receptor* signaling pathway | 11 | Tuberculosis |
| *ADP* biosynthetic process | 11 | Tuberculosis |
| positive regulation of eosinophil degranulation | 11 | Tuberculosis |
| *mRNA cis* splicing, *via* spliceosome | 12 | Tuberculosis |
| negative regulation of *MAP* kinase activity | 12 | Tuberculosis |
| negative regulation of *interleukin-1 beta* production | 11 | Tuberculosis |
| negative regulation of protein serine/threonine kinase activity | 11 | Tuberculosis |
| negative regulation of *tumor necrosis factor* production | 11 | Tuberculosis |
| positive regulation of *MAP* kinase activity | 12 | Tuberculosis |
| positive regulation of dendritic cell cytokine production | 11 | Tuberculosis |
| positive regulation of *mRNA cis* splicing, *via* spliceosome | 11 | Tuberculosis |
| positive regulation of myeloid leukocyte cytokine production involved in immune response | 11 | Tuberculosis |
| positive regulation of protein serine/threonine kinase activity | 11 | Tuberculosis |
| positive regulation of *tumor necrosis factor* production | 11 | Tuberculosis |
| positive regulation of *type II* hypersensitivity | 11 | Tuberculosis |
| positive regulation of *type IIa* hypersensitivity | 12 | Tuberculosis |
| positive regulation of tyrosine phosphorylation of STAT protein | 11 | Tuberculosis |
| protein *K63*-linked ubiquitination | 11 | Tuberculosis |
| regulation of *CD4*-positive, *alpha-beta T* cell differentiation | 11 | Tuberculosis |
| regulation of *MAP* kinase activity | 11 | Tuberculosis |
| regulation of *T-helper 1* cell differentiation | 13 | Tuberculosis |
| regulation of *type IIa* hypersensitivity | 11 | Tuberculosis |
| stimulatory *C-type lectin receptor* signaling pathway | 11 | Tuberculosis |
| *toll-like receptor 3* signaling pathway | 13 | Tuberculosis |
| epidermal growth factor receptor activity | 7 | Diabetes |
| glutamate secretion | 8 | Diabetes |
| *COP*II-coated ER to Golgi transport vesicle | 7 | Diabetes |
| CUT catabolic process | 8 | Diabetes |
| CUT metabolic process | 7 | Diabetes |
| negative regulation of natural killer cell mediated cytotoxicity | 8 | Diabetes |
| negative regulation of natural killer cell mediated immunity | 7 | Diabetes |
| phagocytic vesicle | 7 | Diabetes |
| positive regulation of *T cell* activation | 7 | Diabetes |
| positive regulation of *T cell* mediated cytotoxicity | 8 | Diabetes |
| positive regulation of *T cell* mediated immunity | 7 | Diabetes |
| positive regulation of granulocyte differentiation | 9 | Diabetes |
| protection from natural killer cell mediated cytotoxicity | 9 | Diabetes |
| protein localization to centrosome | 8 | Diabetes |
| protein localization to microtubule organizing center | 7 | Diabetes |
| regulation of *T cell* mediated cytotoxicity | 7 | Diabetes |
| regulation of cytokine production | 7 | Diabetes |
| regulation of natural killer cell mediated cytotoxicity | 7 | Diabetes |

**Table S3.** Augmentation-revealed microarray gene signatures, Sepsis.

| Gene | Class | Platform | Dataset (GSE) | $\Delta$ (yr) | Earliest PMID |
| --- | --- | --- | --- | --- | --- |
| ABR | Antecedent | Microarray | GSE49753 | -31.5 | 7180439 |
| DBP | Antecedent | Microarray | GSE9692 | -24.7 | 6687604 |
| AES | Antecedent | Microarray | GSE28750 | -21.8 | 2301634 |
| CD5 | Antecedent | Microarray | GSE28750 | -21.6 | 1693799 |
| IL1R2 | Antecedent | Microarray | GSE28750 | -18.7 | 7680670 |
| TIMP2 | Antecedent | Microarray | GSE66099 | -16.7 | 9758100 |
| SOCS3 | Antecedent | Microarray | GSE69528 | -15.3 | 10660535 |
| STAT4 | Antecedent | Microarray | GSE49753 | -13.0 | 11257135 |
| VIM | Antecedent | Microarray | GSE64457 | -11.0 | 15649310 |
| HK2 | Antecedent | Microarray | GSE9692 | -6.0 | 11737603 |
| PATZ1 | Subsequent | Microarray | GSE9692 | +18.2 | 41634312 |
| GALNT14 | Subsequent | Microarray | GSE9692 | +17.1 | 39981259 |
| PADI2 | Subsequent | Microarray | GSE9692 | +17.0 | 39405117 |
| MS4A4A | Subsequent | Microarray | GSE9692 | +12.1 | 32655806 |
| TSPO | Subsequent | Microarray | GSE28750 | +7.1 | 30497506 |
| RGL4 | Subsequent | Microarray | GSE64457 | +6.5 | 35965814 |
| ALOX5 | Subsequent | Microarray | GSE9692 | +5.8 | 23567086 |
| ZAP70 | Subsequent | Microarray | GSE28750 | +3.5 | 25557485 |
| PADI4 | Subsequent | Microarray | GSE66099 | +3.4 | 30044533 |
| PIM3 | Subsequent | Microarray | GSE66099 | +2.0 | 28469150 |

**Table S4.** Augmentation-revealed microarray gene signatures, Tuberculosis.

| Gene | Class | Platform | Dataset (GSE) | $\Delta$ (yr) | Earliest PMID |
| --- | --- | --- | --- | --- | --- |
| IMPACT | Antecedent | Microarray | GSE54992 | -67.8 | 21066594 |
| C2 | Antecedent | Microarray | GSE42832 | -47.8 | 4957443 |
| HP | Antecedent | Microarray | GSE42830 | -31.2 | 7179484 |
| TST | Antecedent | Microarray | GSE69581 | -23.8 | 8127325 |
| CD44 | Antecedent | Microarray | GSE54992 | -19.8 | 7513305 |
| FAS | Antecedent | Microarray | GSE42830 | -19.1 | 7523140 |
| IL1B | Antecedent | Microarray | GSE54992 | -15.1 | 10885842 |
| CXCR3 | Antecedent | Microarray | GSE54992 | -14.9 | 10092813 |
| STAT4 | Antecedent | Microarray | GSE54992 | -12.6 | 11441083 |
| MR1 | Antecedent | Microarray | GSE39939 | -3.8 | 20581831 |
| GADD45B | Subsequent | Microarray | GSE42830 | +11.4 | 40010539 |
| SLC2A3 | Subsequent | Microarray | GSE42825 | +11.3 | 39653283 |
| DCXR | Subsequent | Microarray | GSE42825 | +8.5 | 34313729 |
| ACSL4 | Subsequent | Microarray | GSE42825 | +7.1 | 33121683 |
| LDHA | Subsequent | Microarray | GSE54992 | +5.7 | 31201418 |
| CALCOCO2 | Subsequent | Microarray | GSE39939 | +5.7 | 33841389 |
| MGA | Subsequent | Microarray | GSE42832 | +4.9 | 29920796 |
| DDX58 | Subsequent | Microarray | GSE39939 | +4.7 | 31249631 |
| SLAMF1 | Subsequent | Microarray | GSE42830 | +1.8 | 26253450 |
| GZMK | Subsequent | Microarray | GSE42830 | +1.8 | 26156785 |

**Table S5.** Augmentation-revealed microarray gene signatures, Type 2 diabetes.

| Gene | Class | Platform | Dataset (GSE) | $\Delta$ (yr) | Earliest PMID |
| --- | --- | --- | --- | --- | --- |
| BAD | Antecedent | Microarray | GSE19420 | -42.6 | 5647821 |
| PC | Antecedent | Microarray | GSE76895 | -42.3 | 171802 |
| GCA | Antecedent | Microarray | GSE19420 | -34.2 | 974531 |
| HLA-B | Antecedent | Microarray | GSE19420 | -33.9 | 300603 |
| PLN | Antecedent | Microarray | GSE76895 | -29.9 | 2961642 |
| SCD | Antecedent | Microarray | GSE19420 | -22.9 | 3365140 |
| MCC | Antecedent | Microarray | GSE19420 | -18.8 | 1373457 |
| NOS2 | Antecedent | Microarray | GSE19420 | -16.3 | 9098436 |
| CXCR3 | Antecedent | Microarray | GSE76895 | -15.4 | 12145160 |
| BGLAP | Antecedent | Microarray | GSE166502 | -15.2 | 16189279 |
| NUAK1 | Subsequent | Microarray | GSE19420 | +15.0 | 41264843 |
| USP13 | Subsequent | Microarray | GSE19420 | +14.3 | 40035128 |
| IL1RAPL1 | Subsequent | Microarray | GSE19420 | +13.1 | 37702180 |
| HSPA12A | Subsequent | Microarray | GSE19420 | +12.8 | 37916327 |
| SDHA | Subsequent | Microarray | GSE19420 | +6.7 | 28437656 |
| TMPRSS2 | Subsequent | Microarray | GSE19420 | +5.1 | 27790247 |
| HEY1 | Subsequent | Microarray | GSE19420 | +4.1 | 26191142 |
| KCNMA1 | Subsequent | Microarray | GSE19420 | +3.5 | 23646921 |
| ME2 | Subsequent | Microarray | GSE19420 | +2.1 | 23221071 |
| YAP1 | Subsequent | Microarray | GSE19420 | +1.2 | 22320911 |

**Table S6.**
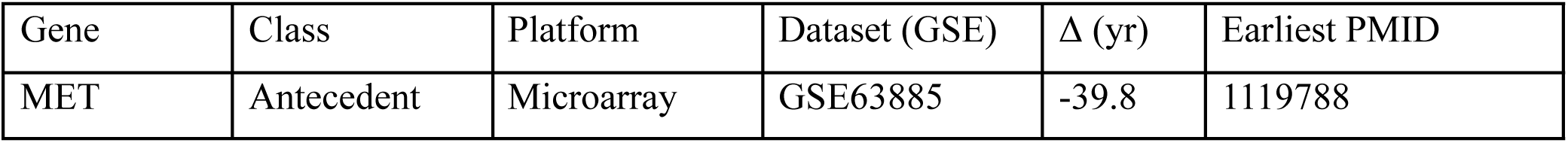
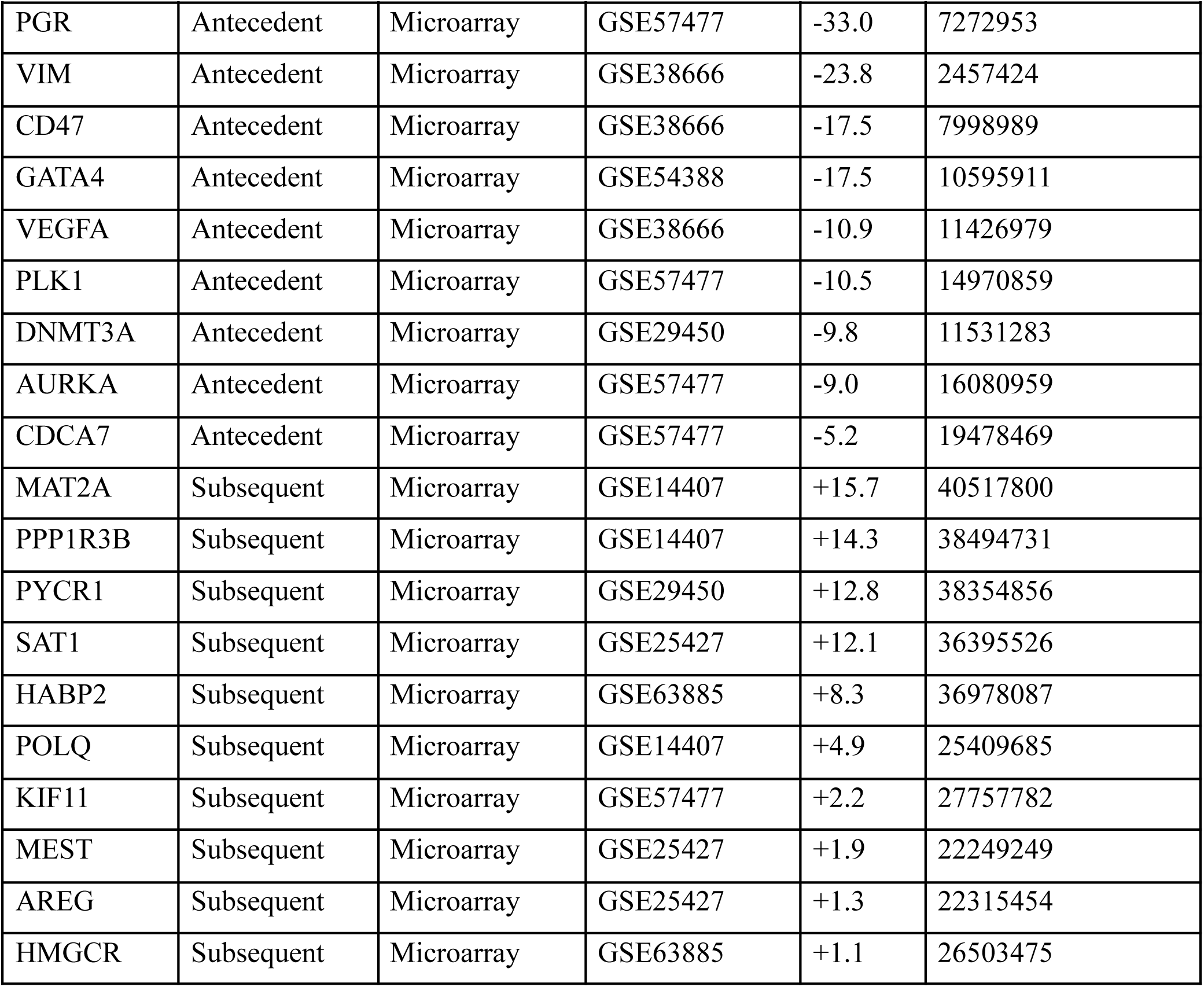
Augmentation-revealed microarray gene signatures, Ovarian cancer.

| Gene | Class | Platform | Dataset (GSE) | $\Delta$ (yr) | Earliest PMID |
| --- | --- | --- | --- | --- | --- |
| MET | Antecedent | Microarray | GSE63885 | -39.8 | 1119788 |
| PGR | Antecedent | Microarray | GSE57477 | -33.0 | 7272953 |
| VIM | Antecedent | Microarray | GSE38666 | -23.8 | 2457424 |
| CD47 | Antecedent | Microarray | GSE38666 | -17.5 | 7998989 |
| GATA4 | Antecedent | Microarray | GSE54388 | -17.5 | 10595911 |
| VEGFA | Antecedent | Microarray | GSE38666 | -10.9 | 11426979 |
| PLK1 | Antecedent | Microarray | GSE57477 | -10.5 | 14970859 |
| DNMT3A | Antecedent | Microarray | GSE29450 | -9.8 | 11531283 |
| AURKA | Antecedent | Microarray | GSE57477 | -9.0 | 16080959 |
| CDCA7 | Antecedent | Microarray | GSE57477 | -5.2 | 19478469 |
| MAT2A | Subsequent | Microarray | GSE14407 | +15.7 | 40517800 |
| PPP1R3B | Subsequent | Microarray | GSE14407 | +14.3 | 38494731 |
| PYCR1 | Subsequent | Microarray | GSE29450 | +12.8 | 38354856 |
| SAT1 | Subsequent | Microarray | GSE25427 | +12.1 | 36395526 |
| HABP2 | Subsequent | Microarray | GSE63885 | +8.3 | 36978087 |
| POLQ | Subsequent | Microarray | GSE14407 | +4.9 | 25409685 |
| KIF11 | Subsequent | Microarray | GSE57477 | +2.2 | 27757782 |
| MEST | Subsequent | Microarray | GSE25427 | +1.9 | 22249249 |
| AREG | Subsequent | Microarray | GSE25427 | +1.3 | 22315454 |
| HMGCR | Subsequent | Microarray | GSE63885 | +1.1 | 26503475 |

**Table S7.**
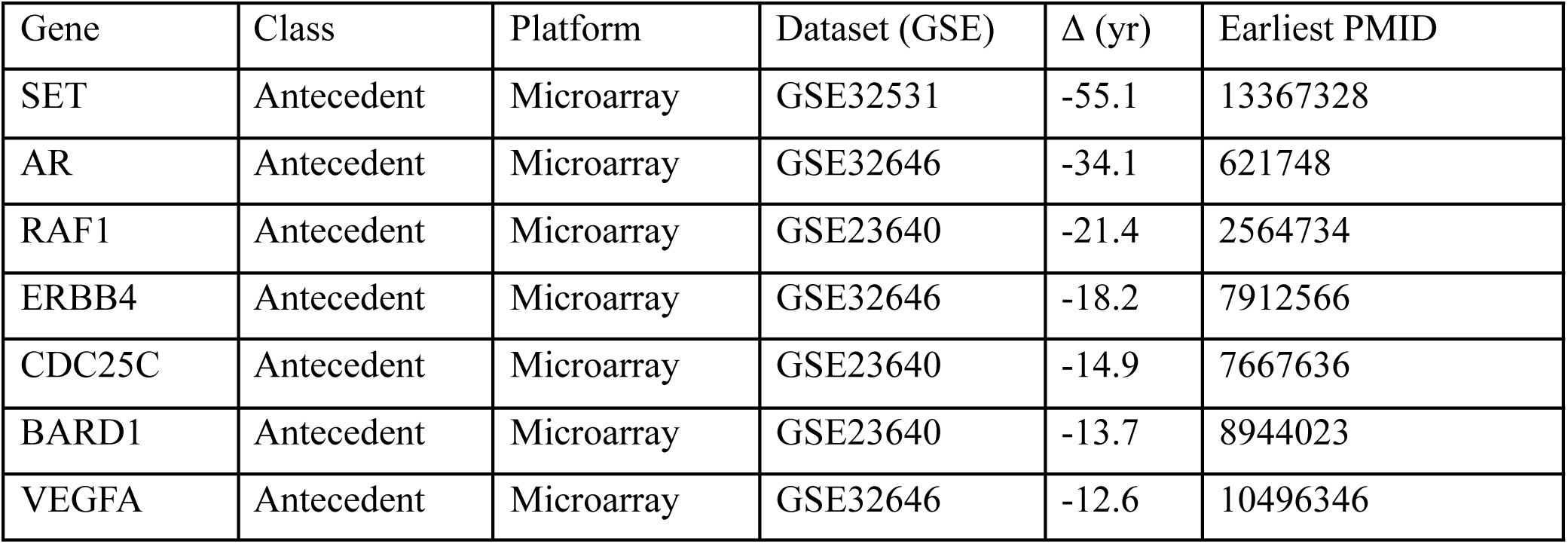
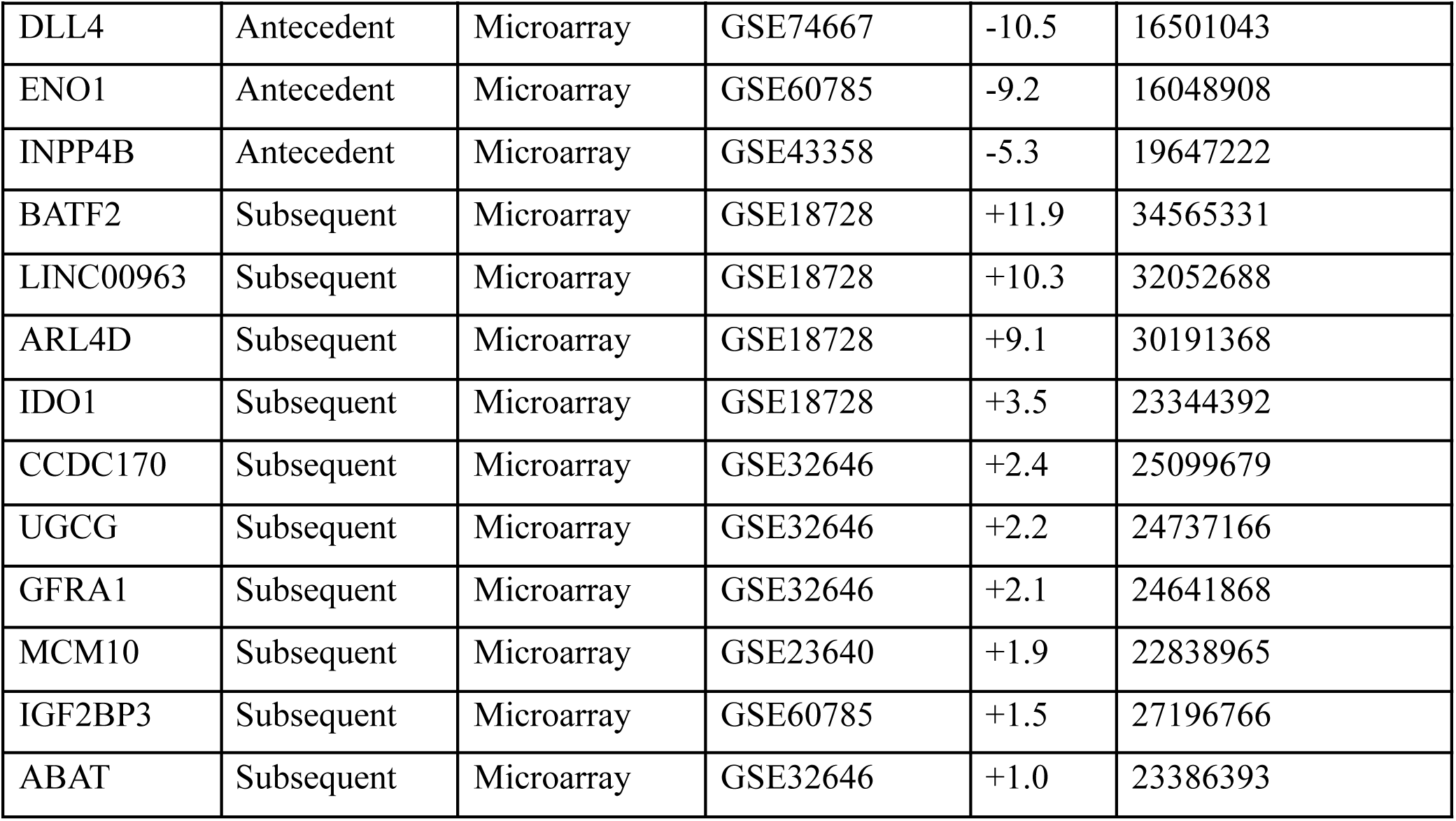
Augmentation-revealed microarray gene signatures, Breast cancer.

| Gene | Class | Platform | Dataset (GSE) | $\Delta$ (yr) | Earliest PMID |
| --- | --- | --- | --- | --- | --- |
| SET | Antecedent | Microarray | GSE32531 | -55.1 | 13367328 |
| AR | Antecedent | Microarray | GSE32646 | -34.1 | 621748 |
| RAF1 | Antecedent | Microarray | GSE23640 | -21.4 | 2564734 |
| ERBB4 | Antecedent | Microarray | GSE32646 | -18.2 | 7912566 |
| CDC25C | Antecedent | Microarray | GSE23640 | -14.9 | 7667636 |
| BARD1 | Antecedent | Microarray | GSE23640 | -13.7 | 8944023 |
| VEGFA | Antecedent | Microarray | GSE32646 | -12.6 | 10496346 |
| DLL4 | Antecedent | Microarray | GSE74667 | -10.5 | 16501043 |
| ENO1 | Antecedent | Microarray | GSE60785 | -9.2 | 16048908 |
| INPP4B | Antecedent | Microarray | GSE43358 | -5.3 | 19647222 |
| BATF2 | Subsequent | Microarray | GSE18728 | +11.9 | 34565331 |
| LINC00963 | Subsequent | Microarray | GSE18728 | +10.3 | 32052688 |
| ARL4D | Subsequent | Microarray | GSE18728 | +9.1 | 30191368 |
| IDO1 | Subsequent | Microarray | GSE18728 | +3.5 | 23344392 |
| CCDC170 | Subsequent | Microarray | GSE32646 | +2.4 | 25099679 |
| UGCG | Subsequent | Microarray | GSE32646 | +2.2 | 24737166 |
| GFRA1 | Subsequent | Microarray | GSE32646 | +2.1 | 24641868 |
| MCM10 | Subsequent | Microarray | GSE23640 | +1.9 | 22838965 |
| IGF2BP3 | Subsequent | Microarray | GSE60785 | +1.5 | 27196766 |
| ABAT | Subsequent | Microarray | GSE32646 | +1.0 | 23386393 |

**Table S8.** Augmentation-revealed RNA-seq signatures, Sepsis.

| Gene | Class | Platform | Dataset (GSE) | $\Delta$ (yr) | Earliest PMID |
| --- | --- | --- | --- | --- | --- |
| BCR | Antecedent | RNA-seq | GSE196117 | -27.3 | 7889565 |
| GAPDH | Antecedent | RNA-seq | GSE122274 | -23.9 | 7840203 |
| HMGCR | Antecedent | RNA-seq | GSE122274 | -17.0 | 11737603 |
| NFIL3 | Antecedent | RNA-seq | GSE133822 | -10.3 | 19341462 |
| ACHE | Antecedent | RNA-seq | GSE122274 | -8.6 | 20408892 |
| OAT | Antecedent | RNA-seq | GSE122274 | -8.0 | 20549270 |
| FASN | Antecedent | RNA-seq | GSE133822 | -7.6 | 23110240 |
| FUS | Antecedent | RNA-seq | GSE196117 | -6.0 | 27347057 |
| IDO1 | Antecedent | RNA-seq | GSE122274 | -4.4 | 24930766 |
| HLA-DPB1 | Antecedent | RNA-seq | GSE196117 | -1.5 | 33939725 |
| SKI | Subsequent | RNA-seq | GSE133822 | +6.5 | 41421230 |
| AMACR | Subsequent | RNA-seq | GSE133822 | +6.2 | 41107920 |
| DPEP2 | Subsequent | RNA-seq | GSE122274 | +6.1 | 41359645 |
| ASS1 | Subsequent | RNA-seq | GSE133822 | +5.9 | 40694828 |
| ZBTB24 | Subsequent | RNA-seq | GSE185263 | +4.1 | 41859080 |
| VAMP2 | Subsequent | RNA-seq | GSE196117 | +3.7 | 41809695 |
| GBA | Subsequent | RNA-seq | GSE185263 | +3.1 | 39950836 |
| SLC38A1 | Subsequent | RNA-seq | GSE185263 | +2.1 | 38228256 |
| ADAM8 | Subsequent | RNA-seq | GSE122274 | +1.7 | 32753701 |
| DDX24 | Subsequent | RNA-seq | GSE196117 | +1.2 | 37749550 |

**Table S9.**
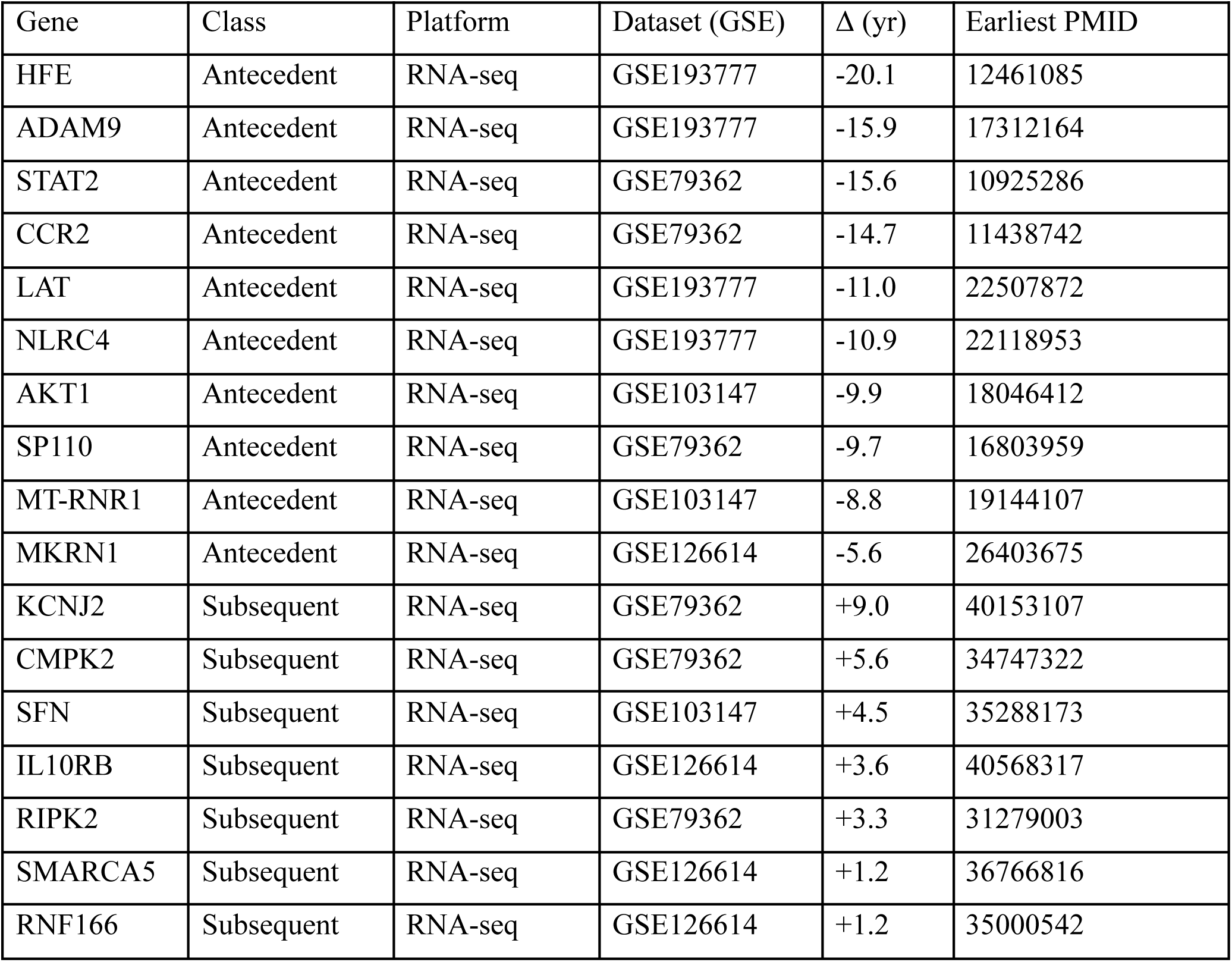
Augmentation-revealed RNA-seq signatures, Tuberculosis.

| Gene | Class | Platform | Dataset (GSE) | $\Delta$ (yr) | Earliest PMID |
| --- | --- | --- | --- | --- | --- |
| HFE | Antecedent | RNA-seq | GSE193777 | -20.1 | 12461085 |
| ADAM9 | Antecedent | RNA-seq | GSE193777 | -15.9 | 17312164 |
| STAT2 | Antecedent | RNA-seq | GSE79362 | -15.6 | 10925286 |
| CCR2 | Antecedent | RNA-seq | GSE79362 | -14.7 | 11438742 |
| LAT | Antecedent | RNA-seq | GSE193777 | -11.0 | 22507872 |
| NLRC4 | Antecedent | RNA-seq | GSE193777 | -10.9 | 22118953 |
| AKT1 | Antecedent | RNA-seq | GSE103147 | -9.9 | 18046412 |
| SP110 | Antecedent | RNA-seq | GSE79362 | -9.7 | 16803959 |
| MT-RNR1 | Antecedent | RNA-seq | GSE103147 | -8.8 | 19144107 |
| MKRN1 | Antecedent | RNA-seq | GSE126614 | -5.6 | 26403675 |
| KCNJ2 | Subsequent | RNA-seq | GSE79362 | +9.0 | 40153107 |
| CMPK2 | Subsequent | RNA-seq | GSE79362 | +5.6 | 34747322 |
| SFN | Subsequent | RNA-seq | GSE103147 | +4.5 | 35288173 |
| IL10RB | Subsequent | RNA-seq | GSE126614 | +3.6 | 40568317 |
| RIPK2 | Subsequent | RNA-seq | GSE79362 | +3.3 | 31279003 |
| SMARCA5 | Subsequent | RNA-seq | GSE126614 | +1.2 | 36766816 |
| RNF166 | Subsequent | RNA-seq | GSE126614 | +1.2 | 35000542 |

**Table S10.** Augmentation-revealed RNA-seq signatures, Type 2 diabetes.

| Gene | Class | Platform | Dataset (GSE) | $\Delta$ (yr) | Earliest PMID |
| --- | --- | --- | --- | --- | --- |
| PPP1R3B | Antecedent | RNA-seq | GSE253502 | -17.4 | 16907705 |
| IL18R1 | Antecedent | RNA-seq | GSE253502 | -15.1 | 18840782 |
| SLC5A2 | Antecedent | RNA-seq | GSE184050 | -12.4 | 19324482 |
| TMEM163 | Antecedent | RNA-seq | GSE184050 | -8.5 | 23209189 |
| PDGFRB | Antecedent | RNA-seq | GSE184050 | -7.0 | 23337395 |
| IRF5 | Antecedent | RNA-seq | GSE253502 | -6.8 | 28082458 |
| OAT | Antecedent | RNA-seq | GSE253502 | -5.5 | 30035378 |
| CCN3 | Antecedent | RNA-seq | GSE184050 | -4.1 | 28809783 |
| TERT | Antecedent | RNA-seq | GSE184050 | -3.6 | 29410390 |
| LY86 | Antecedent | RNA-seq | GSE184050 | -2.8 | 30328000 |
| GRAMD2B | Subsequent | RNA-seq | GSE184050 | +4.7 | 41986506 |
| PRMT5 | Subsequent | RNA-seq | GSE184050 | +3.3 | 39454821 |
| CYB5R2 | Subsequent | RNA-seq | GSE184050 | +2.8 | 39218018 |
| EMB | Subsequent | RNA-seq | GSE253502 | +2.4 | 41936848 |
| CSK | Subsequent | RNA-seq | GSE253502 | +2.2 | 41968706 |
| COQ9 | Subsequent | RNA-seq | GSE184050 | +2.0 | 37516222 |
| STYXL1 | Subsequent | RNA-seq | GSE184050 | +2.0 | 37679740 |
| LST1 | Subsequent | RNA-seq | GSE253502 | +1.9 | 41488617 |

**Table S11.**
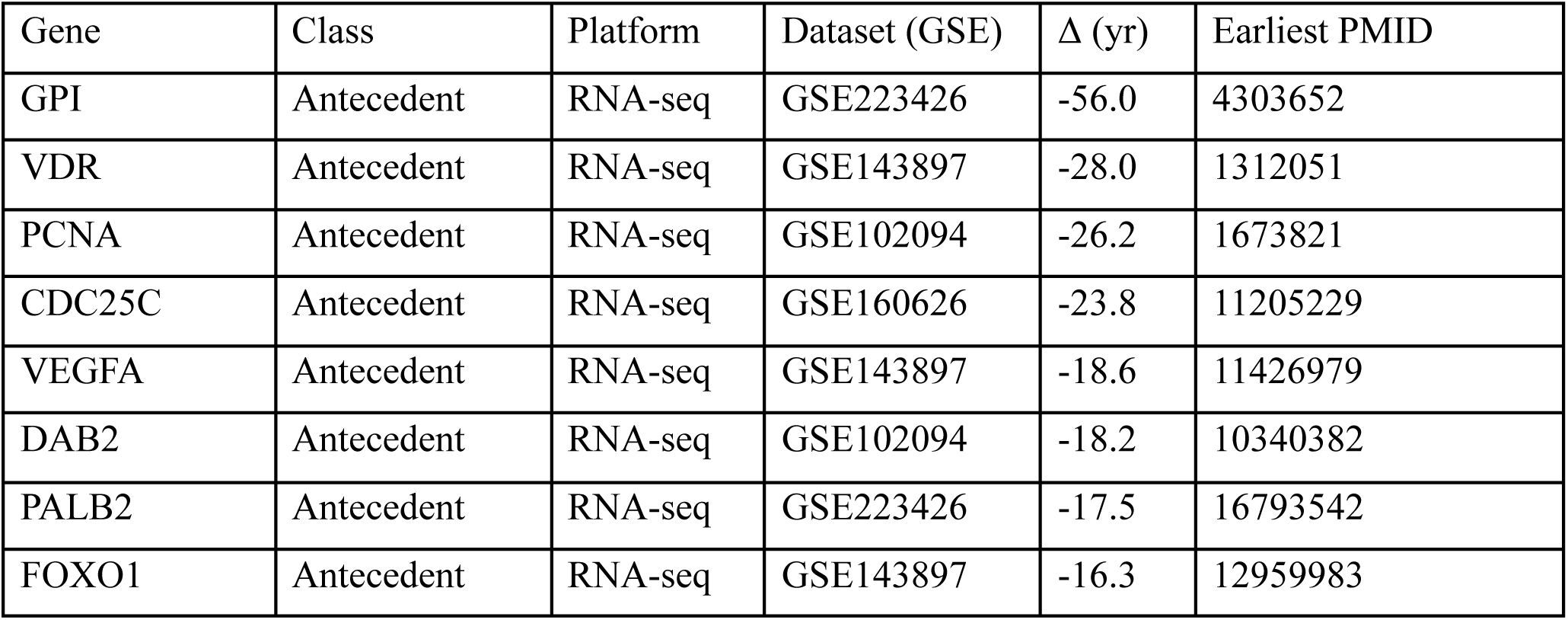
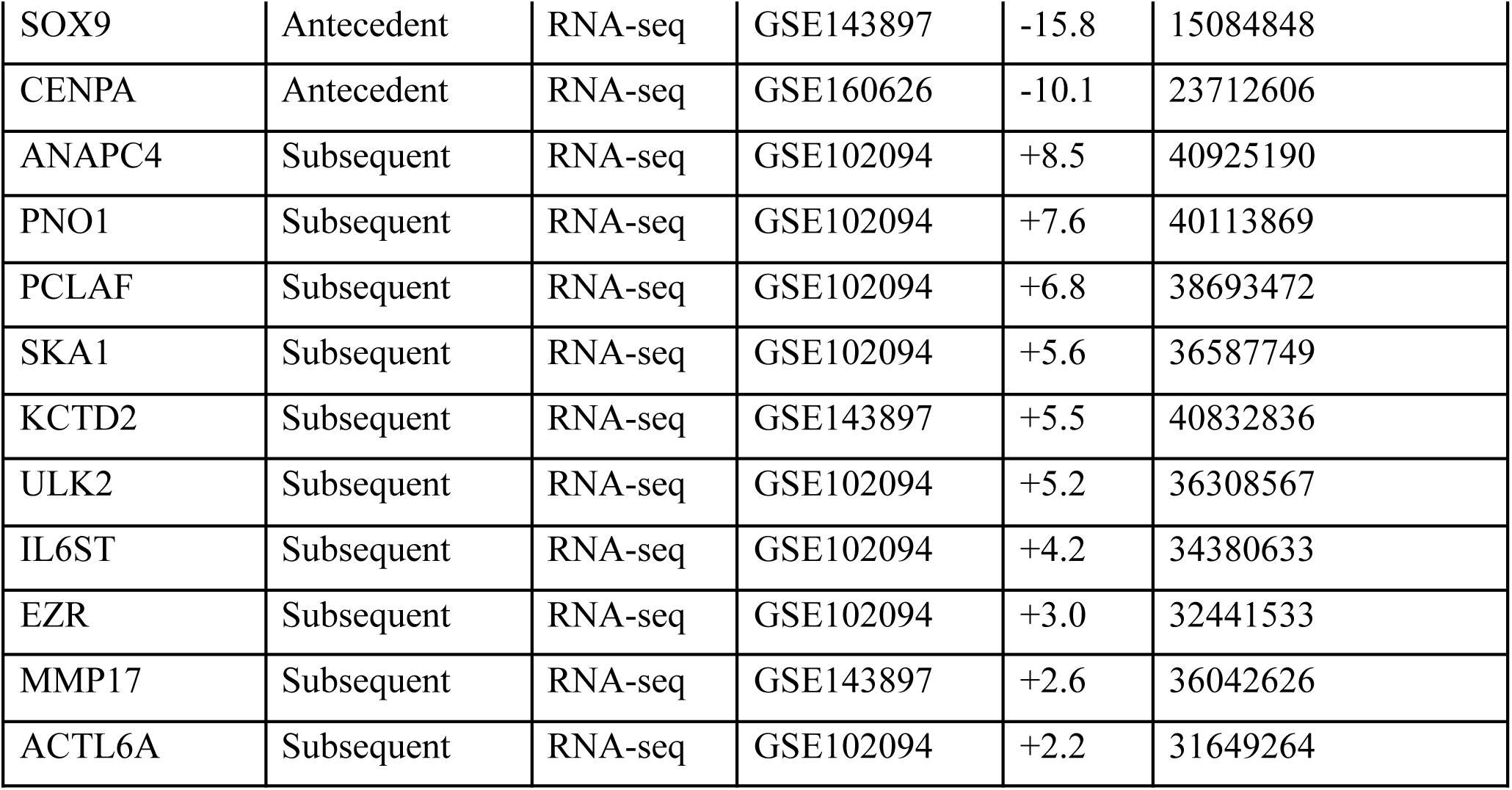
Augmentation-revealed RNA-seq signatures, Ovarian cancer.

| Gene | Class | Platform | Dataset (GSE) | $\Delta$ (yr) | Earliest PMID |
| --- | --- | --- | --- | --- | --- |
| GPI | Antecedent | RNA-seq | GSE223426 | -56.0 | 4303652 |
| VDR | Antecedent | RNA-seq | GSE143897 | -28.0 | 1312051 |
| PCNA | Antecedent | RNA-seq | GSE102094 | -26.2 | 1673821 |
| CDC25C | Antecedent | RNA-seq | GSE160626 | -23.8 | 11205229 |
| VEGFA | Antecedent | RNA-seq | GSE143897 | -18.6 | 11426979 |
| DAB2 | Antecedent | RNA-seq | GSE102094 | -18.2 | 10340382 |
| PALB2 | Antecedent | RNA-seq | GSE223426 | -17.5 | 16793542 |
| FOXO1 | Antecedent | RNA-seq | GSE143897 | -16.3 | 12959983 |
| SOX9 | Antecedent | RNA-seq | GSE143897 | -15.8 | 15084848 |
| CENPA | Antecedent | RNA-seq | GSE160626 | -10.1 | 23712606 |
| ANAPC4 | Subsequent | RNA-seq | GSE102094 | +8.5 | 40925190 |
| PNO1 | Subsequent | RNA-seq | GSE102094 | +7.6 | 40113869 |
| PCLAF | Subsequent | RNA-seq | GSE102094 | +6.8 | 38693472 |
| SKA1 | Subsequent | RNA-seq | GSE102094 | +5.6 | 36587749 |
| KCTD2 | Subsequent | RNA-seq | GSE143897 | +5.5 | 40832836 |
| ULK2 | Subsequent | RNA-seq | GSE102094 | +5.2 | 36308567 |
| IL6ST | Subsequent | RNA-seq | GSE102094 | +4.2 | 34380633 |
| EZR | Subsequent | RNA-seq | GSE102094 | +3.0 | 32441533 |
| MMP17 | Subsequent | RNA-seq | GSE143897 | +2.6 | 36042626 |
| ACTL6A | Subsequent | RNA-seq | GSE102094 | +2.2 | 31649264 |

**Table S12.** Augmentation-revealed RNA-seq signatures, Breast cancer.

| Gene | Class | Platform | Dataset (GSE) | $\Delta$ (yr) | Earliest PMID |
| --- | --- | --- | --- | --- | --- |
| HLA-A | Antecedent | RNA-seq | GSE52194 | -31.9 | 6181407 |
| GAPDH | Antecedent | RNA-seq | GSE52194 | -21.3 | 1638514 |
| MSN | Antecedent | RNA-seq | GSE52194 | -15.3 | 9706140 |
| NMI | Antecedent | RNA-seq | GSE52194 | -11.4 | 11916966 |
| ALK | Antecedent | RNA-seq | GSE52194 | -11.4 | 12115586 |
| RAB11A | Antecedent | RNA-seq | GSE52194 | -9.8 | 14557597 |
| LRIG1 | Antecedent | RNA-seq | GSE52194 | -8.8 | 16168117 |
| KRT18 | Antecedent | RNA-seq | GSE58135 | -6.0 | 18533032 |
| FLNA | Antecedent | RNA-seq | GSE52194 | -3.7 | 20179208 |
| P4HA1 | Antecedent | RNA-seq | GSE52194 | -3.5 | 20723428 |
| GALNT7 | Subsequent | RNA-seq | GSE58135 | +9.2 | 37633945 |
| TIPARP | Subsequent | RNA-seq | GSE52194 | +7.3 | 33799807 |
| CDK17 | Subsequent | RNA-seq | GSE52194 | +7.2 | 33868579 |
| PDE7A | Subsequent | RNA-seq | GSE58135 | +6.9 | 33685991 |
| PATJ | Subsequent | RNA-seq | GSE52194 | +6.1 | 31096822 |
| CREB3L4 | Subsequent | RNA-seq | GSE58135 | +4.6 | 30483806 |
| ACTA2 | Subsequent | RNA-seq | GSE52194 | +3.7 | 28881584 |
| ZMYND8 | Subsequent | RNA-seq | GSE52194 | +3.3 | 28055972 |
| FGF21 | Subsequent | RNA-seq | GSE52194 | +3.1 | 28106545 |
| CGAS | Subsequent | RNA-seq | GSE58135 | +2.6 | 27707838 |

**Table S13.**
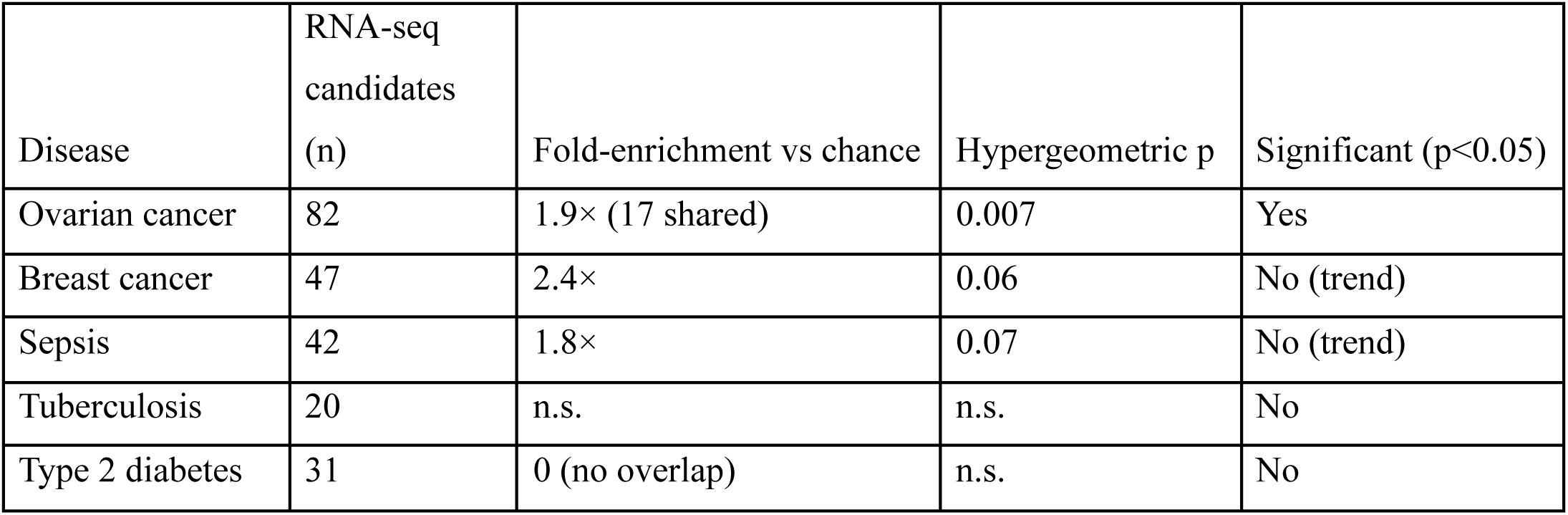
Cross-technology concordance of augmentation-revealed genes. Overlap between augmentation-revealed RNA-seq candidates and the microarray augmentation-revealed candidates for the same disease, tested by the hypergeometric distribution against a transcriptome background. Single-gene concordance is modest and tissue-dependent; the technology-generalisation claim rests primarily on recovery of independently verified disease biology and on pathway-level agreement (Tables S1 and S2).

**Table S14.** Augmentation-revealed RNA-seq pathways ranked by maximum enrichment depth (top 3 enrichment depths).

| Pathway | Depth | Disease |
| --- | --- | --- |
| positive regulation of nuclear-transcribed mRNA catabolic process, deadenylation-dependent decay | 12 | Sepsis |
| regulation of nuclear-transcribed mRNA catabolic process, deadenylation-dependent decay | 11 | Sepsis |
| T-helper cell differentiation | 11 | Sepsis |
| mRNA stabilization | 11 | Sepsis |
| positive regulation of T cell differentiation | 10 | Sepsis |
| regulation of mRNA stability | 10 | Sepsis |
| positive regulation of type II interferon production | 10 | Sepsis |
| positive regulation of tumor necrosis factor production | 11 | Tuberculosis |
| positive regulation of type III hypersensitivity | 11 | Tuberculosis |
| negative regulation of eosinophil degranulation | 11 | Tuberculosis |
| regulation of T cell cytokine production | 10 | Tuberculosis |
| positive regulation of tumor necrosis factor superfamily cytokine production | 10 | Tuberculosis |
| regulation of tumor necrosis factor production | 10 | Tuberculosis |
| positive regulation of peptidyl-tyrosine phosphorylation | 10 | Tuberculosis |
| meiosis I spindle assembly checkpoint signaling | 13 | Ovarian cancer |
| mitotic spindle assembly checkpoint signaling | 12 | Ovarian cancer |
| mitotic G2 DNA damage checkpoint signaling | 12 | Ovarian cancer |
| meiotic spindle assembly checkpoint signaling | 12 | Ovarian cancer |
| positive regulation of ubiquitin protein ligase activity | 12 | Ovarian cancer |
| positive regulation of MAP kinase activity | 12 | Ovarian cancer |
| negative regulation of mitotic metaphase/anaphase transition | 11 | Ovarian cancer |
| T-helper 2 cell differentiation | 12 | Breast cancer |
| positive regulation of miRNA transcription | 11 | Breast cancer |
| negative regulation of miRNA transcription | 11 | Breast cancer |
| kainate selective glutamate receptor activity | 11 | Breast cancer |
| ATP-activated inward rectifier potassium channel activity | 11 | Breast cancer |
| regulation of miRNA transcription | 10 | Breast cancer |
| miRNA transcription | 10 | Breast cancer |
| pyruvate decarboxylation to acetyl-CoA | 13 | Type 2 diabetes |
| mRNA cis splicing, via spliceosome | 12 | Type 2 diabetes |
| acetyl-CoA biosynthetic process | 12 | Type 2 diabetes |
| glycolytic process | 12 | Type 2 diabetes |
| mRNA splicing, via spliceosome | 11 | Type 2 diabetes |
| acyl-CoA biosynthetic process | 11 | Type 2 diabetes |
| regulation of pyruvate decarboxylation to acetyl-CoA | 11 | Type 2 diabetes |
| toll-like receptor 2 signaling pathway | 13 | Tuberculosis |
| toll-like receptor 4 signaling pathway | 13 | Tuberculosis |
| toll-like receptor 9 signaling pathway | 13 | Tuberculosis |
| inflammasome-mediated signaling pathway | 12 | Tuberculosis |
| positive regulation of macrophage cytokine production | 12 | Tuberculosis |
| positive regulation of interferon-beta production | 11 | Tuberculosis |
| regulation of T-helper 17 cell lineage commitment | 14 | Sepsis |
| T-helper 17 cell differentiation | 12 | Sepsis |
| glycolytic process | 12 | Sepsis |
| fatty-acyl-CoA biosynthetic process | 12 | Sepsis |
| cGAS/STING signaling pathway | 12 | Ovarian cancer |
| 'de novo' IMP biosynthetic process | 12 | Ovarian cancer |
| 'de novo' AMP biosynthetic process | 12 | Ovarian cancer |

**Table S15.** Highly specific RNA-seq pathways recurrently enriched across two or more datasets of the same disease. RNA-seq counterpart to Table S1.

| GO:Term | Biological Role | Relevance to Disease | Disease |
| --- | --- | --- | --- |
| T-helper cell differentiation | Commitment of naive CD4+ T cells to helper lineages coordinating adaptive immunity. | Skewed T-helper differentiation underlies the adaptive-immune dysfunction of sepsis. | Sepsis |
| alpha-beta T cell differentiation | Maturation of conventional alpha-beta T lymphocytes. | Impaired alpha-beta T-cell maturation marks the immunosuppressive phase of sepsis. | Sepsis |
| CD4-positive, alpha-beta T cell differentiation | Differentiation of CD4+ helper T cells. | Loss of CD4+ T-cell function tracks sepsis severity and secondary infection. | Sepsis |
| positive regulation of lymphocyte differentiation | Promotes lymphocyte lineage commitment and expansion. | Blunted lymphocyte differentiation contributes to sepsis-induced immunoparalysis. | Sepsis |
| specific granule | Neutrophil secondary granule storing antimicrobial proteins. | Neutrophil granule remodelling shapes bacterial killing in sepsis. | Sepsis |
| ficolin-1-rich granule | Neutrophil granule releasing the complement-activating lectin ficolin-1. | Ficolin/complement activation drives innate defence and tissue injury in sepsis. | Sepsis |
| tertiary granule | Neutrophil gelatinase granule supporting migration and matrix remodelling. | Granule mobilisation governs neutrophil recruitment in sepsis. | Sepsis |
| regulation of leukocyte migration | Controls trafficking of immune cells to sites of infection. | Leukocyte recruitment to the lung granuloma is central to Mtb containment. | Tuberculosis |
| myeloid leukocyte activation | Activation of macrophages, monocytes and other myeloid cells. | Macrophage activation state determines control versus persistence of Mtb. | Tuberculosis |
| lymphocyte differentiation | Lineage commitment of T and B lymphocytes. | Adaptive lymphocyte responses are required for protective TB immunity. | Tuberculosis |
| chemotaxis | Directed cell movement along chemical gradients. | Chemokine-driven recruitment builds and maintains the tuberculous granuloma. | Tuberculosis |
| ficolin-1-rich granule | Neutrophil granule releasing the complement lectin ficolin-1. | Low-grade innate and complement activation accompanies the inflammatory component of type 2 diabetes. | Type 2 diabetes |
| chromosome passenger complex | Aurora-B-containing complex regulating chromosome segregation and cytokinesis. | Mis-segregation drives the chromosomal instability that defines high-grade serous disease. | Ovarian cancer |
| DNA replication origin binding | Licensing of replication origins for genome duplication. | Replication stress and origin dysregulation fuel HGSOC genomic instability. | Ovarian cancer |
| meiotic chromosome separation | Segregation of homologous chromosomes in meiosis. | Reactivated meiotic programmes promote aneuploidy in ovarian tumours. | Ovarian cancer |
| CENP-A containing chromatin assembly | Deposition of the centromeric histone variant CENP-A. | CENP-A over-loading mislocalises centromeres and drives chromosomal instability in HGSOC. | Ovarian cancer |
| positive regulation of mitotic cytokinesis | Promotes the final division of the cytoplasm at mitosis. | Cytokinesis failure yields tetraploid, unstable ovarian tumour cells. | Ovarian cancer |
| spindle midzone | Central spindle structure organising chromosome separation and cytokinesis. | Aberrant spindle function underlies the mitotic errors of serous carcinoma. | Ovarian cancer |
| female meiotic nuclear division | Meiotic nuclear division of the female germline. | Ectopic meiotic-division machinery contributes to ovarian genomic instability. | Ovarian cancer |
| protein localization to cilium | Targeting of proteins to the primary cilium signalling organelle. | Ciliary (Hedgehog) signalling is implicated in breast tumour growth. | Breast cancer |
| actin-based cell projection | Formation of actin-rich membrane protrusions such as filopodia and microvilli. | Cytoskeletal projections support breast-cancer migration and invasion. | Breast cancer |

**Table S16.** Survival gene hazard ratios, breast and ovarian cancer. Cox proportional-hazards results for the genes evaluated in both breast-cancer cohorts (TCGA-BRCA and METABRIC). HR, hazard ratio; CI, 95% confidence interval; n.e., not estimable. Elastic-Net Cox model parameters: TCGA alpha = 0.016, l1_ratio = 0.7, concordance index 0.63; METABRIC alpha = 0.021, l1_ratio = 0.7, concordance index 0.60. Ovarian cancer. Cox proportional-hazards results for the 15 genes significant in TCGA high-grade serous ovarian cancer (420 patients). L1-penalised Cox, 48-gene panel, cross-validated concordance index 0.60. HR, hazard ratio; CI, 95% confidence interval.

| Gene | TCGA HR (95% CI) | TCGA P | METABRIC HR (95% CI) | METABRIC P |
| --- | --- | --- | --- | --- |
| DLL4 | 1.62 (1.24-2.13) | 0.0005 | 1.63 (1.08-2.46) | 0.019 |
| SDR9C7 | 1.27 (1.02-1.57) | 0.032 | 1.57 (1.05-2.35) | 0.030 |
| ARL4D | 0.78 (0.65-0.93) | 0.006 | 0.62 (0.44-0.87) | 0.005 |
| REG3G | 1.92 (1.08-3.39) | 0.025 | 1.22 (0.86-1.74) | 0.265 |
| FAM172A | 1.96 (1.30-2.94) | 0.001 | 1.17 (0.92-1.49) | 0.191 |
| LINC00839 | 1.25 (1.11-1.40) | 0.0001 | 0.67 (0.42-1.05) | 0.082 |
| SLC19A2 | 0.73 (0.58-0.91) | 0.006 | 0.81 (0.54-1.21) | 0.302 |
| TTC39C | 0.73 (0.56-0.96) | 0.023 | 1.00 (0.89-1.13) | 0.981 |
| TAF4B | 0.80 (0.66-0.98) | 0.029 | 0.75 (0.54-1.05) | 0.094 |
| DLK2 | 0.83 (0.72-0.95) | 0.007 | 0.95 (0.78-1.17) | 0.635 |
| STOX2 | 0.86 (0.73-1.00) | 0.058 | 1.01 (0.88-1.16) | 0.864 |
| CGB | 1.18 (0.92-1.51) | 0.184 | 0.78 (0.49-1.24) | 0.298 |
| ZNF684 | 0.72 (0.48-1.08) | 0.116 | 0.86 (0.62-1.20) | 0.386 |
| TIPARP | 1.01 (0.78-1.30) | 0.965 | 0.98 (0.87-1.12) | 0.804 |
| COL6A5 | 0.88 (0.74-1.05) | 0.148 | 0.89 (0.60-1.31) | 0.546 |
| ZIM3 | n.e. | 0.991 | 1.32 (0.85-2.07) | 0.221 |

**Table S16.** Ovarian cancer. Cox proportional-hazards results for the 15 genes significant in TCGA high-grade serous ovarian cancer (420 patients). L1-penalised Cox, 48-gene panel, cross-validated concordance index 0.60. HR, hazard ratio; CI, 95% confidence interval.

| Gene | HR (95% CI) | P |
| --- | --- | --- |
| FCRLA | 0.35 (0.21-0.57) | 2.9e-05 |
| HABP2 | 0.53 (0.39-0.72) | 4.2e-05 |
| PDIA6 | 0.59 (0.46-0.76) | 4.4e-05 |
| ABCA13 | 0.52 (0.38-0.72) | 5.9e-05 |
| BACE1 | 0.71 (0.58-0.87) | 0.001 |
| GGCX | 1.80 (1.26-2.57) | 0.001 |
| KLHL8 | 1.60 (1.18-2.17) | 0.002 |
| PUF60 | 0.74 (0.59-0.93) | 0.009 |
| CCDC102B | 0.59 (0.39-0.89) | 0.012 |
| AKAP11 | 1.50 (1.09-2.07) | 0.012 |
| PCED1B | 0.79 (0.65-0.95) | 0.013 |
| GATA5 | 1.15 (1.03-1.29) | 0.015 |
| SCIRT | 0.76 (0.60-0.95) | 0.016 |
| NPW | 1.10 (1.00-1.21) | 0.047 |
| TEF | 1.28 (1.00-1.64) | 0.047 |

**Table S17.**
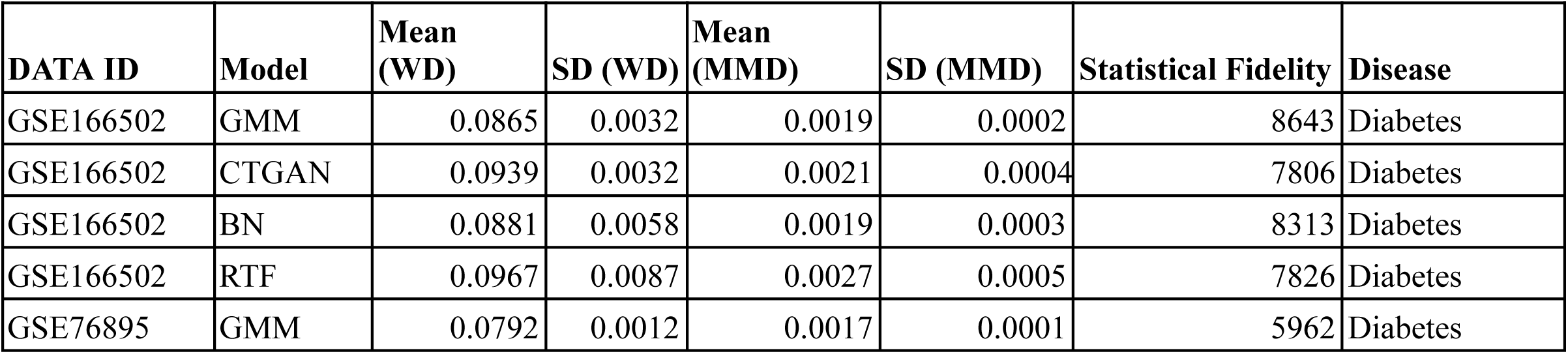
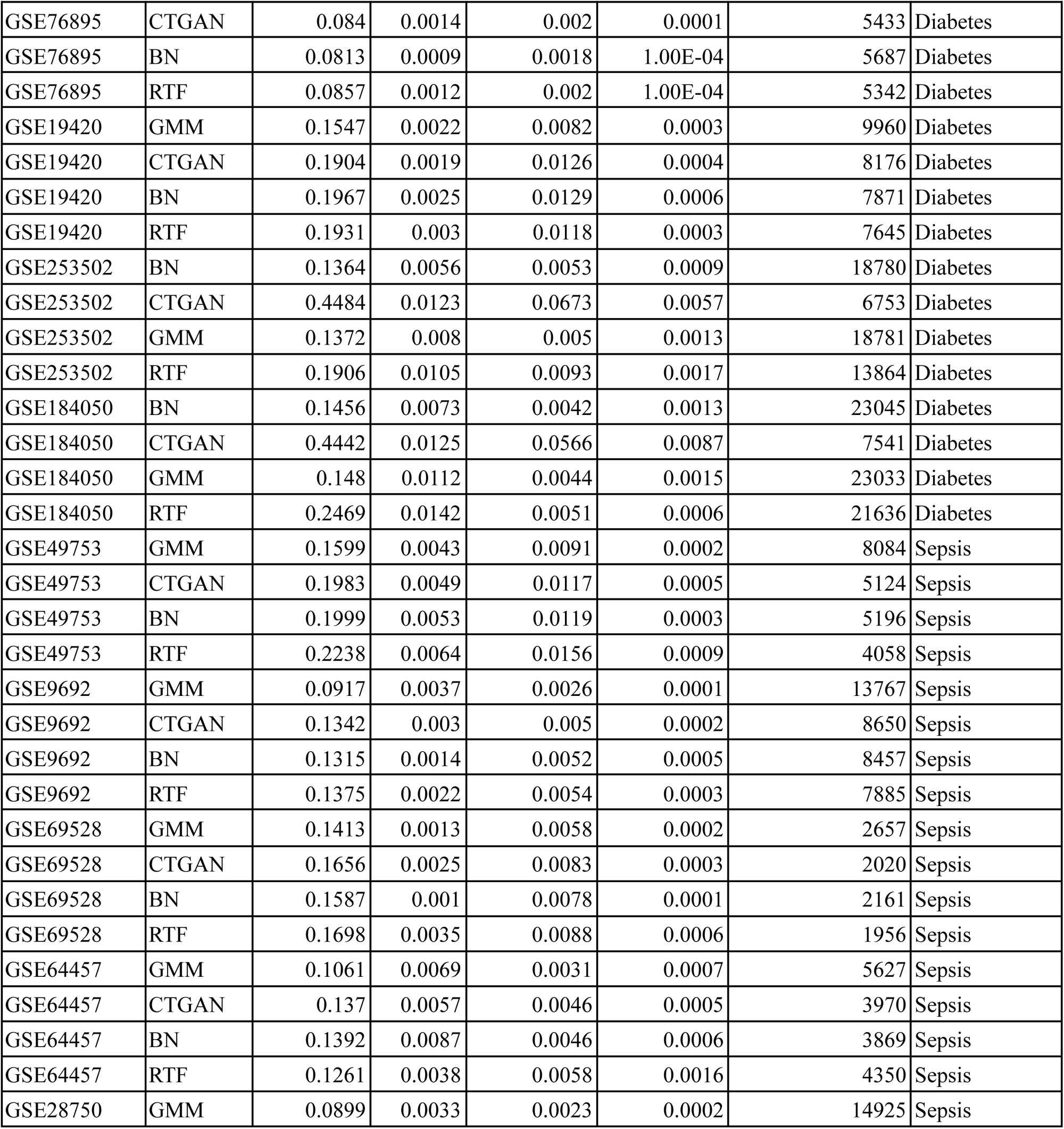
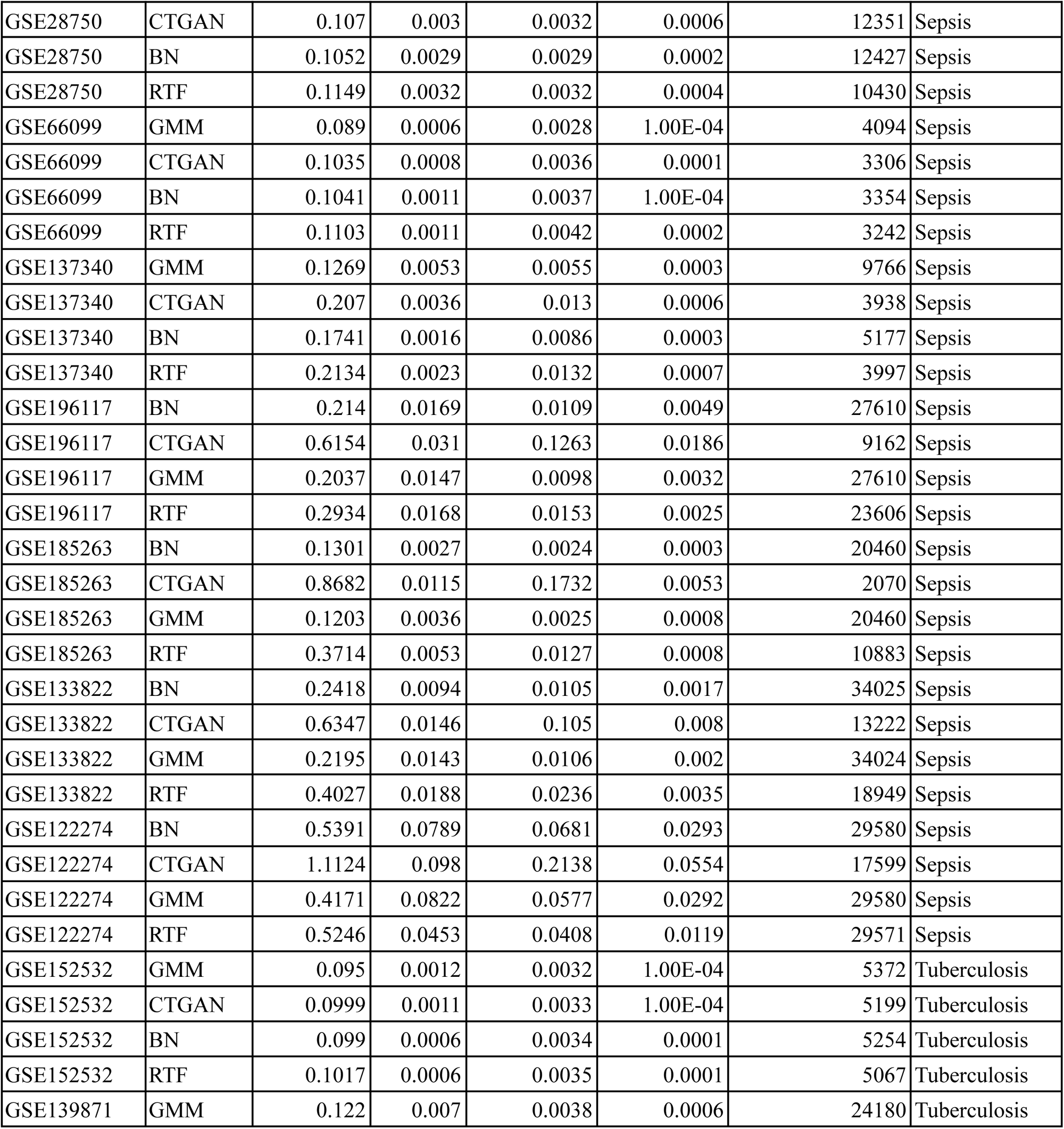
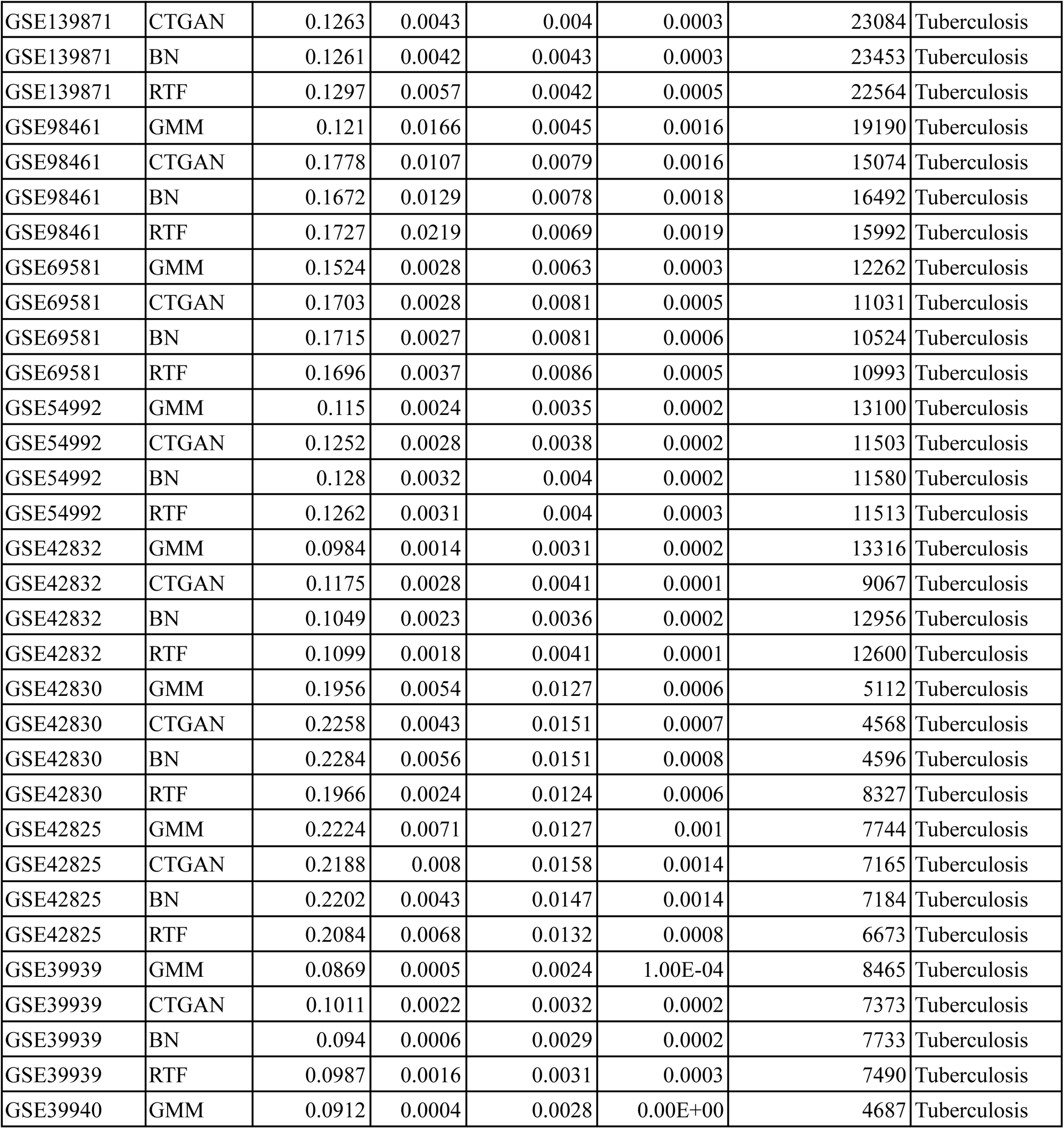
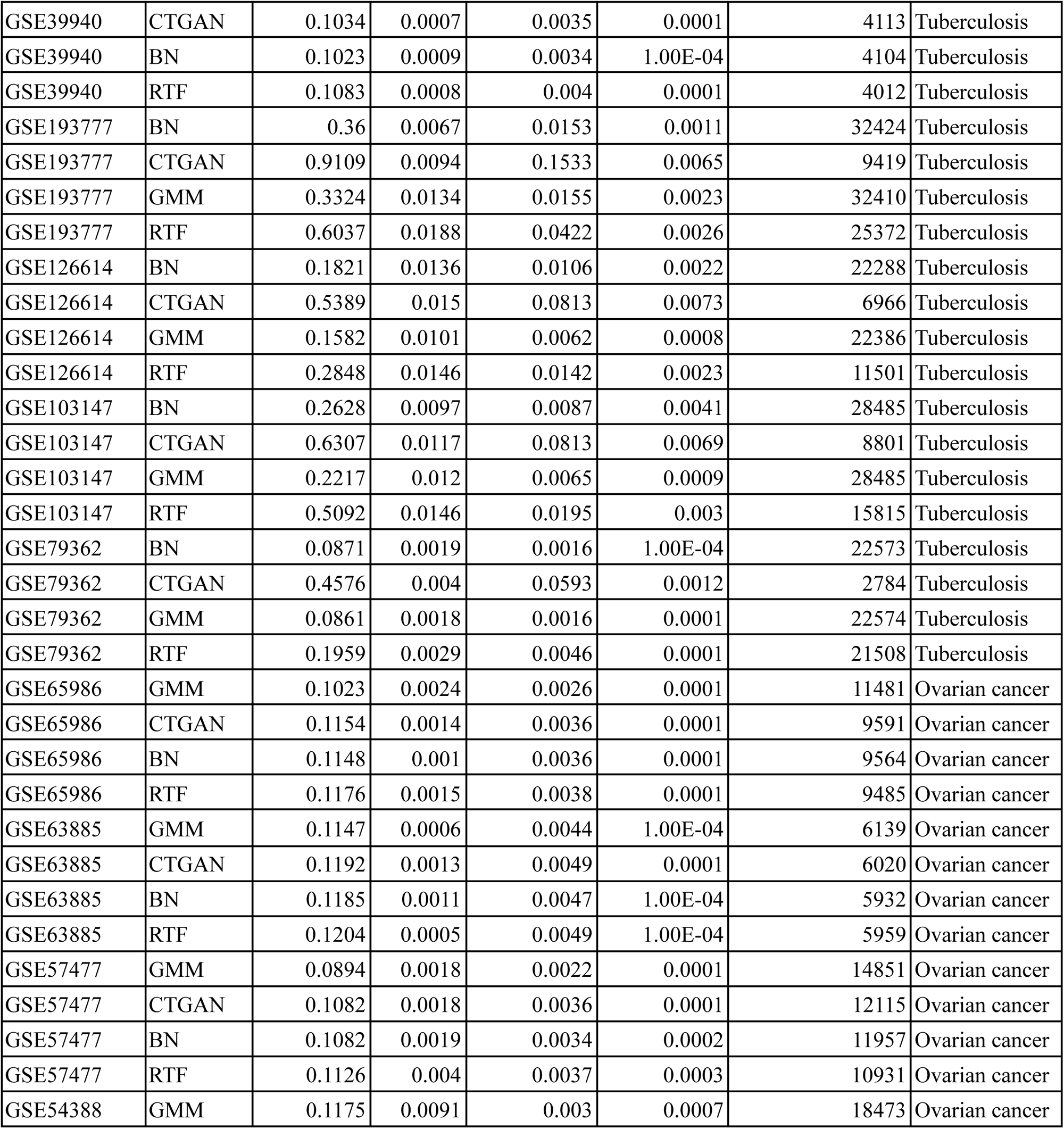
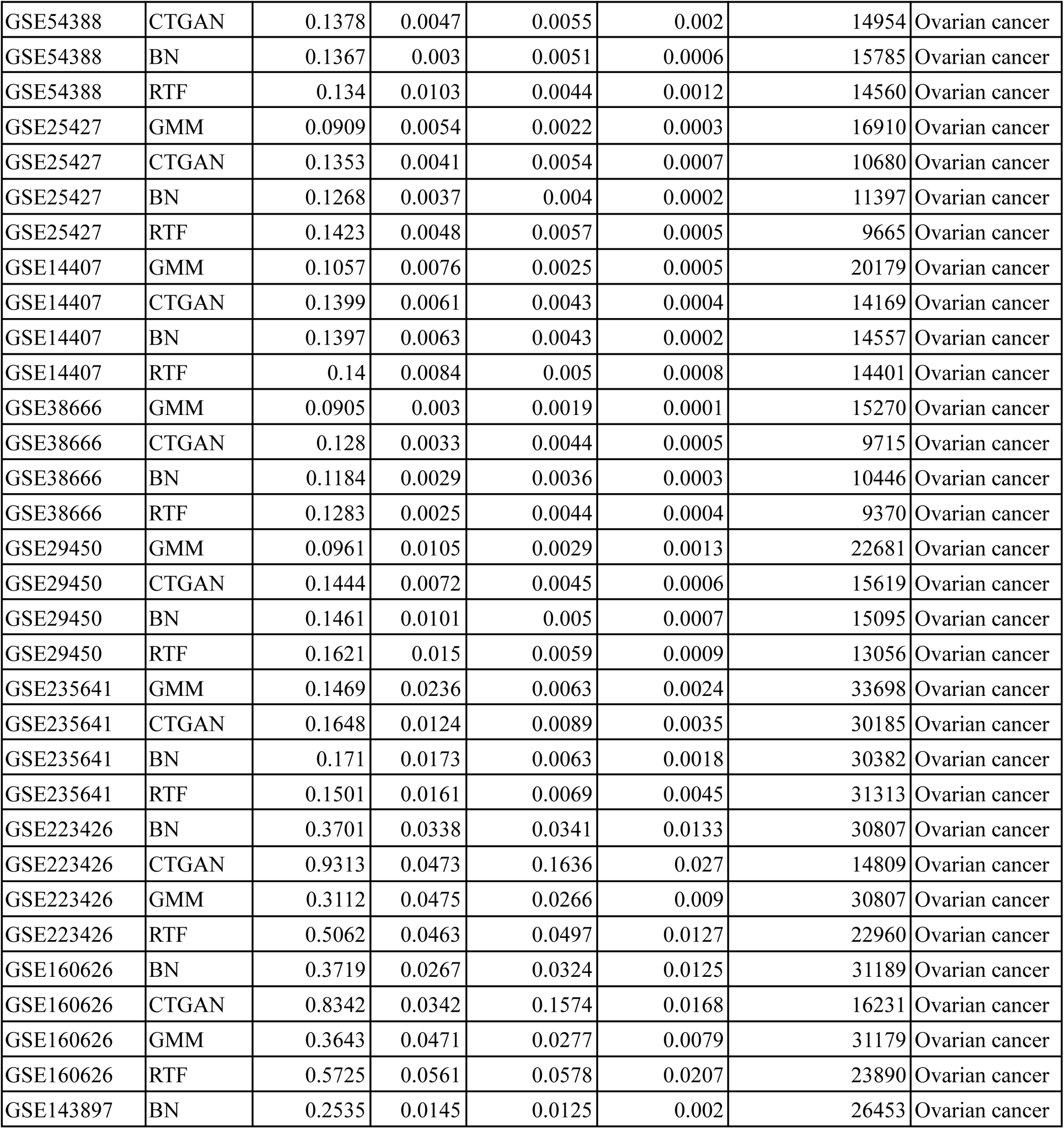
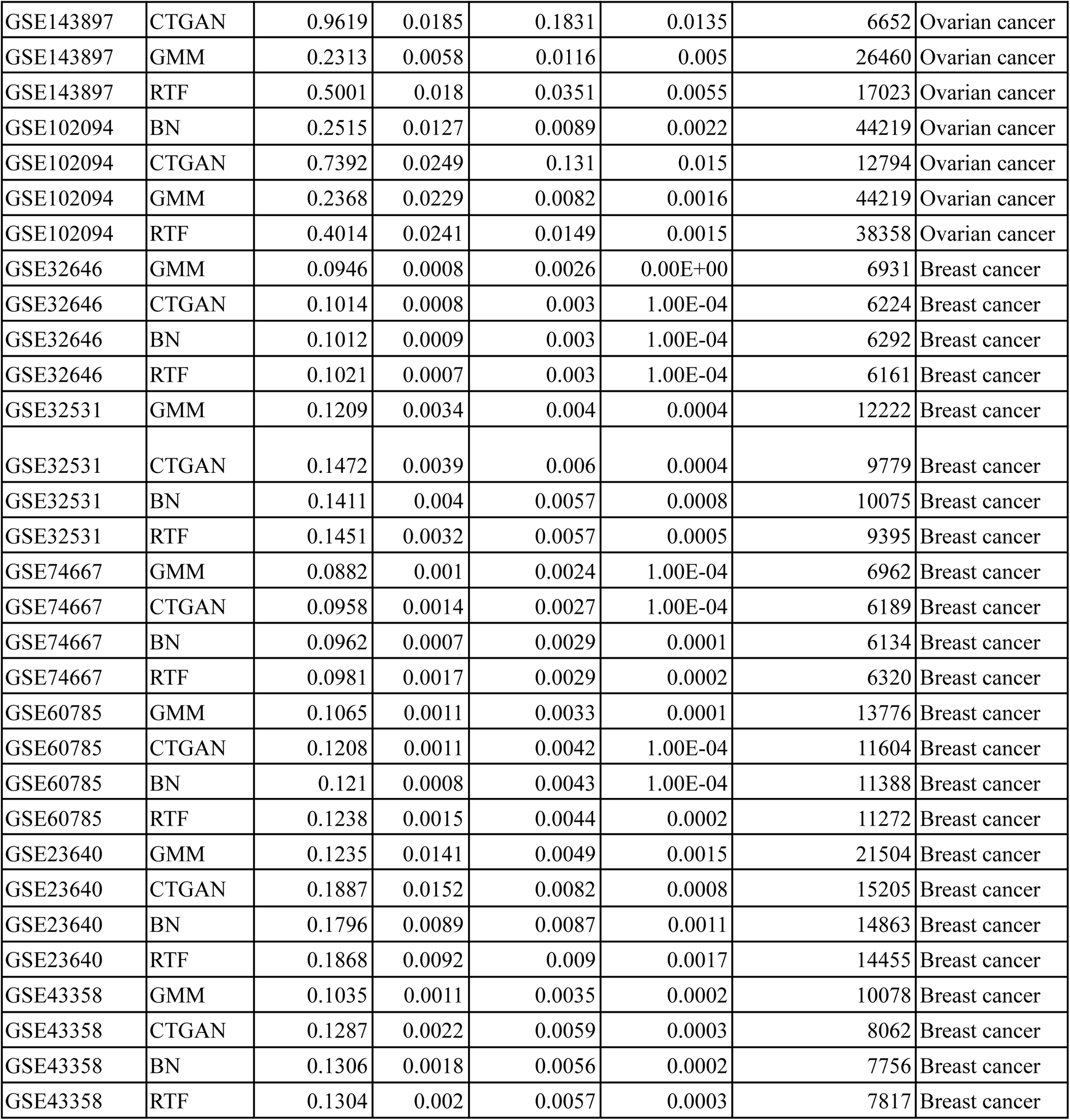
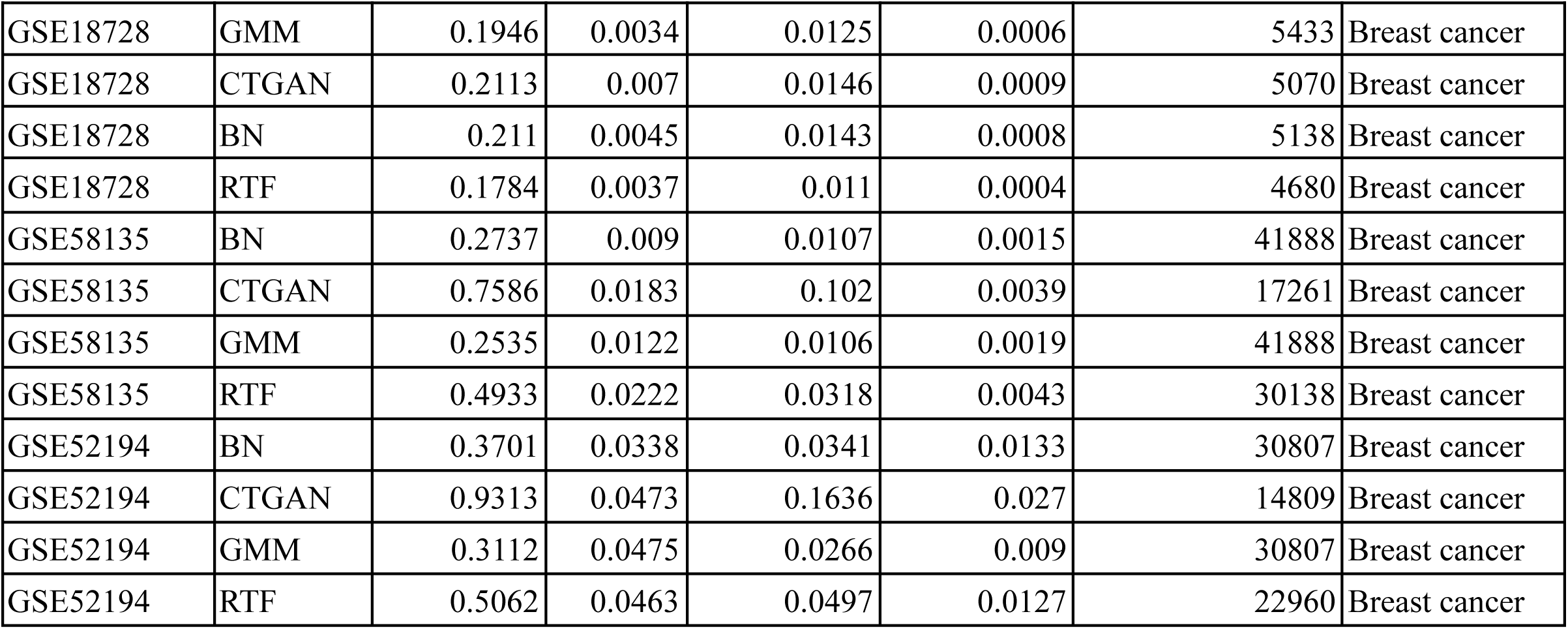
Benchmarking of generative approaches based on statistical fidelity metrics including Wasserstein Distance (WD), Maximum Mean Discrepancy (MMD) and t test. Mean and SD of the distance metrics are reported over 10 bootstrap iterations.

| DATA ID | Model | Mean (WD) | SD (WD) | Mean (MMD) | SD (MMD) | Statistical Fidelity | Disease |
| --- | --- | --- | --- | --- | --- | --- | --- |
| GSE166502 | GMM | 0.0865 | 0.0032 | 0.0019 | 0.0002 | 8643 | Diabetes |
| GSE166502 | CTGAN | 0.0939 | 0.0032 | 0.0021 | 0.0004 | 7806 | Diabetes |
| GSE166502 | BN | 0.0881 | 0.0058 | 0.0019 | 0.0003 | 8313 | Diabetes |
| GSE166502 | RTF | 0.0967 | 0.0087 | 0.0027 | 0.0005 | 7826 | Diabetes |
| GSE76895 | GMM | 0.0792 | 0.0012 | 0.0017 | 0.0001 | 5962 | Diabetes |
| GSE76895 | CTGAN | 0.084 | 0.0014 | 0.002 | 0.0001 | 5433 | Diabetes |
| GSE76895 | BN | 0.0813 | 0.0009 | 0.0018 | 1.00E-04 | 5687 | Diabetes |
| GSE76895 | RTF | 0.0857 | 0.0012 | 0.002 | 1.00E-04 | 5342 | Diabetes |
| GSE19420 | GMM | 0.1547 | 0.0022 | 0.0082 | 0.0003 | 9960 | Diabetes |
| GSE19420 | CTGAN | 0.1904 | 0.0019 | 0.0126 | 0.0004 | 8176 | Diabetes |
| GSE19420 | BN | 0.1967 | 0.0025 | 0.0129 | 0.0006 | 7871 | Diabetes |
| GSE19420 | RTF | 0.1931 | 0.003 | 0.0118 | 0.0003 | 7645 | Diabetes |
| GSE253502 | BN | 0.1364 | 0.0056 | 0.0053 | 0.0009 | 18780 | Diabetes |
| GSE253502 | CTGAN | 0.4484 | 0.0123 | 0.0673 | 0.0057 | 6753 | Diabetes |
| GSE253502 | GMM | 0.1372 | 0.008 | 0.005 | 0.0013 | 18781 | Diabetes |
| GSE253502 | RTF | 0.1906 | 0.0105 | 0.0093 | 0.0017 | 13864 | Diabetes |
| GSE184050 | BN | 0.1456 | 0.0073 | 0.0042 | 0.0013 | 23045 | Diabetes |
| GSE184050 | CTGAN | 0.4442 | 0.0125 | 0.0566 | 0.0087 | 7541 | Diabetes |
| GSE184050 | GMM | 0.148 | 0.0112 | 0.0044 | 0.0015 | 23033 | Diabetes |
| GSE184050 | RTF | 0.2469 | 0.0142 | 0.0051 | 0.0006 | 21636 | Diabetes |
| GSE49753 | GMM | 0.1599 | 0.0043 | 0.0091 | 0.0002 | 8084 | Sepsis |
| GSE49753 | CTGAN | 0.1983 | 0.0049 | 0.0117 | 0.0005 | 5124 | Sepsis |
| GSE49753 | BN | 0.1999 | 0.0053 | 0.0119 | 0.0003 | 5196 | Sepsis |
| GSE49753 | RTF | 0.2238 | 0.0064 | 0.0156 | 0.0009 | 4058 | Sepsis |
| GSE9692 | GMM | 0.0917 | 0.0037 | 0.0026 | 0.0001 | 13767 | Sepsis |
| GSE9692 | CTGAN | 0.1342 | 0.003 | 0.005 | 0.0002 | 8650 | Sepsis |
| GSE9692 | BN | 0.1315 | 0.0014 | 0.0052 | 0.0005 | 8457 | Sepsis |
| GSE9692 | RTF | 0.1375 | 0.0022 | 0.0054 | 0.0003 | 7885 | Sepsis |
| GSE69528 | GMM | 0.1413 | 0.0013 | 0.0058 | 0.0002 | 2657 | Sepsis |
| GSE69528 | CTGAN | 0.1656 | 0.0025 | 0.0083 | 0.0003 | 2020 | Sepsis |
| GSE69528 | BN | 0.1587 | 0.001 | 0.0078 | 0.0001 | 2161 | Sepsis |
| GSE69528 | RTF | 0.1698 | 0.0035 | 0.0088 | 0.0006 | 1956 | Sepsis |
| GSE64457 | GMM | 0.1061 | 0.0069 | 0.0031 | 0.0007 | 5627 | Sepsis |
| GSE64457 | CTGAN | 0.137 | 0.0057 | 0.0046 | 0.0005 | 3970 | Sepsis |
| GSE64457 | BN | 0.1392 | 0.0087 | 0.0046 | 0.0006 | 3869 | Sepsis |
| GSE64457 | RTF | 0.1261 | 0.0038 | 0.0058 | 0.0016 | 4350 | Sepsis |
| GSE28750 | GMM | 0.0899 | 0.0033 | 0.0023 | 0.0002 | 14925 | Sepsis |
| GSE28750 | CTGAN | 0.107 | 0.003 | 0.0032 | 0.0006 | 12351 | Sepsis |
| GSE28750 | BN | 0.1052 | 0.0029 | 0.0029 | 0.0002 | 12427 | Sepsis |
| GSE28750 | RTF | 0.1149 | 0.0032 | 0.0032 | 0.0004 | 10430 | Sepsis |
| GSE66099 | GMM | 0.089 | 0.0006 | 0.0028 | 1.00E-04 | 4094 | Sepsis |
| GSE66099 | CTGAN | 0.1035 | 0.0008 | 0.0036 | 0.0001 | 3306 | Sepsis |
| GSE66099 | BN | 0.1041 | 0.0011 | 0.0037 | 1.00E-04 | 3354 | Sepsis |
| GSE66099 | RTF | 0.1103 | 0.0011 | 0.0042 | 0.0002 | 3242 | Sepsis |
| GSE137340 | GMM | 0.1269 | 0.0053 | 0.0055 | 0.0003 | 9766 | Sepsis |
| GSE137340 | CTGAN | 0.207 | 0.0036 | 0.013 | 0.0006 | 3938 | Sepsis |
| GSE137340 | BN | 0.1741 | 0.0016 | 0.0086 | 0.0003 | 5177 | Sepsis |
| GSE137340 | RTF | 0.2134 | 0.0023 | 0.0132 | 0.0007 | 3997 | Sepsis |
| GSE196117 | BN | 0.214 | 0.0169 | 0.0109 | 0.0049 | 27610 | Sepsis |
| GSE196117 | CTGAN | 0.6154 | 0.031 | 0.1263 | 0.0186 | 9162 | Sepsis |
| GSE196117 | GMM | 0.2037 | 0.0147 | 0.0098 | 0.0032 | 27610 | Sepsis |
| GSE196117 | RTF | 0.2934 | 0.0168 | 0.0153 | 0.0025 | 23606 | Sepsis |
| GSE185263 | BN | 0.1301 | 0.0027 | 0.0024 | 0.0003 | 20460 | Sepsis |
| GSE185263 | CTGAN | 0.8682 | 0.0115 | 0.1732 | 0.0053 | 2070 | Sepsis |
| GSE185263 | GMM | 0.1203 | 0.0036 | 0.0025 | 0.0008 | 20460 | Sepsis |
| GSE185263 | RTF | 0.3714 | 0.0053 | 0.0127 | 0.0008 | 10883 | Sepsis |
| GSE133822 | BN | 0.2418 | 0.0094 | 0.0105 | 0.0017 | 34025 | Sepsis |
| GSE133822 | CTGAN | 0.6347 | 0.0146 | 0.105 | 0.008 | 13222 | Sepsis |
| GSE133822 | GMM | 0.2195 | 0.0143 | 0.0106 | 0.002 | 34024 | Sepsis |
| GSE133822 | RTF | 0.4027 | 0.0188 | 0.0236 | 0.0035 | 18949 | Sepsis |
| GSE122274 | BN | 0.5391 | 0.0789 | 0.0681 | 0.0293 | 29580 | Sepsis |
| GSE122274 | CTGAN | 1.1124 | 0.098 | 0.2138 | 0.0554 | 17599 | Sepsis |
| GSE122274 | GMM | 0.4171 | 0.0822 | 0.0577 | 0.0292 | 29580 | Sepsis |
| GSE122274 | RTF | 0.5246 | 0.0453 | 0.0408 | 0.0119 | 29571 | Sepsis |
| GSE152532 | GMM | 0.095 | 0.0012 | 0.0032 | 1.00E-04 | 5372 | Tuberculosis |
| GSE152532 | CTGAN | 0.0999 | 0.0011 | 0.0033 | 1.00E-04 | 5199 | Tuberculosis |
| GSE152532 | BN | 0.099 | 0.0006 | 0.0034 | 0.0001 | 5254 | Tuberculosis |
| GSE152532 | RTF | 0.1017 | 0.0006 | 0.0035 | 0.0001 | 5067 | Tuberculosis |
| GSE139871 | GMM | 0.122 | 0.007 | 0.0038 | 0.0006 | 24180 | Tuberculosis |
| GSE139871 | CTGAN | 0.1263 | 0.0043 | 0.004 | 0.0003 | 23084 | Tuberculosis |
| GSE139871 | BN | 0.1261 | 0.0042 | 0.0043 | 0.0003 | 23453 | Tuberculosis |
| GSE139871 | RTF | 0.1297 | 0.0057 | 0.0042 | 0.0005 | 22564 | Tuberculosis |
| GSE98461 | GMM | 0.121 | 0.0166 | 0.0045 | 0.0016 | 19190 | Tuberculosis |
| GSE98461 | CTGAN | 0.1778 | 0.0107 | 0.0079 | 0.0016 | 15074 | Tuberculosis |
| GSE98461 | BN | 0.1672 | 0.0129 | 0.0078 | 0.0018 | 16492 | Tuberculosis |
| GSE98461 | RTF | 0.1727 | 0.0219 | 0.0069 | 0.0019 | 15992 | Tuberculosis |
| GSE69581 | GMM | 0.1524 | 0.0028 | 0.0063 | 0.0003 | 12262 | Tuberculosis |
| GSE69581 | CTGAN | 0.1703 | 0.0028 | 0.0081 | 0.0005 | 11031 | Tuberculosis |
| GSE69581 | BN | 0.1715 | 0.0027 | 0.0081 | 0.0006 | 10524 | Tuberculosis |
| GSE69581 | RTF | 0.1696 | 0.0037 | 0.0086 | 0.0005 | 10993 | Tuberculosis |
| GSE54992 | GMM | 0.115 | 0.0024 | 0.0035 | 0.0002 | 13100 | Tuberculosis |
| GSE54992 | CTGAN | 0.1252 | 0.0028 | 0.0038 | 0.0002 | 11503 | Tuberculosis |
| GSE54992 | BN | 0.128 | 0.0032 | 0.004 | 0.0002 | 11580 | Tuberculosis |
| GSE54992 | RTF | 0.1262 | 0.0031 | 0.004 | 0.0003 | 11513 | Tuberculosis |
| GSE42832 | GMM | 0.0984 | 0.0014 | 0.0031 | 0.0002 | 13316 | Tuberculosis |
| GSE42832 | CTGAN | 0.1175 | 0.0028 | 0.0041 | 0.0001 | 9067 | Tuberculosis |
| GSE42832 | BN | 0.1049 | 0.0023 | 0.0036 | 0.0002 | 12956 | Tuberculosis |
| GSE42832 | RTF | 0.1099 | 0.0018 | 0.0041 | 0.0001 | 12600 | Tuberculosis |
| GSE42830 | GMM | 0.1956 | 0.0054 | 0.0127 | 0.0006 | 5112 | Tuberculosis |
| GSE42830 | CTGAN | 0.2258 | 0.0043 | 0.0151 | 0.0007 | 4568 | Tuberculosis |
| GSE42830 | BN | 0.2284 | 0.0056 | 0.0151 | 0.0008 | 4596 | Tuberculosis |
| GSE42830 | RTF | 0.1966 | 0.0024 | 0.0124 | 0.0006 | 8327 | Tuberculosis |
| GSE42825 | GMM | 0.2224 | 0.0071 | 0.0127 | 0.001 | 7744 | Tuberculosis |
| GSE42825 | CTGAN | 0.2188 | 0.008 | 0.0158 | 0.0014 | 7165 | Tuberculosis |
| GSE42825 | BN | 0.2202 | 0.0043 | 0.0147 | 0.0014 | 7184 | Tuberculosis |
| GSE42825 | RTF | 0.2084 | 0.0068 | 0.0132 | 0.0008 | 6673 | Tuberculosis |
| GSE39939 | GMM | 0.0869 | 0.0005 | 0.0024 | 1.00E-04 | 8465 | Tuberculosis |
| GSE39939 | CTGAN | 0.1011 | 0.0022 | 0.0032 | 0.0002 | 7373 | Tuberculosis |
| GSE39939 | BN | 0.094 | 0.0006 | 0.0029 | 0.0002 | 7733 | Tuberculosis |
| GSE39939 | RTF | 0.0987 | 0.0016 | 0.0031 | 0.0003 | 7490 | Tuberculosis |
| GSE39940 | GMM | 0.0912 | 0.0004 | 0.0028 | 0.00E+00 | 4687 | Tuberculosis |
| GSE39940 | CTGAN | 0.1034 | 0.0007 | 0.0035 | 0.0001 | 4113 | Tuberculosis |
| GSE39940 | BN | 0.1023 | 0.0009 | 0.0034 | 1.00E-04 | 4104 | Tuberculosis |
| GSE39940 | RTF | 0.1083 | 0.0008 | 0.004 | 0.0001 | 4012 | Tuberculosis |
| GSE193777 | BN | 0.36 | 0.0067 | 0.0153 | 0.0011 | 32424 | Tuberculosis |
| GSE193777 | CTGAN | 0.9109 | 0.0094 | 0.1533 | 0.0065 | 9419 | Tuberculosis |
| GSE193777 | GMM | 0.3324 | 0.0134 | 0.0155 | 0.0023 | 32410 | Tuberculosis |
| GSE193777 | RTF | 0.6037 | 0.0188 | 0.0422 | 0.0026 | 25372 | Tuberculosis |
| GSE126614 | BN | 0.1821 | 0.0136 | 0.0106 | 0.0022 | 22288 | Tuberculosis |
| GSE126614 | CTGAN | 0.5389 | 0.015 | 0.0813 | 0.0073 | 6966 | Tuberculosis |
| GSE126614 | GMM | 0.1582 | 0.0101 | 0.0062 | 0.0008 | 22386 | Tuberculosis |
| GSE126614 | RTF | 0.2848 | 0.0146 | 0.0142 | 0.0023 | 11501 | Tuberculosis |
| GSE103147 | BN | 0.2628 | 0.0097 | 0.0087 | 0.0041 | 28485 | Tuberculosis |
| GSE103147 | CTGAN | 0.6307 | 0.0117 | 0.0813 | 0.0069 | 8801 | Tuberculosis |
| GSE103147 | GMM | 0.2217 | 0.012 | 0.0065 | 0.0009 | 28485 | Tuberculosis |
| GSE103147 | RTF | 0.5092 | 0.0146 | 0.0195 | 0.003 | 15815 | Tuberculosis |
| GSE79362 | BN | 0.0871 | 0.0019 | 0.0016 | 1.00E-04 | 22573 | Tuberculosis |
| GSE79362 | CTGAN | 0.4576 | 0.004 | 0.0593 | 0.0012 | 2784 | Tuberculosis |
| GSE79362 | GMM | 0.0861 | 0.0018 | 0.0016 | 0.0001 | 22574 | Tuberculosis |
| GSE79362 | RTF | 0.1959 | 0.0029 | 0.0046 | 0.0001 | 21508 | Tuberculosis |
| GSE65986 | GMM | 0.1023 | 0.0024 | 0.0026 | 0.0001 | 11481 | Ovarian cancer |
| GSE65986 | CTGAN | 0.1154 | 0.0014 | 0.0036 | 0.0001 | 9591 | Ovarian cancer |
| GSE65986 | BN | 0.1148 | 0.001 | 0.0036 | 0.0001 | 9564 | Ovarian cancer |
| GSE65986 | RTF | 0.1176 | 0.0015 | 0.0038 | 0.0001 | 9485 | Ovarian cancer |
| GSE63885 | GMM | 0.1147 | 0.0006 | 0.0044 | 1.00E-04 | 6139 | Ovarian cancer |
| GSE63885 | CTGAN | 0.1192 | 0.0013 | 0.0049 | 0.0001 | 6020 | Ovarian cancer |
| GSE63885 | BN | 0.1185 | 0.0011 | 0.0047 | 1.00E-04 | 5932 | Ovarian cancer |
| GSE63885 | RTF | 0.1204 | 0.0005 | 0.0049 | 1.00E-04 | 5959 | Ovarian cancer |
| GSE57477 | GMM | 0.0894 | 0.0018 | 0.0022 | 0.0001 | 14851 | Ovarian cancer |
| GSE57477 | CTGAN | 0.1082 | 0.0018 | 0.0036 | 0.0001 | 12115 | Ovarian cancer |
| GSE57477 | BN | 0.1082 | 0.0019 | 0.0034 | 0.0002 | 11957 | Ovarian cancer |
| GSE57477 | RTF | 0.1126 | 0.004 | 0.0037 | 0.0003 | 10931 | Ovarian cancer |
| GSE54388 | GMM | 0.1175 | 0.0091 | 0.003 | 0.0007 | 18473 | Ovarian cancer |
| GSE54388 | CTGAN | 0.1378 | 0.0047 | 0.0055 | 0.002 | 14954 | Ovarian cancer |
| GSE54388 | BN | 0.1367 | 0.003 | 0.0051 | 0.0006 | 15785 | Ovarian cancer |
| GSE54388 | RTF | 0.134 | 0.0103 | 0.0044 | 0.0012 | 14560 | Ovarian cancer |
| GSE25427 | GMM | 0.0909 | 0.0054 | 0.0022 | 0.0003 | 16910 | Ovarian cancer |
| GSE25427 | CTGAN | 0.1353 | 0.0041 | 0.0054 | 0.0007 | 10680 | Ovarian cancer |
| GSE25427 | BN | 0.1268 | 0.0037 | 0.004 | 0.0002 | 11397 | Ovarian cancer |
| GSE25427 | RTF | 0.1423 | 0.0048 | 0.0057 | 0.0005 | 9665 | Ovarian cancer |
| GSE14407 | GMM | 0.1057 | 0.0076 | 0.0025 | 0.0005 | 20179 | Ovarian cancer |
| GSE14407 | CTGAN | 0.1399 | 0.0061 | 0.0043 | 0.0004 | 14169 | Ovarian cancer |
| GSE14407 | BN | 0.1397 | 0.0063 | 0.0043 | 0.0002 | 14557 | Ovarian cancer |
| GSE14407 | RTF | 0.14 | 0.0084 | 0.005 | 0.0008 | 14401 | Ovarian cancer |
| GSE38666 | GMM | 0.0905 | 0.003 | 0.0019 | 0.0001 | 15270 | Ovarian cancer |
| GSE38666 | CTGAN | 0.128 | 0.0033 | 0.0044 | 0.0005 | 9715 | Ovarian cancer |
| GSE38666 | BN | 0.1184 | 0.0029 | 0.0036 | 0.0003 | 10446 | Ovarian cancer |
| GSE38666 | RTF | 0.1283 | 0.0025 | 0.0044 | 0.0004 | 9370 | Ovarian cancer |
| GSE29450 | GMM | 0.0961 | 0.0105 | 0.0029 | 0.0013 | 22681 | Ovarian cancer |
| GSE29450 | CTGAN | 0.1444 | 0.0072 | 0.0045 | 0.0006 | 15619 | Ovarian cancer |
| GSE29450 | BN | 0.1461 | 0.0101 | 0.005 | 0.0007 | 15095 | Ovarian cancer |
| GSE29450 | RTF | 0.1621 | 0.015 | 0.0059 | 0.0009 | 13056 | Ovarian cancer |
| GSE235641 | GMM | 0.1469 | 0.0236 | 0.0063 | 0.0024 | 33698 | Ovarian cancer |
| GSE235641 | CTGAN | 0.1648 | 0.0124 | 0.0089 | 0.0035 | 30185 | Ovarian cancer |
| GSE235641 | BN | 0.171 | 0.0173 | 0.0063 | 0.0018 | 30382 | Ovarian cancer |
| GSE235641 | RTF | 0.1501 | 0.0161 | 0.0069 | 0.0045 | 31313 | Ovarian cancer |
| GSE223426 | BN | 0.3701 | 0.0338 | 0.0341 | 0.0133 | 30807 | Ovarian cancer |
| GSE223426 | CTGAN | 0.9313 | 0.0473 | 0.1636 | 0.027 | 14809 | Ovarian cancer |
| GSE223426 | GMM | 0.3112 | 0.0475 | 0.0266 | 0.009 | 30807 | Ovarian cancer |
| GSE223426 | RTF | 0.5062 | 0.0463 | 0.0497 | 0.0127 | 22960 | Ovarian cancer |
| GSE160626 | BN | 0.3719 | 0.0267 | 0.0324 | 0.0125 | 31189 | Ovarian cancer |
| GSE160626 | CTGAN | 0.8342 | 0.0342 | 0.1574 | 0.0168 | 16231 | Ovarian cancer |
| GSE160626 | GMM | 0.3643 | 0.0471 | 0.0277 | 0.0079 | 31179 | Ovarian cancer |
| GSE160626 | RTF | 0.5725 | 0.0561 | 0.0578 | 0.0207 | 23890 | Ovarian cancer |
| GSE143897 | BN | 0.2535 | 0.0145 | 0.0125 | 0.002 | 26453 | Ovarian cancer |
| GSE143897 | CTGAN | 0.9619 | 0.0185 | 0.1831 | 0.0135 | 6652 | Ovarian cancer |
| GSE143897 | GMM | 0.2313 | 0.0058 | 0.0116 | 0.005 | 26460 | Ovarian cancer |
| GSE143897 | RTF | 0.5001 | 0.018 | 0.0351 | 0.0055 | 17023 | Ovarian cancer |
| GSE102094 | BN | 0.2515 | 0.0127 | 0.0089 | 0.0022 | 44219 | Ovarian cancer |
| GSE102094 | CTGAN | 0.7392 | 0.0249 | 0.131 | 0.015 | 12794 | Ovarian cancer |
| GSE102094 | GMM | 0.2368 | 0.0229 | 0.0082 | 0.0016 | 44219 | Ovarian cancer |
| GSE102094 | RTF | 0.4014 | 0.0241 | 0.0149 | 0.0015 | 38358 | Ovarian cancer |
| GSE32646 | GMM | 0.0946 | 0.0008 | 0.0026 | 0.00E+00 | 6931 | Breast cancer |
| GSE32646 | CTGAN | 0.1014 | 0.0008 | 0.003 | 1.00E-04 | 6224 | Breast cancer |
| GSE32646 | BN | 0.1012 | 0.0009 | 0.003 | 1.00E-04 | 6292 | Breast cancer |
| GSE32646 | RTF | 0.1021 | 0.0007 | 0.003 | 1.00E-04 | 6161 | Breast cancer |
| GSE32531 | GMM | 0.1209 | 0.0034 | 0.004 | 0.0004 | 12222 | Breast cancer |
| GSE32531 | CTGAN | 0.1472 | 0.0039 | 0.006 | 0.0004 | 9779 | Breast cancer |
| GSE32531 | BN | 0.1411 | 0.004 | 0.0057 | 0.0008 | 10075 | Breast cancer |
| GSE32531 | RTF | 0.1451 | 0.0032 | 0.0057 | 0.0005 | 9395 | Breast cancer |
| GSE74667 | GMM | 0.0882 | 0.001 | 0.0024 | 1.00E-04 | 6962 | Breast cancer |
| GSE74667 | CTGAN | 0.0958 | 0.0014 | 0.0027 | 1.00E-04 | 6189 | Breast cancer |
| GSE74667 | BN | 0.0962 | 0.0007 | 0.0029 | 0.0001 | 6134 | Breast cancer |
| GSE74667 | RTF | 0.0981 | 0.0017 | 0.0029 | 0.0002 | 6320 | Breast cancer |
| GSE60785 | GMM | 0.1065 | 0.0011 | 0.0033 | 0.0001 | 13776 | Breast cancer |
| GSE60785 | CTGAN | 0.1208 | 0.0011 | 0.0042 | 1.00E-04 | 11604 | Breast cancer |
| GSE60785 | BN | 0.121 | 0.0008 | 0.0043 | 1.00E-04 | 11388 | Breast cancer |
| GSE60785 | RTF | 0.1238 | 0.0015 | 0.0044 | 0.0002 | 11272 | Breast cancer |
| GSE23640 | GMM | 0.1235 | 0.0141 | 0.0049 | 0.0015 | 21504 | Breast cancer |
| GSE23640 | CTGAN | 0.1887 | 0.0152 | 0.0082 | 0.0008 | 15205 | Breast cancer |
| GSE23640 | BN | 0.1796 | 0.0089 | 0.0087 | 0.0011 | 14863 | Breast cancer |
| GSE23640 | RTF | 0.1868 | 0.0092 | 0.009 | 0.0017 | 14455 | Breast cancer |
| GSE43358 | GMM | 0.1035 | 0.0011 | 0.0035 | 0.0002 | 10078 | Breast cancer |
| GSE43358 | CTGAN | 0.1287 | 0.0022 | 0.0059 | 0.0003 | 8062 | Breast cancer |
| GSE43358 | BN | 0.1306 | 0.0018 | 0.0056 | 0.0002 | 7756 | Breast cancer |
| GSE43358 | RTF | 0.1304 | 0.002 | 0.0057 | 0.0003 | 7817 | Breast cancer |
| GSE18728 | GMM | 0.1946 | 0.0034 | 0.0125 | 0.0006 | 5433 | Breast cancer |
| GSE18728 | CTGAN | 0.2113 | 0.007 | 0.0146 | 0.0009 | 5070 | Breast cancer |
| GSE18728 | BN | 0.211 | 0.0045 | 0.0143 | 0.0008 | 5138 | Breast cancer |
| GSE18728 | RTF | 0.1784 | 0.0037 | 0.011 | 0.0004 | 4680 | Breast cancer |
| GSE58135 | BN | 0.2737 | 0.009 | 0.0107 | 0.0015 | 41888 | Breast cancer |
| GSE58135 | CTGAN | 0.7586 | 0.0183 | 0.102 | 0.0039 | 17261 | Breast cancer |
| GSE58135 | GMM | 0.2535 | 0.0122 | 0.0106 | 0.0019 | 41888 | Breast cancer |
| GSE58135 | RTF | 0.4933 | 0.0222 | 0.0318 | 0.0043 | 30138 | Breast cancer |
| GSE52194 | BN | 0.3701 | 0.0338 | 0.0341 | 0.0133 | 30807 | Breast cancer |
| GSE52194 | CTGAN | 0.9313 | 0.0473 | 0.1636 | 0.027 | 14809 | Breast cancer |
| GSE52194 | GMM | 0.3112 | 0.0475 | 0.0266 | 0.009 | 30807 | Breast cancer |
| GSE52194 | RTF | 0.5062 | 0.0463 | 0.0497 | 0.0127 | 22960 | Breast cancer |

## Bibliography

1. Golub, T. R. et al. Molecular classification of cancer: class discovery and class prediction by gene expression monitoring. Science 286, 531–537 (1999). 10.1126/science.286.5439.531

2. Barrett, T. et al. NCBI GEO: archive for functional genomics data sets-update. Nucleic Acids Res. 41, D991–D995 (2013). 10.1093/nar/gks1193

3. Clarke, R. et al. The properties of high-dimensional data spaces: implications for exploring gene and protein expression data. Nat. Rev. Cancer 8, 37–49 (2008). 10.1038/nrc2294

4. Button, K. S. et al. Power failure: why small sample size undermines the reliability of neuroscience. Nat. Rev. Neurosci. 14, 365–376 (2013). 10.1038/nrn3475

5. Abayomi-Alli, O. O., Damaševičius, R., Wieczorek, M. & Woźniak, M. Data Augmentation Using Principal Component Resampling for Image Recognition by Deep Learning. In Artificial Intelligence and Soft Computing – ICAISC 2019 (2020). 10.1007/978-3-030-61534-5_4

6. Chawla, N. V., Bowyer, K. W., Hall, L. O. & Kegelmeyer, W. P. SMOTE: synthetic minority over-sampling technique. Preprint at https://arxiv.org/abs/1106.1813 (2011).

7. Kaur, D. et al. Application of Bayesian networks to generate synthetic health data. J. Am. Med. Inform. Assoc. 28, 801–811 (2021). 10.1093/jamia/ocaa303

8. Lèbre, S., Becq, J., Devaux, F., Stumpf, M. P. & Lelandais, G. Statistical inference of the time-varying structure of gene-regulation networks. BMC Syst. Biol. 4, 130 (2010). 10.1186/1752-0509-4-130

9. Goodfellow, I. J., et al. Generative adversarial networks. Preprint at https://arxiv.org/abs/1406.2661 (2014).

10. Ho, J., Jain, A. & Abbeel, P. Denoising Diffusion Probabilistic Models. Preprint at https://arxiv.org/abs/2006.11239 (2020).

11. Kingma, D. P. & Welling, M. Auto-Encoding Variational Bayes. Preprint at https://arxiv.org/abs/1312.6114 (2013).

12. Beaulieu-Jones, B. K. et al. Privacy-preserving generative deep neural networks support clinical data sharing. Circ. Cardiovasc. Qual. Outcomes 12, e005122 (2019). 10.1161/CIRCOUTCOMES.118.005122

13. Ghahramani, A., Watt, F. M. & Luscombe, N. M. Generative adversarial networks simulate gene expression and predict perturbations in single cells. Preprint at 10.1101/262501 (2018).

14. 14. Nußberger, J., Boesel, F., Lenz, S., Binder, H. & Hess, M. Synthetic observations from deep generative models and binary omics data with limited sample size. Preprint at 10.1101/2020.06.11.147058 (2020).

15. Cui, H. et al. scGPT: toward building a foundation model for single-cell multi-omics using generative AI. Nat. Methods 21, 1470–1480 (2024). 10.1038/s41592-024-02201-0

16. Hao, M. et al. Large-scale foundation model on single-cell transcriptomics. Nat. Methods 21, 1481–1491 (2024). 10.1038/s41592-024-02305-7

17. Theodoris, C. V. et al. Transfer learning enables predictions in network biology. Nature 618, 616–624 (2023). 10.1038/s41586-023-06139-9

18. Yang, H., Robinson, P. & Wang, K. Phenolyzer: phenotype-based prioritization of candidate genes for human diseases. Nat. Methods 12, 841–843 (2015). 10.1038/nmeth.3484

19. Rubbi, A., Di Francesco, A. G., Lotfollahi, M. & Liò, P. A Standardized Framework for Evaluating Gene Expression Generative Models. Preprint at https://arxiv.org/abs/2603.11244 (2026).

20. Li, C. C., Chiang, T. C., Wu, T. S., Pacheco-Rodriguez, G., Moss, J. & Lee, F. J. ARL4D recruits cytohesin-2/ARNO to modulate actin remodeling. Mol. Biol. Cell 18, 4420–4437 (2007). 10.1091/mbc.e07-02-0149

21. Su, Z. Z., et al. Cloning and characterization of SARI (suppressor of AP-1, regulated by IFN). Proc. Natl. Acad. Sci. USA 105, 20906–20911 (2008). 10.1073/pnas.0807975106

22. Sun, F., et al. Long noncoding RNA LINC00963 induces NOP2 expression by sponging tumor suppressor miR-542-3p to promote metastasis in prostate cancer. Aging 12, 11500–11516 (2020). 10.18632/aging.103236

23. Karimbayli, J., Pellarin, I., Belletti, B. & Baldassarre, G. Insights into the structural and functional activities of forgotten Kinases: PCTAIRE CDKs. Mol. Cancer 23, 135 (2024). 10.1186/s12943-024-02043-6

24. Ma, Q., Baldwin, K. T., Renzelli, A. J., McDaniel, A. & Dong, L. TCDD-inducible poly(ADP-ribose) polymerase: a novel response to 2,3,7,8-tetrachlorodibenzo-p-dioxin. Biochem. Biophys. Res. Commun. 289, 499–506 (2001). 10.1006/bbrc.2001.5987

25. Shigehara, Y. et al. Mutations in SDR9C7 gene encoding an enzyme for vitamin A metabolism underlie autosomal recessive congenital ichthyosis. Hum. Mol. Genet. 25, 4484–4493 (2016). 10.1093/hmg/ddw277

26. Falender, A. E., Freiman, R. N., Geles, K. G., Lo, K. C., Hwang, K., Lamb, D. J., Morris, P. L., Tjian, R. & Richards, J. S. Maintenance of spermatogenesis requires TAF4b, a gonad-specific subunit of TFIID. Genes Dev. 19, 794–803 (2005). 10.1101/gad.1290105

27. Andrzejewski, S. et al. PGC-1α promotes breast cancer metastasis and confers bioenergetic flexibility against metabolic drugs. Cell Metab. 26, 778–787.e5 (2017). 10.1016/j.cmet.2017.09.006

28. Zhang, S. et al. Targeting TAF1 with BAY-299 induces antitumor immunity in triple-negative breast cancer. Biochem. Biophys. Res. Commun. 665, 55–63 (2023). 10.1016/j.bbrc.2023.04.100

29. Apte, R. S., Chen, D. S. & Ferrara, N. VEGF in Signaling and Disease: Beyond Discovery and Development. Cell 176, 1248–1264 (2019). 10.1016/j.cell.2019.01.021

30. Finn, R. S. et al. PD 0332991, a selective cyclin D kinase 4/6 inhibitor, preferentially inhibits proliferation of luminal estrogen receptor-positive human breast cancer cell lines in vitro. Breast Cancer Res. 11, R77 (2009). 10.1186/bcr2419

31. Ray, P. S. et al. FOXC1 is a potential prognostic biomarker with functional significance in basal-like breast cancer. Cancer Res. 70, 3870–3876 (2010).

32. Jubb, A. M. et al. Expression of vascular notch ligand delta-like 4 and inflammatory markers in breast cancer. Am. J. Pathol. 176, 2019–2028 (2010). 10.2353/ajpath.2010.090908

33. Ye, L. et al. Overexpression of CDCA7 predicts poor prognosis and induces EZH2-mediated progression of triple-negative breast cancer. Int. J. Cancer 143, 2602–2613 (2018). 10.1002/ijc.31766

34. Christensen, E. M., Patel, S. M., Korasick, D. A., Campbell, A. C., Krause, K. L., Becker, D. F. & Tanner, J. J. Resolving the cofactor-binding site in the proline biosynthetic enzyme human Δ1-pyrroline-5-carboxylate reductase 1. J. Biol. Chem. 292, 7233–7243 (2017). 10.1074/jbc.M117.780288

35. Ou, Y., Wang, S. J., Li, D., Chu, B. & Gu, W. Activation of SAT1 engages polyamine metabolism with p53-mediated ferroptotic responses. Proc. Natl. Acad. Sci. USA 113, E6806–E6812 (2016). 10.1073/pnas.1607152113

36. Sanderson, S. M., Gao, X., Dai, Z. & Locasale, J. W. Methionine metabolism in health and cancer: a nexus of diet and precision medicine. Nat. Rev. Cancer 19, 625–637 (2019). 10.1038/s41568-019-0187-8

37. Wilson, T. J., Gilfillan, S. & Colonna, M. Fc Receptor-Like A Associates with Intracellular IgG and IgM but Is Dispensable for Antigen-Specific Immune Responses. J. Immunol. 185, 2960–2967 (2010). 10.4049/jimmunol.1001428

38. Wu, S. M., Cheung, W. F., Frazier, D. & Stafford, D. W. Cloning and expression of the cDNA for human γ-glutamyl carboxylase. Science 254, 1634–1636 (1991). 10.1126/science.1749935

39. Monk, B. J., Choi, D. C., Pugmire, G. & Burger, R. A. Activity of bevacizumab (rhuMAB VEGF) in advanced refractory epithelial ovarian cancer. Gynecol. Oncol. 96, 902–905 (2005). 10.1016/j.ygyno.2004.12.001

40. Han, J. et al. CENPA is one of the potential key genes associated with the proliferation and prognosis of ovarian cancer based on integrated bioinformatics analysis and regulated by MYBL2. Transl. Cancer Res. 10, 4076–4086 (2021). 10.21037/tcr-21-175

41. Fazili, Z., Sun, W., Mittelstaedt, S., Cohen, C. & Xu, X. X. Disabled-2 inactivation is an early step in ovarian tumorigenicity. Oncogene 18, 3104–3113 (1999). 10.1038/sj.onc.1202649

42. Nam, S. et al. Control of Rapsyn Stability by the CUL-3-containing E3 Ligase Complex. J. Biol. Chem. 284, 8195–8206 (2009). 10.1074/jbc.M808230200

43. Lester, L. B., Coghlan, V. M., Nauert, B. & Scott, J. D. Cloning and Characterization of a Novel A-kinase Anchoring Protein: AKAP 220, Association with Testicular Peroxisomes. J. Biol. Chem. 271, 9460–9465 (1996). 10.1074/jbc.271.16.9460

44. Römisch, J., Vermöhlen, S., Feussner, A. & Stöhr, H. A. The FVII Activating Protease Cleaves Single-Chain Plasminogen Activators. Pathophysiol. Haemost. Thromb. 29, 292–299 (1999). 10.1159/000022515

45. Zhou, Y. et al. TTD: Therapeutic Target Database describing target druggability information. Nucleic Acids Res. 52, D1465–D1477 (2024). 10.1093/nar/gkad751

46. Fonseca-Correa, J. I. & Correa-Rotter, R. Sodium-Glucose Cotransporter 2 Inhibitors Mechanisms of Action: A Review. Front. Med. 8, 777861 (2021). 10.3389/fmed.2021.777861

47. Victoriano, M. et al. From virtual experiments to biomedical insight with synthetic data. Nat. Mach. Intell. 8, 866–879 (2026). 10.1038/s42256-026-01244-6

48. Tosh, K., et al. Variants in the SP110 gene are associated with genetic susceptibility to tuberculosis in West Africa. Proc. Natl. Acad. Sci. USA 103, 10364–10368 (2006). 10.1073/pnas.0603340103

49. Fedele, M., Crescenzi, E. & Cerchia, L. The POZ/BTB and AT-Hook Containing Zinc Finger 1 (PATZ1) Transcription Regulator: Physiological Functions and Disease Involvement. Int. J. Mol. Sci. 18, 2524 (2017). 10.3390/ijms18122524

50. Maroněk, M. & Gardlík, R. The Citrullination-Neutrophil Extracellular Trap Axis in Chronic Diseases. J. Innate Immun. 14, 393–417 (2022). 10.1159/000522331

51. Zhong, Z. et al. New mitochondrial DNA synthesis enables NLRP3 inflammasome activation. Nature 560, 198–203 (2018). 10.1038/s41586-018-0372-z

52. Mendonca, L. E. et al. Human alveolar macrophage metabolism is compromised during Mycobacterium tuberculosis infection. Front. Immunol. 13, 1044592 (2022). 10.3389/fimmu.2022.1044592

53. Guo, L. et al. NUAK1 Promotes Diabetic Kidney Disease by Accelerating Renal Tubular Senescence via the ROS/P53 Axis. Diabetes 74, 2405–2417 (2025). 10.2337/db25-0417

54. Lohman, D. C., et al. Mitochondrial COQ9 is a lipid-binding protein that associates with COQ7 to enable coenzyme Q biosynthesis. Proc. Natl. Acad. Sci. USA 111, E4697–E4705 (2014). 10.1073/pnas.1413128111

55. Farook, M. R. et al. Loss of mitochondrial pyruvate carrier 1 supports proline-dependent proliferation and collagen biosynthesis in ovarian cancer. Mol. Metab. 81, 101900 (2024). 10.1016/j.molmet.2024.101900

56. Wang, M. H., Ronsin, C., Gesnel, M. C., Coupey, L., Skeel, A., Leonard, E. J. & Breathnach, R. Identification of the ron Gene Product as the Receptor for the Human Macrophage Stimulating Protein. Science 266, 117–119 (1994). 10.1126/science.7939629

57. Brckalo, T., Calzetti, F., Pérez-Cabezas, B., Borràs, F. E., Cassatella, M. A. & López-Botet, M. Functional analysis of the CD300e receptor in human monocytes and myeloid dendritic cells. Eur. J. Immunol. 40, 722–732 (2010). 10.1002/eji.200939468

58. Ohoka, N., Yoshii, S., Hattori, T., Onozaki, K. & Hayashi, H. TRB3, a novel ER stress-inducible gene, is induced via ATF4-CHOP pathway and is involved in cell death. EMBO J. 24, 1243–1255 (2005). 10.1038/sj.emboj.7600596

59. Zhang, Y. M., Rock, C. O. & Jackowski, S. Feedback regulation of murine pantothenate kinase 3 by coenzyme A and coenzyme A thioesters. J. Biol. Chem. 280, 32594–32601 (2005). 10.1074/jbc.M506275200

60. Brown, E. J. et al. Mutations in the formin gene INF2 cause focal segmental glomerulosclerosis. Nat. Genet. 42, 72–76 (2010). 10.1038/ng.505

61. Gottweis, J. et al. Accelerating scientific discovery with Co-Scientist. Nature 655, 487–496 (2026). 10.1038/s41586-026-10644-y

62. Chmelařová, M. et al. Methylation analysis of tumour suppressor genes in ovarian cancer using MS-MLPA. Folia Biol. (Praha) 58, 246–250 (2012).

63. Montavon, C. et al. Prognostic and diagnostic significance of DNA methylation patterns in high grade serous ovarian cancer. Gynecol. Oncol. 124, 582–588 (2012).

64. Lachmann, A. et al. Massive mining of publicly available RNA-seq data from human and mouse. Nat. Commun. 9, 1366 (2018). 10.1038/s41467-018-03751-6

65. Robinson, M. D., McCarthy, D. J. & Smyth, G. K. edgeR: a Bioconductor package for differential expression analysis of digital gene expression data. Bioinformatics 26, 139–140 (2010). 10.1093/bioinformatics/btp616

66. Ritchie, M. E. et al. limma powers differential expression analyses for RNA-sequencing and microarray studies. Nucleic Acids Res. 43, e47 (2015). 10.1093/nar/gkv007

67. Law, C. W., Chen, Y., Shi, W. & Smyth, G. K. voom: precision weights unlock linear model analysis tools for RNA-seq read counts. Genome Biol. 15, R29 (2014). 10.1186/gb-2014-15-2-r29

68. Xu, L., Skoularidou, M., Cuesta-Infante, A. & Veeramachaneni, K. Modeling tabular data using conditional GAN. Adv. Neural Inf. Process. Syst. 32 (2019). 10.48550/arXiv.1907.00503

69. Scutari, M. Learning Bayesian networks with the bnlearn R package. J. Stat. Softw. 35, 1–22 (2010). 10.18637/jss.v035.i03

70. Solatorio, A. V. & Dupriez, O. REaLTabFormer: generating realistic relational and tabular data using transformers. Preprint at arXiv 10.48550/arXiv.2302.02041 (2023).

71. Kolberg, L. et al. g:Profiler-interoperable web service for functional enrichment analysis and gene identifier mapping (2023 update). Nucleic Acids Res. 51, W207–W212 (2023). 10.1093/nar/gkad347

72. Cerami, E. et al. The cBio cancer genomics portal: an open platform for exploring multidimensional cancer genomics data. Cancer Discov. 2, 401–404 (2012). 10.1158/2159-8290.CD-12-0095

## Supplementary References

1. Hotchkiss, R. S., Monneret, G. & Payen, D. Sepsis-induced immunosuppression: from cellular dysfunctions to immunotherapy. Nat. Rev. Immunol. 13, 862–874 (2013). 10.1038/nri3552

2. Fedele, M., Crescenzi, E. & Cerchia, L. The POZ/BTB and AT-Hook Containing Zinc Finger 1 (PATZ1) Transcription Regulator: Physiological Functions and Disease Involvement. Int. J. Mol. Sci. 18, 2524 (2017). 10.3390/ijms18122524

3. Maroněk, M. & Gardlík, R. The Citrullination-Neutrophil Extracellular Trap Axis in Chronic Diseases. J. Innate Immun. 14, 393–417 (2022). 10.1159/000522331

4. Nakaya, M. et al. Inflammatory T cell responses rely on amino acid transporter ASCT2 facilitation of glutamine uptake and mTORC1 kinase activation. Immunity 40, 692–705 (2014). 10.1016/j.immuni.2014.04.007

5. Ohoka, N., Yoshii, S., Hattori, T., Onozaki, K. & Hayashi, H. TRB3, a novel ER stress-inducible gene, is induced via ATF4-CHOP pathway and is involved in cell death. EMBO J. 24, 1243–1255 (2005). 10.1038/sj.emboj.7600596

6. Downs, I., Vijayan, S., Sidiq, T. & Kobayashi, K. S. CITA/NLRC5: a critical transcriptional regulator of MHC class I gene expression. BioFactors 42, 349–357 (2016). 10.1002/biof.1285

7. Ma, W. et al. Exploring the relationship between sepsis and Golgi apparatus dysfunction: bioinformatics insights and diagnostic marker discovery. Front. Genet. 16, 1483493 (2025). 10.3389/fgene.2025.1483493

8. Zhu, D., Zhu, K. & Guo, S. Identification of key genes related to immune cells in patients with gram-negative sepsis based on weighted gene co-expression network analysis. Ann. Transl. Med. 10, 787–787 (2022). 10.21037/atm-22-3307

9. Tattevin, P. et al. Enhanced indoleamine 2,3-dioxygenase activity in patients with severe sepsis and septic shock. J. Infect. Dis. 201, 956–966 (2010). 10.1086/650996

10. Saelee, C. et al. Toll-like receptor-mediated innate immune responses by recognition of the recombinant dormancy-associated Mycobacterium tuberculosis proteins Rv2659c and Rv1738. PLoS ONE 17, e0273517 (2022). 10.1371/journal.pone.0273517

11. Zihad, S. M. N. K. et al. Role of pattern recognition receptors in sensing Mycobacterium tuberculosis. Heliyon 9, e20636 (2023). 10.1016/j.heliyon.2023.e20636

12. Shariq, M. et al. The exploitation of host autophagy and ubiquitin machinery by Mycobacterium tuberculosis in shaping immune responses and host defense during infection. Autophagy 19, 3–23 (2023). 10.1080/15548627.2021.2021495

13. Song, C. H. et al. Role of mitogen-activated protein kinase pathways in the production of tumor necrosis factor-α, interleukin-10, and monocyte chemotactic protein-1 by Mycobacterium tuberculosis H37Rv-infected human monocytes. J. Clin. Immunol. 23, 194–201 (2003). 10.1023/A:1023309928879

14. Lin, P. L., Plessner, H. L., Voitenok, N. N. & Flynn, J. L. Tumor necrosis factor and tuberculosis. J. Investig. Dermatol. Symp. Proc. 12, 22–25 (2007). 10.1038/sj.jidsymp.5650027

15. Zhong, Z. et al. New mitochondrial DNA synthesis enables NLRP3 inflammasome activation. Nature 560, 198–203 (2018). 10.1038/s41586-018-0372-z

16. Ma, J. et al. The roles of inflammasomes in host defense against Mycobacterium tuberculosis. Pathogens 10, 120 (2021). 10.3390/pathogens10020120

17. Cheng, Y. & Schorey, J. S. Mycobacterium tuberculosis-induced IFN-β production requires cytosolic DNA and RNA sensing pathways. J. Exp. Med. 215, 2919–2935 (2018). 10.1084/jem.20180508

18. Mendonca, L. E. et al. Human alveolar macrophage metabolism is compromised during Mycobacterium tuberculosis infection. Front. Immunol. 13, 1044592 (2022). 10.3389/fimmu.2022.1044592

19. Liebermann, D. A. & Hoffman, B. Gadd45 in the response of hematopoietic cells to genotoxic stress. Blood Cells Mol. Dis. 39, 329–335 (2007). 10.1016/j.bcmd.2007.06.006

20. Wang, M. H., Ronsin, C., Gesnel, M. C., Coupey, L., Skeel, A., Leonard, E. J. & Breathnach, R. Identification of the ron Gene Product as the Receptor for the Human Macrophage Stimulating Protein. Science 266, 117–119 (1994). 10.1126/science.7939629

21. Brckalo, T., Calzetti, F., Pérez-Cabezas, B., Borràs, F. E., Cassatella, M. A. & López-Botet, M. Functional analysis of the CD300e receptor in human monocytes and myeloid dendritic cells. Eur. J. Immunol. 40, 722–732 (2010). 10.1002/eji.200939468

22. Hartman, M. L. & Czyz, M. BCL-w: apoptotic and non-apoptotic role in health and disease. Cell Death Dis. 11, 260 (2020). 10.1038/s41419-020-2417-0

95. Tosh, K., et al. Variants in the SP110 gene are associated with genetic susceptibility to tuberculosis in West Africa. Proc. Natl. Acad. Sci. USA 103, 10364–10368 (2006). 10.1073/pnas.0603340103

24. Kjer-Nielsen, L. et al. MR1 presents microbial vitamin B metabolites to MAIT cells. Nature 491, 717–723 (2012). 10.1038/nature11605

25. Guo, L. et al. NUAK1 Promotes Diabetic Kidney Disease by Accelerating Renal Tubular Senescence via the ROS/P53 Axis. Diabetes 74, 2405–2417 (2025). 10.2337/db25-0417

98. Lohman, D. C., et al. Mitochondrial COQ9 is a lipid-binding protein that associates with COQ7 to enable coenzyme Q biosynthesis. Proc. Natl. Acad. Sci. USA 111, E4697–E4705 (2014). 10.1073/pnas.1413128111

27. Jia, Z. et al. Protein arginine methyltransferase PRMT5 regulates fatty acid metabolism and lipid droplet biogenesis in white adipose tissues. Adv. Sci. 7, 2002602 (2020). 10.1002/advs.202002602

28. Zhang, Y. M., Rock, C. O. & Jackowski, S. Feedback regulation of murine pantothenate kinase 3 by coenzyme A and coenzyme A thioesters. J. Biol. Chem. 280, 32594–32601 (2005). 10.1074/jbc.M506275200

29. Sharma, L. K. et al. A high-throughput screen reveals new small-molecule activators and inhibitors of pantothenate kinases. J. Med. Chem. 58, 1563–1568 (2015). 10.1021/jm501585q

30. Korobova, F., Ramabhadran, V. & Higgs, H. N. An actin-dependent step in mitochondrial fission mediated by the ER-associated formin INF2. Science 339, 464–467 (2013). 10.1126/science.1228360

31. Chakrabarti, R. et al. INF2-mediated actin polymerization at the ER stimulates mitochondrial calcium uptake, inner membrane constriction, and division. J. Cell Biol. 217, 251–268 (2018). 10.1083/jcb.201709111

32. Brown, E. J. et al. Mutations in the formin gene INF2 cause focal segmental glomerulosclerosis. Nat. Genet. 42, 72–76 (2010). 10.1038/ng.505

33. Fonseca-Correa, J. I. & Correa-Rotter, R. Sodium-Glucose Cotransporter 2 Inhibitors Mechanisms of Action: A Review. Front. Med. 8, 777861 (2021). 10.3389/fmed.2021.777861

34. Chakraborty, S. et al. Role of Tmem163 in zinc-regulated insulin storage of MIN6 cells: functional exploration of an Indian type 2 diabetes GWAS associated gene. Biochem. Biophys. Res. Commun. 522, 1022–1029 (2020). 10.1016/j.bbrc.2019.11.117

